# Host-aware Identification of Intrinsic Gene Expression Biopart Parameters from Combinatorial Libraries

**DOI:** 10.64898/2026.01.21.700808

**Authors:** Jesús Picó, Andrés Arboleda-García, David R. Penas, Julio R. Banga, Alejandro Vignoni, Yadira Boada

## Abstract

Model-based design in synthetic biology is limited by the lack of quantitative, mechanistically interpretable biopart parameters that remain valid across genetic and physiological contexts. This limitation is particularly acute for transcriptional units, whose expression phenotypes emerge from nonlinear coupling between plasmid copy number, transcription, translation, and host resource allocation. Here we introduce a context- and host-aware framework for the absolute characterisation of gene expression bioparts embedded in combinatorial libraries of constitutive transcriptional units. Our approach leverages a digital twin of *Escherichia coli* conditioned on experimentally measured growth rate, used as a low-dimensional physiological proxy for cellular state. By embedding this growth-conditioned digital twin in a model-in-the-loop identification strategy, host–circuit interactions are explicitly accounted for and decoupled from intrinsic biopart properties.

Using structured combinatorial libraries, we identify biophysically interpretable and transferable parameters for plasmid origins, promoters, and ribosome binding sites. In particular, we uncover an *intrinsic translation initiation capacity* of ribosome binding sites that remains invariant across genetic contexts and growth conditions, while context-dependent translation rates emerge as physiological projections of this invariant descriptor. This intrinsic parameterisation enables accurate prediction of protein synthesis across diverse host states, supports incremental and patchwork library expansion, and reveals localized failures of modularity that are obscured by phenotype-only characterisation.

Together, these results establish a principled link between DNA sequence, intrinsic biopart parameters, and circuit-level phenotypes, providing a scalable and host-aware foundation for predictive design in synthetic biology.

## 1 Introduction

Synthetic Biology aspires to become an engineering discipline: the rational and systematic design of complex biological systems with novel functions, guided by principles and tools originating from engineering. In this context, the Design–Build–Test–Learn (DBTL) cycle has emerged as the dominant paradigm for the development of synthetic biological systems, and recent years have seen substantial efforts to integrate mathematical modelling, computational optimisation, dynamic regulation, automation, and data-driven analysis into unified DBTL pipelines [1–3]. Despite this progress, significant challenges remain across all stages of the DBTL cycle. In particular, model-based design—a cornerstone of engineering practice—is still severely constrained by the gap between how biological systems are represented in mathematical models and the type of quantitative, mechanistically interpretable information required in the wet lab to reliably construct genetic circuits with prescribed performance.

At present, this gap prevents the full predictive power of mathematical models from being exploited for the rational design of genetic circuits. The challenges associated with modelling and characterising gene expression systems are multifaceted, requiring an understanding of dynamics that are directly and indirectly coupled through cellular resource allocation [4–6]. These challenges are further compounded by practical limitations in experimental observability, the high nonlinearity and dimensionality of mechanistic models, and the pervasive presence of structural and practical identifiability issues [7–9]. As a result, many existing models achieve good predictive performance only at the cost of losing mechanistic interpretability: model parameters often lack a clear biological meaning and cannot be directly mapped to the DNA sequences of the corresponding biological building blocks (bioparts). Consequently, such models are of limited use for systematic circuit design and typically provide only qualitative guidance for biopart selection or circuit prototyping [10]. There remains, therefore, a strong need for coarse-grained yet mechanistically grounded models that establish an explicit link between phenotype, model parameters, and biopart DNA sequences [11, 12].

Recent advances in synthetic biology modelling have increasingly focused on host–circuit interactions, particularly on the constraints imposed by cellular resource allocation [13–16]. These host-aware models have successfully explained a range of non-intuitive behaviours arising from competition for shared cellular resources, revealing how synthetic circuit function is modulated by the physiological state of the host cell [17–19]. In parallel, multi-scale modelling frameworks have been proposed to connect intracellular dynamics with macroscopic behaviour at the bioprocess level, thereby bridging the gap between circuit design and bioprocess optimisation [20–22]. Despite these advances, host-aware models have not yet been systematically exploited as part of an explicit parameter-identification framework for the experimental characterisation of genetic bioparts with improved predictability and transferability.

Transcriptional units (TUs) constitute one of the fundamental modular elements in synthetic biology and are widely used as building blocks for the construction of increasingly complex genetic circuits [23, 24]. Although conceptually simple, TUs combine three of the most commonly used bioparts in synthetic biology—plasmid origins of replication (ORIs), promoters, and ribosome binding sites (RBSs)—whose quantitative characterisation is essential for model-based circuit design. However, even for such elementary devices, reliable characterisation remains challenging. On the one hand, TU phenotypes, typically quantified through protein synthesis rates, are inherently host-dependent due to competition for shared cellular resources [25, 26]. On the other hand, context dependence manifests not merely as experimental variability, but as systematic confounding between transcriptional, translational, and copy-number effects. Because the synthesis rate depends multiplicatively on plasmid copy number and promoter and RBS strengths, standard experimental measurements and parameter estimation approaches can only recover lumped parameter combinations. In other words, the underlying models are not identifiable. Crucially, this limitation does not stem from a lack of data, but from a lack of mechanistic and experimental structure enabling parameter identifiability.

Most experimental strategies for biopart characterization have focused on restricted library designs, either by varying a single biopart at a time within otherwise fixed constructs [27, 28], or by treating entire expression cassettes as indivisible functional units evaluated under narrowly controlled experimental conditions [29]. These approaches have been instrumental in revealing reproducibility limits and context effects, but they provide only limited leverage to disentangle the individual contributions of plasmid copy number, promoter activity, and translational efficiency.

In parallel, recent genome-wide studies have demonstrated that absolute, context-independent quantification of translational processes is in principle achievable *in vivo*, using high-throughput sequencing approaches applied to native genes [30]. While these studies establish an important biophysical baseline for translation in *E. coli*, they do not address the characterization of synthetic bioparts embedded in heterologous genetic contexts, nor do they provide a parameterized modeling framework that can be directly reused for the modular design and prediction of synthetic transcriptional units.

Overall, existing approaches either lack the combinatorial diversity required to resolve coupled biopart effects, or operate outside the synthetic biology design space of exogenous constructs. As a result, a principled methodology that combines absolute, biophysically interpretable parameterization with combinatorial experimental design and explicit host–circuit coupling remains missing.

In this study, we propose a context- and host-aware methodology for the model-based characterisation of bioparts embedded within combinatorial libraries of constitutive transcriptional units. Our approach explicitly frames biopart characterisation as a system identification problem, in which parameter identifiability emerges from the joint design of the model structure and the experimental library.

We employ a host-aware digital twin of *Escherichia coli* that integrates mechanistic modelling with real-time physiological measurements, using the experimentally measured specific growth rate as a dynamic proxy for the cellular state. Because growth rate reflects nutrient availability, endogenous metabolism, and the resource burden imposed by synthetic gene expression [5, 31], it provides a compact yet informative observable that constrains host–circuit–environment interactions in a data-informed and mechanistically consistent manner [32, 33].

For transcriptional unit expression, we adopt a modelling framework that explicitly incorporates host–circuit interactions while preserving context-independent parameters associated with individual bioparts. The model accounts for gene copy number, promoter strength, and ribosome binding site properties through a parsimonious parametrization that separates intrinsic part-specific effects from growth-dependent physiological modulation. While cellular context modulates global resource levels and environmental variables, it does not alter the intrinsic values of biopart-specific parameters, enabling an absolute yet context-aware characterisation.

Parameter estimation is performed in a model-in-the-loop fashion: the host digital twin, driven by growth-rate measurements, infers the cellular resource fluxes consistent with the observed physiological state, which are then combined with synthesis-rate measurements from a combinatorial transcriptional unit library to estimate biopart parameters via global optimisation.

Crucially, the combinatorial architecture of the transcriptional unit library ensures that each biopart is observed across multiple, partially overlapping genetic contexts. From an identifiability perspective, this transforms the library into a structured experimental design that provides independent excitation of parameter combinations that would otherwise remain confounded. This strategy not only enables robust and transferable biopart characterisation, but also supports incremental library expansion, in which previously characterised bioparts act as anchors for the identification of new ones. Altogether, our framework establishes a quantitative mapping from biopart DNA sequences to a biophysical parameter space, and from these parameters to circuit-level phenotypes, laying the foundation for scalable, host-aware, model-driven engineering of synthetic gene circuits.

## 2 Results

### 2.1 Experimental regime and growth–burden landscape

We constructed a library of 35 constitutive transcriptional units by combinatorial assembly spanning two plasmid copy numbers, four promoters, and five ribosome binding sites (RBSs), organized into the core library 𝓛_24_, the extended library 𝓛_30_, and the reduced sublibrary 𝓛_5_ (Figure 1A and Methods 3.1). All TUs constitutively expressed the fluorescent protein GFPmut3. An extensive experimental characterization of the entire TU library was performed by measuring absorbance and fluorescence at 5-minute intervals under batch growth conditions (Methods 3.2).

**Fig. 1.**
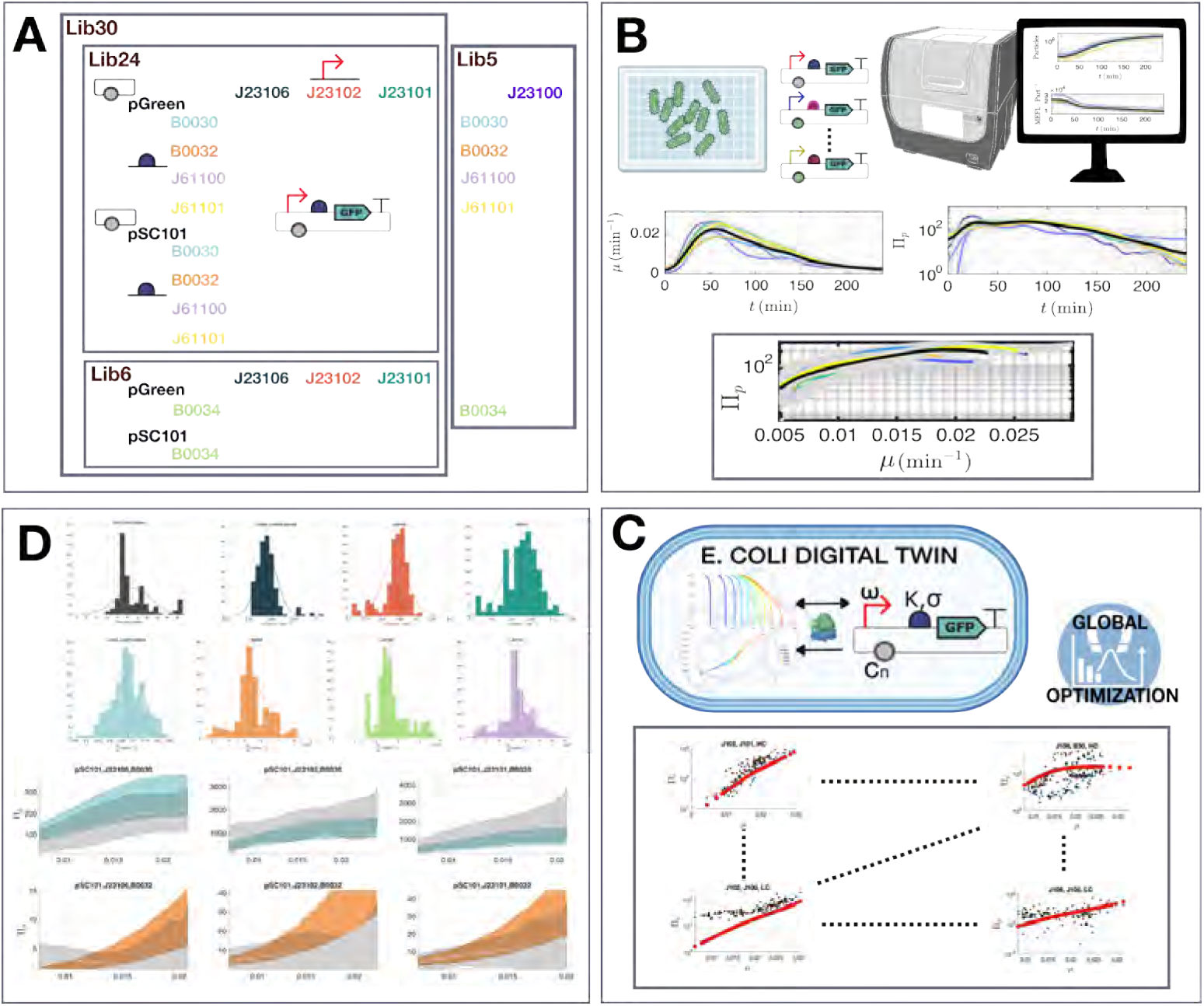
Overview of the combinatorial TU libraries and model-in-the-loop workflow. **A**, Combinatorial design of the constitutive transcriptional-unit (TU) library expressing *gfp* (GFPmut3), assembled from plasmid origins, promoters and RBSs, with each biopart reused across multiple overlapping genetic contexts. The core library *L*_24_ is used as a reference dataset for biopart parameter identification, and the reduced sub-libraries *L*_6_ and *L*_5_ are used for incremental identification and validation of additional parts. **B**, Experimental pipeline and primary observables: time-resolved optical density and fluorescence measurements are converted into cell-specific growth rate *µ*(*t*) and TU synthesis rate Π*_p_*(*t*), yielding TU-specific relationships Π*_p_*(*µ*) across the batch trajectory, with growth rate acting as the primary physiological observable driving inference. **C**, Host-aware *E. coli* digital twin: the experimentally measured growth rate *µ*, rather than unobservable intracellular variables, is used as a proxy to infer a physiologically consistent host state (ribosomal flux and nutrient/elongation regime), which is combined with biopart parameters to predict TU synthesis rates. **D**, Global parameter inference from combinatorial libraries: biopart parameters are estimated by fitting digital-twin predictions to the measured Π*_p_*(*µ*) curves across all TUs, producing intrinsic, context-invariant and identifiable part-specific parameter distributions (via cross-validation) that are then propagated to generate predictive envelopes for Π*_p_*(*µ*) in each genetic context.

To enable quantitative and comparable measurements across constructs and experiments, raw absorbance and fluorescence data were converted into standardized units of Equivalent Particle Count (Particles) and Molecules of Equivalent Fluorescein (MEFL), respectively [11, 34, 35] (Supplementary Information S2.2). Using these standardized signals, we computed time-resolved values of the cell-specific growth rate *µ* and the TU synthesis rate Π*_p_*, expressed in min^−1^ and MEFL · Particle^−1^ · min^−1^, respectively (Figure 1B and Supplementary Information S2.3).

We hypothesized that the relationship between TU synthesis rate Π*_p_* and cell-specific growth rate *µ* captures the characteristic behavior of each TU as a function of the host’s metabolic state. As such, this relationship provides a compact yet informative representation of TU performance across varying physiological conditions and constitutes the primary experimental observable for subsequent model-based analysis (see Section 2.2).

Across the 35 TUs, synthesis rates spanned approximately three orders of magnitude, ranging from 10^0^ to 10^3^ MEFL · Particle^−1^ · min^−1^ (Supplementary Fig. S2.2). When evaluated at the maximum specific growth rate, these values were approximately uniformly distributed across the library (Figure 2A and Supplementary Fig. S2.3), indicating broad coverage of expression strengths within a single experimental framework.

**Fig. 2.**
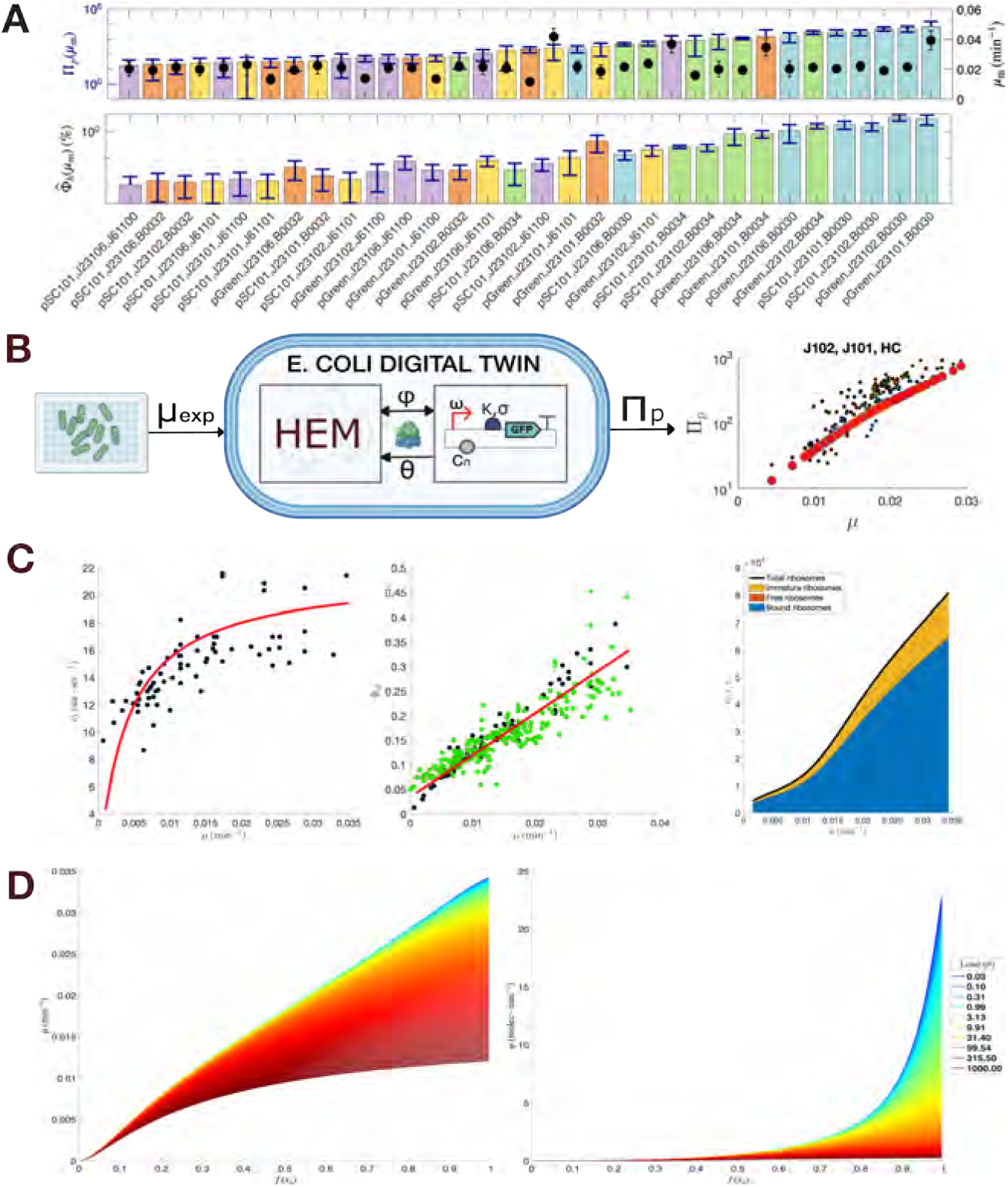
**A, top:** Experimental maximum specific growth rates *µ*_m_ (black dots) and the corresponding TU synthesis rates Π*_p_*(*µ*_m_) (colored bars), ordered by increasing synthesis rate. Values represent means across single-well experiments, with vertical bars showing standard deviations. Colors denote RBS variants. **A, bottom:** Estimated cell burden induced by each TU, expressed as the percentage of host resources captured relative to endogenous gene expression. **B:** Model-in-the-loop framework coupling HEM and TU models for resource-aware synthesis rate prediction. **C:** (Left) Fitted relationship between peptide elongation rate and specific growth rate. (Center) Predicted protein mass fraction of mature active ribosomes (red) compared with values from [4, 38]. (Right) Predicted numbers of bound, free, and immature ribosomes; the black line denotes total ribosomes. **D:** (Left) Predicted growth rate as a function of substrate availability *f* (*s*) under varying exogenous loads. (Right) Corresponding flux of free ribosomes.

The average maximum specific growth rates across TUs ranged from 0.01 to 0.03 min^−1^, corresponding to cell doubling times between 23 and 69 min. These growth rates indicate that TU expression imposed a limited, yet non-negligible, metabolic burden on host cells under the explored conditions. To quantify this effect, we estimated the burden as the fraction of cellular translational resources sequestered by the exogenous constructs (Methods 3.2). As shown in Figure 2A (bottom), the inferred fractional burden remained below 3.5% for all variants (median ≈ 1%; Supplementary Fig. S2.3b and Supplementary Table S2), consistent with the observed growth rates and indicative of a regime in which host–circuit interactions are present and measurable, but not dominated by strong saturation effects. In this regime, synthesis rates remain sensitive to the underlying biopart parameters, providing favourable conditions for subsequent model-based identification. Furthermore, synthesis rates increased monotonically with *µ* across the explored range (Supplementary Fig. S2.2), in agreement with the expected coupling between cellular growth and translational capacity.

Analysis of expression noise revealed distinct scaling behaviors with mean synthesis rate. The Fano factor increased approximately proportionally with Π*_p_*, reaching values above 10^2^ for the most active constructs, while the coefficient of variation decreased from values close to 1 to below 0.2 (Supplementary Fig. S2.4). Thus, although absolute fluctuations increase with expression level, relative noise decreases, consistent with a shift in the dominant contribution to variability as synthesis rates increase, consistent with a transition from promoter-dominated stochasticity to translational resource limitation at high expression regimes [36, 37].

Together, this comprehensive dataset provides a rich, growth-resolved characterization of TU performance and constitutes the experimental foundation for the subsequent model-based, context-aware estimation of biologically meaningful biopart parameters—namely plasmid copy number, promoter strength, and RBS efficiency—presented in the following sections.

### 2.2 Digital-twin-based, context-aware synthesis rate prediction

A central challenge in quantitative biopart characterization is that TU synthesis rates depend not only on intrinsic biopart properties, but also on the physiological state of the host cell. Nutrient availability, endogenous resource allocation, and the translational load imposed by exogenous expression jointly modulate protein synthesis, confounding direct comparisons across genetic constructs and experimental conditions. To address this problem, we developed a *model-in-the-loop* framework based on a digital twin of *E. coli*, which explicitly incorporates host–circuit interactions while remaining anchored to experimentally observable quantities.

The digital twin couples two mechanistic components (Figure 2B): (i) a resource-aware TU synthesis model that describes protein production as a function of ribosomal availability and biopart parameters, and (ii) a Host Equivalent Model (HEM) that captures endogenous host growth and ribosome allocation as functions of nutrient availability and cellular resource fluxes (Methods 3.3–3.4). Importantly, the digital twin is driven by experimentally measured specific growth rates, which act as a compact and informative proxy of the global cellular state, avoiding explicit inference of unobserved substrate levels.

#### Growth rate as a physiological constraint

Rather than attempting to infer unobservable intracellular variables directly, we condition the digital twin on the experimentally measured specific growth rate *µ*. Growth rate integrates the effects of substrate availability, endogenous metabolism, and the translational burden imposed by exogenous gene expression, and therefore provides a strong physiological constraint on host–circuit interactions. By injecting *µ* into the HEM, the digital twin infers a consistent ribosomal resource flux and elongation regime compatible with both host physiology and TU expression, without requiring explicit knowledge of intracellular substrate concentrations or absolute ribosome counts.

#### Resource-aware TU synthesis prediction

Within this framework, the synthesis rate of a protein expressed from a TU is predicted as a function of growth state and intrinsic biopart parameters using the resource-aware TU model (Methods 3.3). This formulation separates *context-dependent variables*—such as ribosomal flux and elongation capacity, inferred by the digital twin—from *biopart-intrinsic parameters* associated with plasmid copy number, promoter strength, and RBS translation properties. As a result, changes in synthesis rate across conditions are attributed to shifts in host state, while the inferred biopart parameters remain invariant.

#### Host Equivalent Model and validation

The HEM predicts endogenous ribosome allocation, growth rate, and ribosomal flux in the absence of exogenous expression, and provides the baseline against which circuit-induced loading is evaluated (Methods 3.4). The HEM recapitulates reported dependencies of ribosomal mass fraction, ribosome numbers, and elongation rates on growth rate (Figure 2C), ensuring that the inferred cellular states are physiologically plausible. When exogenous TUs are present, their translational demand is incorporated as an additional load on the host resource balance, resulting in reduced growth rate and ribosomal flux as the load increases (Figure 2D).

#### Model-in-the-loop decoupling of host, circuit, and environment

By conditioning the digital twin on experimental growth-rate measurements, the model-in-the-loop strategy effectively decouples host, circuit, and environmental effects. For each TU and each observed growth state, the digital twin yields consensus estimates of the ribosomal flux and circuit-induced load that are consistent with both host physiology and TU expression (Methods 3.5). These inferred quantities are then used to predict synthesis rates and to formulate parameter-estimation problems in which the only remaining unknowns are the intrinsic biopart parameters.

This approach collapses high-dimensional and partially unobservable host–circuit–environment interactions onto a one-dimensional physiological observable, enabling robust, context-aware comparison of synthesis-rate data across diverse genetic constructs. As shown below, this digital-twin formulation provides the foundation for global parameter estimation and for exploiting the combinatorial structure of TU libraries to overcome identifiability limitations inherent to isolated construct characterization.

### 2.3 Incremental parameter identification using combinatorial libraries

We approached biopart characterisation as an incremental system identification problem, in which parameter identifiability is progressively improved by expanding the combinatorial diversity of the transcriptional-unit library while reusing previously identified parameters as anchors. Rather than attempting to infer all biopart parameters simultaneously from a single dataset, we followed a staged strategy guided by practical identifiability analysis (Supplementary Information S5).

We first analysed the extended library 𝓛_30_, which spans a broad range of promoters, ribosome binding sites (RBSs), and plasmid copy-number contexts. Parameter estimation using the full host-aware model revealed that the RBS sensitivity parameter *σ*^0^ is only weakly identifiable across the explored growth-rate regime, consistent with the presence of a near-invariant sensitivity direction in the model (Supplementary Information S5). This behaviour reflects the moderate range of growth conditions probed experimentally, which does not sufficiently excite sensitivity-related modes.

On the basis of this analysis, and supported by the physiological regime explored experimentally, we fixed the inverse sensitivity to a biologically plausible value, 1*/σ*^0^ = 0.02, and adopted this reduced translation model for subsequent estimation steps. As shown in Supplementary Information S5, this choice can be interpreted as selecting a representative gauge within an invariant sensitivity subspace, without altering the data-supported estimates of the remaining parameters, while substantially improving numerical conditioning.

Using this reduced model, we next focused on the core library 𝓛_24_, which pro-vides dense combinatorial coverage of promoters, RBSs, and origins of replication under constitutive expression. Global parameter estimation across all transcriptional units and experimental instances yielded consistent and well-constrained estimates of plasmid copy number *N*, promoter transcription rate *ω*, and the intrinsic RBS parameter *κ*^0^ = *K*^0^*/σ*^0^ (Figure 3A; Supplementary Tables S10–S12). Leave-one-out cross-validation confirmed the robustness of these estimates, with narrow predictive intervals and no systematic bias across growth conditions.

**Fig. 3.**
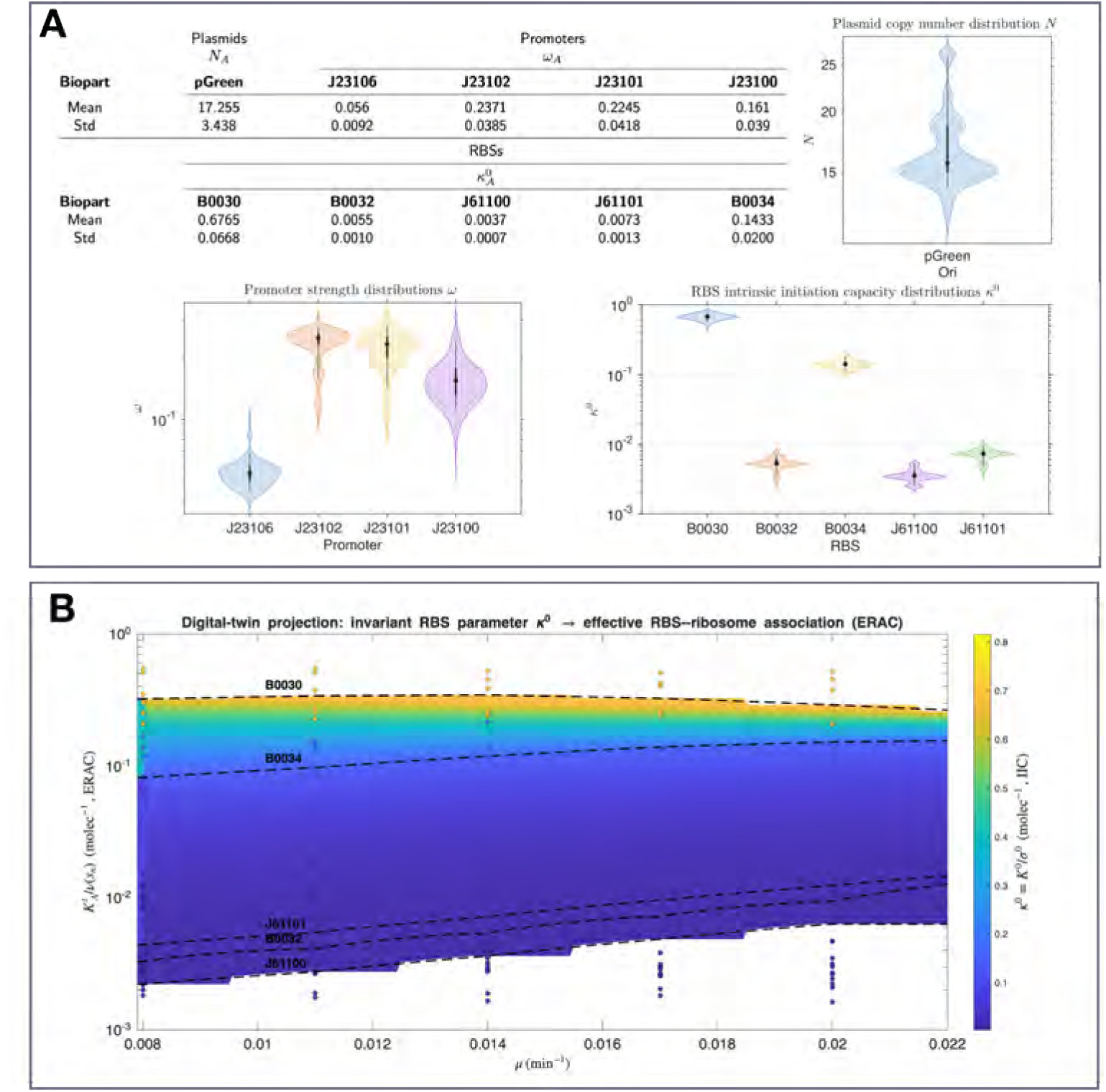
Biopart parameter identification and digital-twin projection of an invariant RBS descriptor. **A**, Estimated biopart parameters from the combinatorial library (shown as distributions across cross-validation runs). Plasmid copy number *N_A_* is inferred for each origin (with pSC101 anchored to its known value), promoter strengths are reported as transcription rates *ω*, and RBS translation properties are summarized by the intrinsic initiation capacity *κ*^0^ = *K*^0^*/σ*^0^ (IIC) and the sensitivity-related parameter (*ρ* as defined in Methods). Violin plots and summary statistics highlight the dispersion and stability of the inferred part parameters across genetic contexts (see Supplementary Section S10). **B**, Digital-twin projection from the invariant RBS parameter *κ*^0^ to an effective, growth-dependent initiation metric. The surface shows the median digital-twin mapping from (*µ, κ*^0^) to ERAC *≡ K^t^_A_ /ν*(*sn*) (units of molec*^−^*^1^), interpreted as an effective RBS–ribosome association constant under the inferred physiological state. Points correspond to TU-specific experimental means (one point per genetic context and growth condition). Dashed curves indicate iso-*κ*^0^ traces predicted by the model, illustrating how a single part-intrinsic parameter generates a continuum of effective initiation behaviour across host growth states and collapses context-dependent measurements onto low-dimensional manifolds.

Crucially, parameters identified from 𝓛_24_ could then be reused as fixed anchors to characterise libraries with reduced combinatorial diversity. In library 𝓛_6_, which introduces the RBS B0034 in new genetic contexts, inheriting all previously identified parameters enabled reliable identification of the new RBS intrinsic initiation capacity despite the limited library size. The same incremental logic was applied to library 𝓛_5_, which probes the promoter J23100 in combination with previously characterised RBSs. In this case, inheriting RBS parameters allowed accurate estimation of the promoter transcription rate while revealing context-specific deviations analysed further below.

Together, these results demonstrate that combinatorial TU libraries can be systematically exploited for incremental biopart identification, provided that identifiability constraints are explicitly accounted for and parameter inheritance is used to control model dimensionality.

Having established a robust incremental identification strategy across combinatorial libraries, we next examine the nature of the RBS parameters identified by this approach, and show how a single invariant descriptor captures translation initiation behaviour across diverse genetic and physiological contexts.

### 2.4 Identification of intrinsic RBS parameters and invariant initiation capacity

Within the host-aware synthesis model, each ribosome binding site (RBS) is characterised by two parameters: a nominal initiation strength *K*^0^ and a sensitivity parameter *σ*^0^ that captures how initiation effectiveness is modulated by translational demand and ribosomal flux (Methods 3.3). While both parameters influence the effective translation rate, they are not independently identifiable across the experimental regimes explored here.

Instead, parameter estimation across combinatorial libraries consistently reveals that the ratio

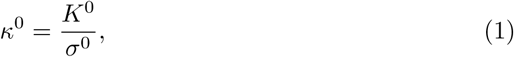

is robustly identifiable and transferable across genetic contexts and growth conditions. We refer to *κ*^0^ as the *intrinsic initiation capacity* (IIC) of an RBS, as it quantifies the inherent ability of the RBS to recruit ribosomes while explicitly accounting for its sensitivity to translational demand. Importantly, *κ*^0^ is invariant with respect to host growth state and resource availability, in contrast to effective translation metrics that depend on cellular context.

Using the core library 𝓛_24_ and the reduced translation model with fixed 1*/σ*^0^ = 0.02, we obtained well-separated posterior distributions for *κ*^0^ across all five RBSs (Figure 3A). These distributions remain stable under leave-one-out cross-validation and are preserved upon library extension to 𝓛_6_ (RBS B0034) and 𝓛_5_ (promoter J23100), demonstrating the transferability of *κ*^0^ across incremental library designs (Supplementary Information S5).

#### From invariant RBS capacity to effective translation across growth states

Although *κ*^0^ is invariant, experimental measurements access only context-dependent quantities, such as the effective translation rate *K_A_^t^*, which varies with growth rate *µ*, ribosomal flux *φ*, and nutrient-dependent elongation kinetics. The digital twin provides an explicit mapping between these layers by projecting the intrinsic parameter *κ*^0^ onto a growth-conditioned effective RBS–ribosome association constant,

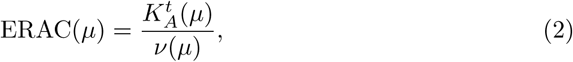

which represents an effective initiation metric under the inferred physiological state and can be directly compared across growth conditions.

Figure 3B visualises this projection as a continuous surface in the (*µ, κ*^0^) plane. Isocurves of constant *κ*^0^ define families of growth-dependent effective translation behaviours, while experimental transcriptional units appear as scattered points on this surface, corresponding to different host states and genetic contexts. This representation clarifies the distinction between *what experiments observe*—context-dependent effective translation—and *what the model identifies*—an invariant, biophysical RBS descriptor.

The estimated values of ERAC span approximately 0.05–0.3 molecule^−1^, consistent with reported effective initiation constants inferred from independent experimental and modelling approaches [39, 40]. Unlike conventional effective translation metrics, however, *κ*^0^ is explicitly disentangled from growth-dependent resource allocation, enabling absolute and transferable RBS characterisation.

#### Implications for modular design and prediction

By separating invariant RBS properties from physiological modulation, the host-aware framework reconciles apparent context dependence with underlying modularity. Once *κ*^0^ is identified, effective translation rates under new growth conditions or in novel genetic assemblies can be predicted by propagating *κ*^0^ through the digital twin, without re-estimating RBS-specific parameters. This establishes *κ*^0^ as a fundamental descriptor linking DNA sequence to translation efficiency across host states and provides a principled bridge between biophysical RBS characterisation and circuit-level performance.

### 2.5 Detection of context-dependent promoter–RBS non-modularity

A central assumption underlying modular genetic design is that promoters and ribosome binding sites (RBSs) contribute independently to gene expression. Within our framework, this assumption translates into promoter-specific transcription rates and RBS-specific intrinsic translation parameters that are consistent across genetic contexts. Deviations from this behaviour can thus be interpreted as signatures of biological non-modularity.

Across the combinatorial libraries 𝓛_24_ and 𝓛_30_, parameter estimates for promoter strengths *ω* and RBS intrinsic initiation capacities *κ* were internally consistent and unimodal (see Supplementary Information S10 for full parameter distributions), supporting the validity of the modular description for the vast majority of constructs (Fig. 3A; Supplementary Informations S6–S9). In particular, the inferred promoter strengths were stable across different RBS and origin-of-replication contexts, and RBS parameters remained invariant across promoters, as expected under a modular architecture.

In contrast, a distinct behaviour was observed for constructs containing the promoter J23100. When analysing the reduced library 𝓛_5_, which isolates J23100 in combination with a subset of previously characterised RBSs, the inferred transcription rate *ω*_J23100_ exhibited a clear bimodal distribution (Fig. 3A; Supplementary Information S9). This bimodality was not observed for the same RBSs under other promoters, nor for other promoters within the same plasmid backbone, indicating that it cannot be attributed to experimental variability or to limitations of the estimation procedure. Importantly, the host-aware digital twin remained internally consistent in this regime: growth-rate predictions, inferred resource fluxes, and synthesis-rate fits showed no systematic deviations (Supplementary Fig. S9.2). Rather than signalling model breakdown, the anomalous parameter estimates point to a genuine context-dependent interaction between the promoter and the RBS sequences. A plausible mechanistic explanation is altered accessibility of the 5^′^ untranslated region, arising from promoter-dependent transcription start sites and local RNA secondary structure, which in turn modulates effective translation initiation. Such promoter–RBS coupling effects have been previously reported for uninsulated Anderson promoters and synthetic constructs lacking standardized 5^′^ insulation elements [29, 41].

From a methodological perspective, this result highlights a key advantage of the proposed framework: the digital twin does not enforce modularity, but instead provides a quantitative diagnostic for its violation. Localized inconsistencies in inferred biopart parameters can be traced back to specific genetic contexts, allowing non-modular interactions to be identified rather than absorbed into effective, context-dependent parameters. In practice, such effects can often be mitigated by the use of standardized 5^′^ insulators or bicistronic design elements, restoring promoter–RBS independence and improving predictability.

Overall, the selective breakdown of modularity observed for J23100—contrasted with the consistency of the remaining libraries—further supports the robustness of our identification strategy. At the same time, it demonstrates its ability to reveal subtle, sequence-dependent context effects that are typically obscured in purely phenomenological or context-averaged characterisation approaches.

### 2.6 Summary of the identified biopart parameter set

By combining a host-aware digital twin with a structured combinatorial library design, we obtained a compact and internally consistent set of biopart parameters that quantitatively link DNA-level components to context-dependent gene expression phenotypes. Across the successive libraries analysed (𝓛_30_, 𝓛_24_, 𝓛_6_ and 𝓛_5_), this approach enabled the incremental identification of plasmid copy numbers, promoter strengths, and ribosome binding site (RBS) parameters with clear biophysical interpretation.

For promoters and origins of replication, the inferred parameters correspond to absolute transcriptional capacities and effective gene copy numbers that are transferable across genetic backgrounds, except in clearly localized cases of context-dependent non-modularity (Section 2.5). For RBSs, we identified an intrinsic, context-invariant parameter—the intrinsic initiation capacity *κ*^0^—that captures the ability of an RBS to recruit ribosomes independently of host growth state, while explicitly accounting for its sensitivity to translational demand (Section 2.4).

Together, these parameters form a minimal but sufficient description of transcriptional units within the explored design space: host-independent where possible, and explicitly host-aware where required. Importantly, the incremental estimation strategy—guided by practical identifiability analysis—allowed each new library to expand the characterized parameter set without re-estimating previously identified components (Supplementary Information S5).

This resulting parameter set establishes a quantitative and reusable mapping from biopart DNA sequences to biophysical parameters, and from these parameters to effective synthesis rates across cellular growth states. As such, it provides the foundation for predictive, modular, and scalable model-based design of synthetic transcriptional units under realistic host constraints. Taken together, this parameter set constitutes a minimal, identifiable and physiologically grounded coordinate system for transcriptional-unit design

## 3 Methods

### 3.1 Library plasmids

We constructed structured combinatorial libraries of constitutive transcriptional units (TUs) to enable model-based identification of intrinsic biopart parameters across diverse genetic and physiological contexts (Supplementary Information S2.1). Each TU expresses GFPmut3 under the control of a constitutive promoter and ribosome binding site (RBS), cloned into a plasmid backbone with a defined origin of replication (ORI). The library architecture deliberately reuses each biopart across multiple TUs, ensuring that copy-number, transcriptional, and translational contributions are systematically disentangled through host-aware, model-based fitting.

#### Core and extended combinatorial libraries

The core library, 𝓛_24_, comprises all 2 × 3 × 4 = 24 combinations of two ORIs (pSC101, pGreen), three constitutive promoters (J23106, J23102, J23101), and four RBSs (B0030, B0032, J61100, J61101). This library provides dense combinatorial coverage and serves as the reference dataset for identifying plasmid copy numbers, promoter transcription rates, and intrinsic RBS parameters.

An extended library, 𝓛_30_, was constructed by incorporating an additional RBS (B0034) across the same ORI and promoter set, yielding six new transcriptional units. The full library 𝓛_30_ was used to analyse practical identifiability, assess parameter robustness under increased combinatorial diversity, and inform model reduction decisions in the translation module.

#### Incremental promoter characterisation sublibrary

To evaluate incremental biopart identification under limited combinatorial excitation, we additionally constructed a reduced sublibrary (𝓛_5_) containing only pGreen-based constructs expressing promoter J23100 in combination with a subset of previously characterised RBSs. This sublibrary enables promoter parameter estimation while inheriting ORI- and RBS-specific parameters identified from 𝓛_24_ and 𝓛_30_, providing a stringent test of parameter transferability and modularity.

#### Construct identifiers and sequences

A complete mapping of transcriptional unit identifiers to ORI–promoter–RBS combinations is provided in Supplementary Information, Table S2, together with assembly details and metadata in Supplementary Information S2.1. Reporter, terminator, and backbone elements were kept constant across constructs unless explicitly stated otherwise.

### 3.2 Experimental measurements and data processing

#### Cultivation and measurements

All transcriptional unit (TU) variants were assayed in microtiter plates under batch-growth conditions, with time-resolved measurements of optical density and fluorescence acquired at 5-min intervals. Each construct was tested across multiple independent experimental instances (performed on different days) and multiple replicate wells per instance, enabling separation of biological variability from experimental noise. Detailed culture conditions, media composition, and replication structure are provided in Supplementary Information S2.1.

#### Calibration to standardized units

Raw optical density and fluorescence signals were converted into calibrated, comparable units using established protocols: *Equivalent Particle Count* (Particles) for absorbance and *Molecules of Equivalent Fluorescein* (MEFL) for fluorescence [34, 35, 42, 43] (Supplementary Information S2.2). This calibration enables absolute comparison of synthesis rates across constructs, experiments, and growth conditions. The reporter signal per cell is denoted as *p*(*t*), expressed in MEFL·Particle^−1^.

#### Growth rate estimation

Specific growth rates were computed from calibrated particle counts as

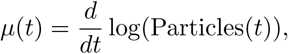

using a fourth-order central-difference scheme combined with Gaussian smoothing and uniform filtering criteria across all constructs (Supplementary Information S2.3). For subsequent analyses, only data within the exponential growth regime—between lag phase and stationary/death phases—were retained, as described below. The resulting growth-rate trajectories constitute the primary physiological observable driving the host-aware digital twin.

#### Synthesis rate estimation

Assuming that protein dilution by cell growth dominates over active degradation, the apparent synthesis rate Π(*t*) was obtained from

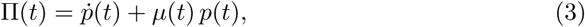

which corresponds to rearranging Eq. (5). Time derivatives were computed using central-difference methods (Supplementary Information S2.3). To minimise artefacts associated with non-steady-state dynamics, data corresponding to the lag phase (prior to reaching the maximum specific growth rate) and the death phase (after reaching maximum cell density) were excluded. All synthesis rates reported in the manuscript are expressed in MEFL·Particle^−1^·min^−1^ (equivalently, molecules per cell per minute under the applied calibration).

#### Background correction

Autofluorescence and non-specific background contributions were corrected using a dedicated background construct (pARKA) expressing LuxI, selected to match the reporter protein size and expression burden. The apparent background synthesis rate was modelled as a function of the specific growth rate *µ* and subtracted from TU measurements to obtain corrected synthesis rates. The background correction procedure and its validation are described in Supplementary Information S2.3.

#### Burden-related quantities

Where required (Results and Discussion), the translational burden imposed by each TU was quantified as the fraction of cellular proteome or translational resources allocated to exogenous expression, using mass-balance expressions of the form

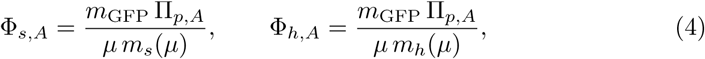

evaluated at the maximum specific growth rate *µ* = *µ*_m_ (Supplementary Information S3.4), with the strain- and host-specific protein-mass functions *m_s_*(*µ*) and *m_h_*(*µ*) defined in Supplementary Information S3.

### 3.3 Resource-aware TU synthesis model

We model each transcriptional unit as a constitutive expression cassette producing a reporter protein *A* (GFPmut3 in this work). Over the experimental time scales considered here, active protein degradation is negligible compared to dilution by growth, yielding

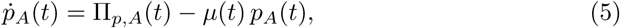

where *p_A_*is the number of protein molecules per cell, *µ* is the specific growth rate, and Π*_p,A_* is the synthesis rate.

#### Ribosome-limited synthesis rate

Following the host-aware formulation in [15], we treat ribosomes as the dominant shared limiting resource for translation. This formulation enables synthesis rates to be expressed explicitly as functions of host physiological state. The synthesis rate is written as

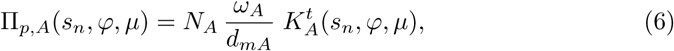

where *N_A_* is gene copy number (ORI-dependent), *ω_A_* is the transcription rate (promoter-dependent), *d_mA_* is the mRNA degradation rate, and *K_A_^t^* is an effective translation rate that combines intrinsic RBS properties with cellular context. Context enters through the flux of free ribosomes *φ* = *µr* and through nutrient-dependent elongation kinetics.

#### Normalized substrate and elongation factor

To avoid dependence on experiment-specific Monod parameters, we introduce a normalized substrate variable *s_n_* = *s/K_s_* and define an elongation factor *f* (*s_n_*) ∈ [0, 1] such that *ν*(*s_n_*) = *ν*_max_*f* (*s_n_*), where *ν* is the peptide elongation rate. The relationship between extra-cellular substrate, intracellular availability, and *f* (*s_n_*) is derived in Supplementary Information S3.1 and S3.5.

#### RBS parametrization and intrinsic initiation capacity

Translation initiation is parametrized by two intrinsic RBS parameters: a nominal strength *K_A_*^0^ and a sensitivity parameter *σ_A_*^0^ capturing how initiation effectiveness varies with elongation demand and ribosomal flux (Supplementary Information S3.1). Under this parametrization, the effective translation rate can be written as

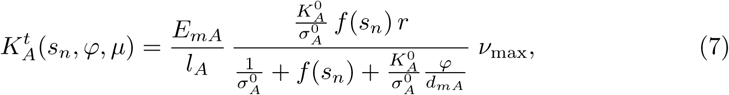

where *𝓛_A_* is the protein length and *E_mA_* captures polysome effects. This formulation separates intrinsic biopart parameters (*N_A_, ω_A_, K_A_*^0^*, σ_A_*^0^) from host- and environment-dependent variables (*µ, φ, f*).

We define the ratio *κ*^0^ = *K*^0^*/σ*^0^ as the *intrinsic initiation capacity* (IIC) of an RBS, which quantifies its inherent ability to recruit ribosomes independently of host growth state while accounting for its sensitivity to translational demand.

### 3.4 Host-cell endogenous model (HEM)

The Host Equivalent Model (HEM) captures the endogenous growth physiology of *E. coli* in the absence of exogenous expression, it is parameterised independently of the synthetic circuits considered in this work, and provides a mapping between nutrient availability and the host translational state. In particular, it predicts (i) the specific growth rate *µ* and (ii) the flux of free ribosomes *φ* as functions of the normalized substrate factor *f* (*s_n_*) and the endogenous resource-allocation terms (Supplementary Information S3.2).

In compact form, the growth rate is

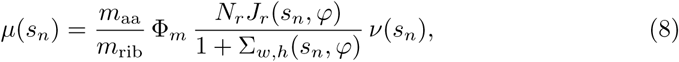

where *m*_aa_ is the average amino-acid mass, *m*_rib_ is the ribosomal protein mass, Φ*_m_*is the mature ribosome fraction, and *J_r_* is the resource recruitment strength (RRS) of ribosomal genes. The term Σ*_w,h_* denotes the weighted sum of RRS contributions from endogenous ribosomal and non-ribosomal genes (Supplementary Information S3.6).

The corresponding flux of free ribosomes *φ* = *µr* is obtained from an implicit relationship of the form

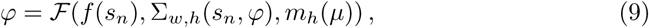

where *m_h_*(*µ*) is the host protein mass content as a function of growth rate (Supplementary Information S3.7). In practice, Eq. (8)–(9) are solved iteratively as described in Methods 3.5.

The HEM parameter values used in this work and the validation against literature ribosome allocation trends are reported in Supplementary Information S3.2 (see also Fig. S3.3).

### 3.5 Host–circuit interaction and model-in-the-loop digital twin

Exogenous gene expression alters host growth by diverting translational resources. In the host-aware formalism, this effect appears as an additional load term *θ* added to the endogenous weighted resource-allocation sum:

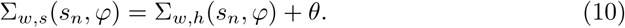

Here, *θ* quantifies the effective translational demand of the exogenous TU (Supplementary Information S3.3).

#### Circuit load model

For a constitutive TU with parameter set Θ = {*N_A_, ω_A_, κ_A_*^0^*, ρ_A_*^0^*, 𝓛_A_, . . .*}, the load is written as

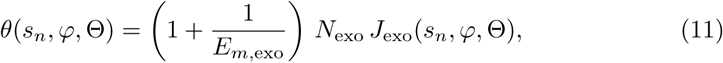

where *J*_exo_ is the resource recruitment strength of the exogenous gene(s) (Supplementary Information S3.1).

#### Digital twin (model-in-the-loop) inference using experimental growth rate

When experimental data are available, we use the measured growth rate *µ*^∗^ as a proxy for the physiological state, and infer a *consistent* pair (*φ*^∗^*, f* ^∗^(*s_n_*)) that satisfies both the host and circuit constraints. Concretely, the HEM defines a host relationship between (*φ, f*) and *θ* through Eq. (10), while the TU model defines a circuit relationship between (*φ, f*) and *θ* through Eq. (11). Their intersection yields (*φ*^∗^*, f* ^∗^) and a consensus load *θ*^∗^ for each experimental growth measurement. The detailed algorithm and implementation are given in Supplementary Information S3.3, Algorithm 2.

Finally, (*φ*^∗^*, f* ^∗^) are injected into the TU synthesis model (Methods 3.3) to evaluate Π*_p,A_* for parameter estimation and prediction (Results 2.2 and subsequent sections).

Using the measured growth rate *µ*^∗^ to drive the digital twin decouples host–circuit–environment interactions. Rather than requiring explicit knowledge of external substrate levels or endogenous allocation states, the measured growth rate constrains the effective elongation regime and ribosomal flux experienced by the TU. This improves practical predictability and stabilizes the mapping between biopart parameters and observed synthesis rates across experimental contexts (Supplementary Information S3.3 and S4).

### 3.6 Parameter estimation

Model parameters were estimated by minimizing the discrepancy between experimentally measured and model-predicted synthesis rates across combinatorial libraries of transcriptional units (TUs). All estimation problems were formulated as nonlinear, nonconvex optimization tasks and solved using a combination of the Enhanced Scatter Search (eSS) metaheuristic implemented in the MEIGO toolbox [44] and Bayesian Adaptive Direct Search (BADS) [45].

The objective function aggregates synthesis-rate errors across all measured growth-rate values, TUs, and experimental replicates included in each library. In all cases, parameters were estimated using the model-in-the-loop digital-twin formulation described in Methods Sections 3.3–3.5, in which the experimentally measured specific growth rate *µ* is injected into the host digital twin to infer a consistent cellular resource state prior to synthesis-rate prediction.

Parameter-estimation problems were defined at different levels of model complexity depending on the library. Specifically, we performed: (i) full estimation of copy-number, promoter, and RBS parameters (*κ*^0^*, ρ*^0^) for the extended library 𝓛_30_; (ii) full estimation of copy-number, promoter, and RBS *κ*^0^ parameters for the base library 𝓛_24_; and (iii) incremental estimation of new bioparts (e.g. RBS B0034) in extended libraries by inheriting previously characterised parameters from 𝓛_24_. Parameters known *a priori* (e.g. the copy number of pSC101) were fixed and excluded from the optimization.

Solver settings followed standard recommendations for both eSS and BADS and required no extensive manual tuning. For each estimation problem, multiple independent runs were performed to assess convergence consistency. Reported parameter values correspond to the best-performing solutions according to a normalized cost function. Full details on objective-function definition, solver configuration, and convergence assessment are provided in Supplementary Information S4.

### 3.7 Sensitivity analysis and practical identifiability

Practical identifiability of biopart parameters was assessed *a posteriori* using local sensitivity analysis of the synthesis rate with respect to the estimated parameters. This analysis was performed using the experimentally calibrated digital-twin model and the parameter estimates obtained from each library fit.

For each TU and each experimentally observed growth rate *µ*, the digital twin provides analytical expressions for the relative sensitivities

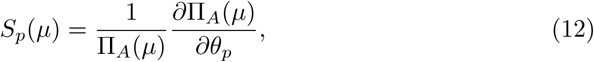

where *θ_p_* denotes a free model parameter. Absolute sensitivities were obtained by multiplying the relative sensitivities by the experimentally measured synthesis rate Π_A_^exp^(*µ*),

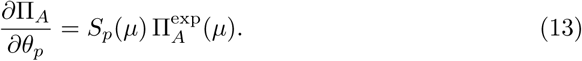

Stacking sensitivity rows across all TUs and growth-rate values yields the empirical Jacobian (sensitivity matrix) *J* (*θ*^^^) evaluated at the estimated parameter vector *θ*^^^. Only parameters treated as free in each estimation problem were included as columns of *J*, ensuring consistency between identifiability analysis and parameter-estimation settings.

Practical identifiability was quantified by singular value decomposition of the Jacobian, *J* = *USV* ^⊤^, and by defining the *effective rank* as the number of singular values exceeding a tolerance of 10^−2^ times the largest singular value. This criterion provides a robust estimate of the number of independent parameter combinations supported by the experimental data [7, 46, 47]. Sensitivity matrices were constructed using different levels of experimental aggregation (global TU means, experimental-instance means, and individual wells), yielding consistent effective-rank estimates.

A complete description of the sensitivity-analysis procedure, numerical implementation, and identifiability results for all libraries is provided in Supplementary Information S5.

## 4 Discussion

### 4.1 From relative tuning to absolute, host-aware biopart characterisation

Experimental characterisation of gene expression bioparts has traditionally relied on relative metrics obtained under narrowly defined experimental conditions. Promoter and RBS strengths are typically reported as dimensionless quantities, normalised to a reference construct and measured at a single growth state or environmental condition [27, 28]. While such approaches are effective for local tuning and comparative ranking, they provide limited predictive power when bioparts are reused in new genetic, physiological, or environmental contexts.

In this work, we argue that the central obstacle to absolute biopart characterisation is not the absence of mechanistic models, but the lack of experimental observables that sufficiently constrain host–circuit interactions. Gene expression does not occur in isolation: transcription and translation are coupled to cellular physiology through shared resources, and their effective rates vary systematically with the growth state of the cell. Ignoring this coupling inevitably yields parameters that are context-dependent by construction, even when derived from mechanistic equations.

Our approach reframes biopart characterisation as a system identification problem in which absolute, biologically interpretable parameters can be recovered only if the cellular state is explicitly accounted for. To this end, we leverage the specific growth rate as a low-dimensional yet information-rich physiological observable. Growth rate integrates nutrient availability, endogenous metabolic allocation, and the translational burden imposed by synthetic gene expression [5, 31], and thus provides a natural proxy for the internal state of the host cell.

By embedding growth-rate measurements into a host-aware digital twin, we obtain a model-in-the-loop framework in which host physiology is not treated as an unobserved disturbance, but as a measured input. This strategy does not eliminate context dependence; rather, it makes it explicit and quantifiable. Biopart-specific parameters are defined as intrinsic quantities that remain invariant across contexts, while effective expression rates emerge from their interaction with the growth-dependent cellular state. Absolute characterisation is therefore achieved not by ignoring the host, but by accounting for it in a principled and experimentally grounded manner.

### 4.2 Digital twins as a bridge between physiology, identifiability, and modular design

A central contribution of this work is the use of a host-aware digital twin not merely as a predictive model, but as an enabling abstraction that reconciles physiological realism with parameter identifiability and modular biopart characterisation. While host-aware models are increasingly used to explain context dependence [10, 31], their practical adoption for quantitative biopart characterisation has remained limited due to model complexity, unobservable intracellular variables, and identifiability constraints.

The key methodological step is conditioning the digital twin on the experimentally measured specific growth rate. Growth rate acts as a low-dimensional yet information-rich proxy for the cellular state, integrating nutrient availability, endogenous metabolism, and circuit-induced burden [5]. By anchoring the host model to the observed growth trajectory, the digital twin strongly constrains degrees of freedom that would otherwise remain practically unidentifiable, collapsing high-dimensional host–circuit–environment interactions onto a physiologically consistent manifold.

From an identifiability perspective, this model-in-the-loop strategy plays a role analogous to experimental feedback control. Conditioning on growth rate regularises inference by eliminating compensatory trade-offs between ribosome availability, elongation kinetics, and circuit load. Importantly, this does not require fine-grained molecular measurements, but relies solely on bulk growth and expression data, making the approach experimentally scalable.

The digital twin thus serves as a conceptual and computational bridge between mechanistic host-aware modelling and library-based system identification. By retaining sufficient physiological structure while remaining identifiable from standard datasets, it enables stable and interpretable parameter estimation at the biopart level. More broadly, this work highlights a general role for digital twins in synthetic biology: not only as predictive simulators, but as identifiability-enhancing abstractions that transform context dependence from a source of variability into a source of information.

### 4.3 Design implications and scalability: incremental and patchwork library characterisation

A major challenge in quantitative biopart characterisation is scalability. Exhaustive combinatorial libraries rapidly become experimentally impractical as the number of parts increases. Our results show that such exhaustive designs are not required, provided that experimental design and model structure are aligned with identifiability principles.

Practical identifiability analysis reveals that parameter identifiability is governed primarily by the diversity of genetic contexts and host states, rather than by the number of measurements or experimental replicates. This insight motivates an incremental, *patchwork* strategy for library design, in which small, carefully constructed sublibraries are combined to progressively identify new bioparts while reusing previously characterised ones as anchors.

In this framework, an initial base library establishes a well-identified reference parameter set. Subsequent library extensions—such as the introduction of new RBSs or promoters—require estimation only of the newly introduced parameters, while previously identified components are inherited. This strategy is formally supported by the effective-rank analysis of the sensitivity matrix, which shows that beyond a certain level of combinatorial diversity, additional contexts improve robustness without increasing the intrinsic dimensionality of the identifiable parameter space.

By conditioning inference on growth rate, the host-aware digital twin enables heterogeneous datasets collected across different days, conditions, and partial libraries to be integrated into a coherent estimation framework. Together, these results outline a scalable roadmap for biopart characterisation that replaces monolithic calibration with modular, extensible system identification.

### 4.4 Intrinsic RBS parameters and the limits of modularity

A central conceptual result of this work is the identification of intrinsic, biophysically interpretable parameters that characterise ribosome binding sites (RBSs) across genetic and physiological contexts. In particular, we show that the ratio

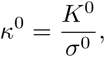

which we term the *intrinsic initiation capacity* (IIC), behaves as a structural invariant of the RBS sequence. Across a wide range of transcriptional units, plasmid copy numbers, and host growth states, *κ* remains stable and transferable, while effective translation rates vary substantially with cellular physiology.

This distinction clarifies the relationship between intrinsic biopart properties and commonly used context-dependent expression metrics. Effective translation rates and related quantities collapse multiple physiological effects—ribosome availability, elongation kinetics, and resource competition—into a single observable. While valuable for tuning expression within a fixed regime, such metrics are not intrinsic properties of the RBS and cannot be expected to transfer across growth conditions. In contrast, *κ* captures the inherent ability of an RBS to recruit ribosomes in a resource-normalised manner, with context-dependent expression emerging as a projection of this invariant parameter onto different physiological states.

Complementary to *κ*, the parameter *ρ*^0^ = 1*/σ*^0^ sets the scale over which translation initiation responds to changes in translational demand. For the RBSs studied here, inferred values of *ρ*^0^ ≈ 0.02 indicate that effective initiation remains strongly coupled to cellular physiology over the explored growth range. Consistent with the practical identifiability analysis, *ρ*^0^ is not independently identifiable in this regime and is therefore best interpreted as a gauge parameter that fixes the projection of *κ* onto growth-dependent effective translation rates, rather than as a transferable biopart descriptor in its own right.

Importantly, the identification of intrinsic RBS parameters does not imply full modularity of gene expression. On the contrary, our analysis reveals that departures from modularity can be detected and localised within the same framework. The promoter J23100 provides a clear example, exhibiting context-dependent behaviour consistent with promoter–RBS sequence interactions that alter 5^′^-UTR structure and ribosome accessibility. Rather than masking such effects, the host-aware digital twin exposes them as localised violations of parameter invariance.

Taken together, these results delineate both the power and the limits of modularity in synthetic biology. By separating intrinsic biopart properties from physiological projections, host-aware modelling enables robust parameter reuse while simultaneously providing a diagnostic lens to identify genuine biological non-modularity when it arises.

## 5 Conclusions

The central challenge of model-based synthetic biology is not merely to describe gene expression dynamics, but to establish a quantitative and reusable mapping between DNA sequences, intrinsic biopart properties, and circuit-level phenotypes across cellular contexts. In this work, we have addressed a key missing component of this challenge by introducing a context- and host-aware framework for the absolute characterisation of bioparts embedded in combinatorial transcriptional unit libraries.

By treating biopart characterisation as a system identification problem, and by leveraging a host digital twin driven by experimental growth-rate measurements, we demonstrate that intrinsically meaningful parameters—including plasmid copy number, promoter strength, and the intrinsic RBS initiation capacity *κ*—can be reliably identified despite strong host–circuit coupling. Importantly, the combinatorial structure of the libraries transforms experimental context variability from a source of confounding into a source of identifiability, enabling incremental and scalable library expansion through parameter inheritance.

Beyond parameter estimation, the framework presented here provides a diagnostic lens on biological modularity. The emergence of context-dependent deviations, exemplified by the J23100 promoter–RBS interactions, highlights how host-aware models can simultaneously support prediction and reveal genuine sequence-level non-modularity. This dual predictive–diagnostic capability is essential for advancing synthetic biology from relative tuning toward principled engineering.

More broadly, our results establish the foundation for closing the Design–Build–Test–Learn (DBTL) loop in synthetic biology. Intrinsic biopart parameters inferred in a host-aware manner define explicit design targets for model-based circuit construction. When combined with inverse sequence-design approaches that map desired expression properties to concrete DNA sequences [48], this framework enables a fully model-driven workflow: from system-level specifications, to intrinsic biopart requirements, to sequence-level implementation.

Together, these results establish host-aware, model-based intrinsic characterisation of gene expression bioparts as a viable foundation for predictive and modular synthetic biology, enabling the reuse of quantitative biopart descriptions across contexts, projects, and host organisms, and supporting the transition from bespoke circuit construction to scalable, engineering-grade biological design.

## Supplementary information

This article has accompanying supplementary files.

## Funding

This work was supported by MICIU/AEI/10.13039/501100011033 (grant PID2023-151077OB-I00, DBTL4SYNBIOCON). A. Arboleda-Garcia was supported by Universitat Politècnica de València (grant PAID-01-21397). Y. Boada was supported by Universitat Politècnica de València (grant PAID-06-24) and by the Secretaŕıa de Educacíon Superior, Ciencia, Tecnoloǵıa e Innovacíon of Ecuador (Scholarship Convocatoria Abierta 2011).

D.R.P. and J.R.B. acknowledge support by the European Commission – NextGenerationEU, through Momentum CSIC Programme: Develop Your Digital Talent. D.R.P. is hired under the Generation D initiative, promoted by Red.es, an organisation attached to the Ministry for Digital Transformation and the Civil Service, for the attraction and retention of talent through grants and training contracts, financed by the Recovery, Transformation and Resilience Plan through the European Union’s Next Generation funds. J. R. Banga acknowledge support by MICIU/AEI/10.13039/501100011033 and ERDF/EU (grant PID2023-146275NB-C22, DYNAMO-bio).

## Conflict of interest/Competing interests

The authors declare no competing interests.

## Ethics approval and consent to participate

Not applicable

## Consent for publication

Not applicable

## Data availability

All experimental data supporting the findings of this study are available within the paper and its Supplementary Information. Processed datasets and scripts used to generate the figures are available from the corresponding author upon reasonable request. Raw data and analysis-ready datasets will be deposited in a public repository prior to publication.

## Materials availability

Plasmids and biological materials generated in this study are available from the corresponding author upon reasonable request.

## Code availability

All custom code used for model simulation, parameter estimation, and data analysis is available from the corresponding author upon reasonable request. The code will be made publicly available upon publication.

## Author contributions

J.P., A.V. and J.R.B. conceived the study and contributed to project funding acquisition. J.P. developed the theoretical framework, designed the digital-twin-based methodology, implemented the core computational framework, analysed the data, and wrote the manuscript. A.A.-G. performed the experimental work, including plasmid construction and cultivation experiments. A.V. and Y.B. contributed to experimental design, data processing, and standardisation of measurements. D.R.P. developed and executed global optimisation workflows and carried out global parameter estimation analyses. J.R.B. contributed to identifiability analysis and provided conceptual guidance on optimisation strategies, model calibration, and interpretation. All authors discussed the results and contributed to the final manuscript.

## Declaration of AI Assistance

The authors used AI-assisted tools to support language editing and text refinement. The scientific content, analysis, and conclusions were fully conceived and validated by the authors.

## Notes

### Competing Interest Statement

The authors have declared no competing interest.

### Summary of Updates

This version includes minor typographical corrections, improved figure S3.4 and Table S7 clarity, in the Supplementary Information. The main results and conclusions remain unchanged.

