## Supplementary Information for "Host-aware Identification of Intrinsic Gene Expression Biopart Parameters from Combinatorial Libraries"

---

* Jesús Picó

*Email addresses:* (Jesús Picó), (Andrés Arboleda-García), (David R. Penas), (Julio R. Banga), (Alejandro Vignoni), (Yadira Boada)

**S6 Characterization of library $\mathcal{L}_{24}$.** 30

**S7 Characterization of RBS B0034 using the library $\mathcal{L}_6$** 33

**S8 RBS intrinsic initiation capacity (IIC) and effective ribosome–RBS association capacity (ERAC)** 34

**S9 Characterization of promoter J23100 using the library $\mathcal{L}_5$** 35

**S10 Summary of estimated parameters and their distributions.** 37

### S1. General overview

This study introduces a host-aware, model-based methodology for the quantitative characterization of genetic bioparts embedded within a combinatorial library of constitutive transcriptional units (TUs). The approach enables the estimation of biologically meaningful parameters—plasmid copy number, promoter transcription rate, and ribosome binding site (RBS) translational efficiency—expressed in standardized units with direct biophysical interpretation. Together, these elements define a host-aware, digital-twin-guided workflow that provides a coherent and scalable framework for quantitative biopart characterization and predictive design of synthetic gene circuits. Figure S1.1 summarizes the full workflow.

**Combinatorial TU library and experimental characterization.** Each TU combines an origin of replication, promoter, and RBS driving expression of *GFPmut3*. The combinatorial design ensures that every biopart appears embedded in multiple distinct genetic and physiological contexts (Fig. S1.1A), which is essential for resolving parameter lack of identifiability that arise when TUs are studied in isolation. For each construct, high-resolution measurements of optical density and GFP fluorescence are used to compute time-resolved specific growth rates and TU synthesis rates (Fig. S1.1B).

**Host-aware modeling and digital twin.** To capture the coupling between synthetic gene expression and cell physiology, we employ a host-aware modeling framework combining: (i) an endogenous Host Equivalent Model (HEM) describing resource allocation and growth, and (ii) a mechanistic TU synthesis model parameterized by biopart-specific quantities.

To improve predictive capacity and mitigate uncertainty inherent to purely mechanistic descriptions, we introduce a hybrid *E. coli* digital twin (Fig. S1.1C) that integrates real-time experimental growth-rate measurements. The specific growth rate provides an informative proxy for the cellular resource state—reflecting substrate availability, endogenous metabolism, and gene expression burden—and thus constrains the host–circuit interaction in a mechanistically consistent manner Scott et al. (2010); Ceroni et al. (2015). This model-in-the-loop strategy enables the digital twin to infer internal resource fluxes and deliver context-aware predictions of TU synthesis rates.

**Context-independent parameterization of bioparts.** The TU synthesis model explicitly separates: (a) global physiological variables (growth rate, resource flux, substrate availability), determined by the digital twin, from (b) intrinsic biopart parameters (copy number, promoter strength, RBS nominal strength and sensitivity).

While the host state modulates translation capacity, the biopart-specific parameters remain context-independent. This separation enables a quantitative mapping from DNA sequences to biophysical parameters and from these to circuit-level phenotypes.

**Parameter estimation via global optimization.** Predicted synthesis rates from the digital twin are matched to experimental TU data through a global heuristic optimization procedure (Fig. S1.1D). This allows simultaneous estimation of all biopart parameters using the full combinatorial dataset and propagates uncertainty naturally through Monte Carlo sampling of the solution landscape. The resulting parameter distributions (Fig. S1.1E) are then used to compute confidence intervals for TU synthesis-rate predictions over the full range of growth conditions.

**Role of the combinatorial library.** The combinatorial structure of the TU library is essential for resolving structural and practical identifiability limitations inherent to individual constructs. Because each promoter, RBS, and origin appears in multiple distinct contexts, the global sensitivity (Jacobian) matrix becomes structured and effectively full-rank. This enables reliable, context-independent characterization of all bioparts and supports incremental expansion of the library by adding new parts whose parameters can then be inferred using previously characterized bioparts as anchors.

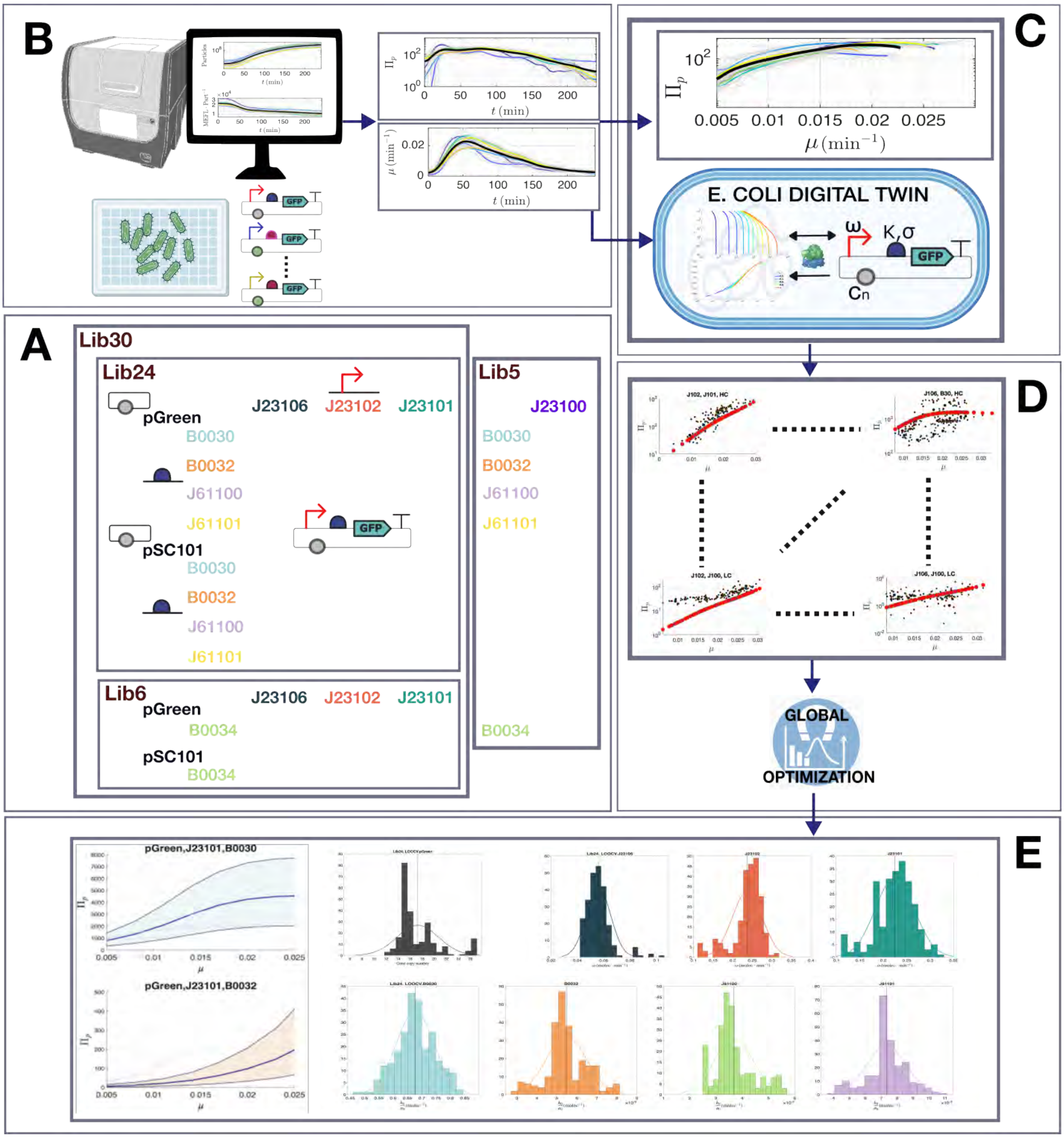

**Figure S1.1: Overview of the host-aware biopart characterization workflow.** (A) Combinatorial library of constitutive transcriptional units (TUs) constructed from multiple plasmid origins, promoters, and ribosome binding sites (RBSs), ensuring that each biopart appears across diverse genetic contexts. (B) High-resolution experimental characterization: time-series of optical density and GFP fluorescence are processed to obtain specific growth rate $\mu(t)$ and TU synthesis rate $\Pi(t)$ for every construct. (C) Host-aware *E. coli* digital twin integrating a mechanistic resource-allocation model with real-time experimental growth-rate measurements. The digital twin provides context-dependent estimates of cellular resource fluxes and predictions of growth-coupled TU synthesis dynamics. (D) Global optimization loop combining digital-twin predictions with experimental synthesis-rate data to estimate biologically interpretable biopart parameters (copy number, promoter transcription rate, and RBS translation efficiency). (E) Posterior parameter distributions and TU synthesis-rate fits across the library. These distributions quantify uncertainty and support context-independent, mechanistically grounded characterization of genetic bioparts.

### S2. TU library building and testing

#### *S2.1. Library plasmids*

A combinatorial library of transcriptional units (TUs) was constructed to enable host-aware characterization of individual genetic parts (plasmid origin, promoter, and ribosome binding site). The libraries were designed to span a broad dynamic range of expression while maintaining compatibility with common modular assembly standards and calibration workflows (see Supplementary Section S2.4).

**Design rationale.** Each TU consists of a single-copy reporter cassette encoding superfolder GFP (GFPmut3) under the control of a constitutive promoter and a ribosome binding site (RBS), followed by a transcriptional terminator to ensure proper transcriptional termination, and cloned in a vector backbone carrying a defined plasmid origin of replication (ORI). The resulting expression rate, $\Pi_p$, reflects the combined effects of gene copy number (determined by the plasmid origin), promoter transcriptional strength, and translational efficiency (RBS). By combining these parts systematically, we generated a set of TUs that differ in expression capacity and metabolic load but share identical reporter context, enabling direct comparison and model-based deconvolution of part-specific parameters.

**Constitutive promoters and RBSs.** Promoter variants were selected from the *E. coli* Anderson constitutive promoter collection, covering low to high transcriptional activity: J23106, J23102, and J23101. Translation initiation was modulated using four RBSs of differing predicted strength: B0030, B0032, J61100, and J61101. An additional RBS, B0034, and promoter, J23100, were later included to extend the dynamic range and to serve as an external reference for predictive validation of model transferability (see Supplementary Table S1). All RBS and promoter sequences were obtained from the iGEM Registry of Standard Biological Parts iGEM Foundation (2003).

**Plasmid origins of replication.** Two plasmid backbones differing in replication origin were used to modulate gene copy number and intracellular resource demand: pSC101 (low-copy origin, 5 copies per cell) and pGreen (medium-copy origin, approximately 20-25 copies per cell). These origins were chosen to represent typical low to moderate limits of replication and host load in *E. coli* 10G under aerobic growth.

**Combinatorial libraries.** The base library ($\mathcal{L}_{24}$) comprised all $2 \times 3 \times 4 = 24$ possible combinations of ORI, promoter, and RBS from the sets defined above. This library constitutes the core combinatorial design and was used to establish a consistent reference parameter set. An extended library ($\mathcal{L}_{30}$) was constructed by incorporating the additional RBS B0034 into $\mathcal{L}_{24}$, yielding six new constructs (denoted as sublibrary $\mathcal{L}_6$ when considered in isolation). The full library $\mathcal{L}_{30}$ was used to analyse practical identifiability, assess parameter robustness under increased combinatorial diversity, and inform the reduction of model parameters. Finally, a reduced sublibrary ($\mathcal{L}_5$), containing only constructs with promoter J23100, was employed to evaluate incremental biopart characterization under limited combinatorial excitation, while reusing previously inferred ORI and RBS parameters (see Supplementary Table S1).

**Reporter and terminators.** All constructs encode GFPmut3 (E0040) with a standard double terminator (B0015). Reporter calibration was performed using fluorescein-equivalent units (MEFL) and cell counts converted to calibrated particle numbers following Boada et al. (2019); Beal et al. (2022); González-Cebrián et al. (2023), ensuring quantitative comparability across experiments.

**Assembly and sequence verification.** All plasmids were assembled using the **Golden Gate method** with *BsaI-HFv2* (NEB) and *T4 DNA ligase* in 15 µL reactions containing 1 µL of each DNA module (promoter, RBS, CDS, terminator, and destination vector), 1.5 µL of 10× Thermo Buffer, 1.5 µL BSA, and nuclease-free water. The thermocycling protocol consisted of 25 cycles (37 °C for 3 min, 16 °C for 4 min) followed by 50 °C for 5 min and 80 °C for 10 min for enzyme inactivation. Assembled constructs were transformed into *E. coli* 10G (50 µL chemically competent cells) and plated on LB agar containing either 50 µg/mL kanamycin or 100 µg/mL chloramphenicol, then incubated overnight at 37 °C. Correct clones were identified by colony PCR using primers 204/205 (for pGreen plasmids) and 230/231 (for pSC101 plasmids) and analyzed on 1% agarose gels (1× TAE buffer, GeneRuler 1 kb Plus DNA Ladder).

- Forward 204: GCAACCTCTCGGGCTTCTGGAT
- Reverse 205: ACAGCGACTTAGTTTACCCGCCA
- Forward 230: AGCGGATAACAATTTCACACAGGAGGCC
- Reverse 231: GCCAGGGTTTTCCCAGTCACGAC

Positive clones were purified using the Qiagen Miniprep Kit, quantified spectrophotometrically, and verified by Sanger sequencing (Eurofins Genomics). All plasmid sequences were verified and deposited in Supplementary Table S2, including sequence features, part identifiers, and GenBank accessions.

**Culture and measurement conditions.** *E. coli* 10G cultures were grown in LB medium (10 g/L tryptone, 5 g/L yeast extract, 10 g/L NaCl) supplemented with the appropriate antibiotic at 37 °C with shaking at 220 rpm. Overnight cultures were diluted 1:100 into fresh medium and grown to $OD_{600} \approx 0.4$ before measurements. Optical density and fluorescence were recorded in black, clear-bottom 96-well plates using Cytation3 plate reader (37 °C, orbital shaking). OD was measured at 600 nm, and fluorescence was recorded using the appropriate filter sets (GFP: Ex/Em 485/510 nm). Detailed information on growth calibration and fluorescence quantification procedures is provided in Supplementary Section S2.2.

**Table S1: Library plasmids and part combinations.** Combinatorial transcriptional units (TUs) used for host-aware characterization. Each TU expresses GFPmut3 under a constitutive promoter and RBS, cloned in a plasmid backbone with a defined origin of replication (ORI). The base library ($\mathcal{L}_{24}$) includes all $2 \times 3 \times 4 = 24$ possible combinations of ORI, promoter, and RBS; the extended library ($\mathcal{L}_{30}$) adds the RBS B0034. Library $\mathcal{L}_5$ adds the promoter J23100 only with the plasmid ORI pGreen. When convenient, alternative IDs are used for the same TU emphasizing the library or sublibrary they belong to.

| IDs | ORI | Promoter | RBS | Reporter | Notes / Use |
|---|---|---|---|---|---|
| L30_01, L24_01 | pSC101 | J23106 | B0030 | GFPmut3 | Base library |
| L30_02, L24_02 | pSC101 | J23106 | B0032 | GFPmut3 | Base library |
| L30_03, L24_03 | pSC101 | J23106 | J61100 | GFPmut3 | Base library |
| L30_04, L24_04 | pSC101 | J23106 | J61101 | GFPmut3 | Base library |
| L30_05, L24_05 | pSC101 | J23102 | B0030 | GFPmut3 | Base library |
| L30_06, L24_06 | pSC101 | J23102 | B0032 | GFPmut3 | Base library |
| L30_07, L24_07 | pSC101 | J23102 | J61100 | GFPmut3 | Base library |
| L30_08, L24_08 | pSC101 | J23102 | J61101 | GFPmut3 | Base library |
| L30_09, L24_09 | pSC101 | J23101 | B0030 | GFPmut3 | Base library |
| L30_10, L24_10 | pSC101 | J23101 | B0032 | GFPmut3 | Base library |
| L30_11, L24_11 | pSC101 | J23101 | J61100 | GFPmut3 | Base library |
| L30_12, L24_12 | pSC101 | J23101 | J61101 | GFPmut3 | Base library |
| L30_13, L24_13 | pGreen | J23106 | B0030 | GFPmut3 | Base library |
| L30_14, L24_14 | pGreen | J23106 | B0032 | GFPmut3 | Base library |
| L30_15, L24_15 | pGreen | J23106 | J61100 | GFPmut3 | Base library |
| L30_16, L24_16 | pGreen | J23106 | J61101 | GFPmut3 | Base library |
| L30_17, L24_17 | pGreen | J23102 | B0030 | GFPmut3 | Base library |
| L30_18, L24_18 | pGreen | J23102 | B0032 | GFPmut3 | Base library |
| L30_19, L24_19 | pGreen | J23102 | J61100 | GFPmut3 | Base library |
| L30_20, L24_20 | pGreen | J23102 | J61101 | GFPmut3 | Base library |
| L30_21, L24_21 | pGreen | J23101 | B0030 | GFPmut3 | Base library |
| L30_22, L24_22 | pGreen | J23101 | B0032 | GFPmut3 | Base library |
| L30_23, L24_23 | pGreen | J23101 | J61100 | GFPmut3 | Base library |
| L30_24, L24_24 | pGreen | J23101 | J61101 | GFPmut3 | Base library |
| L30_25,L6_01 | pSC101 | J23106 | B0034 | GFPmut3 | Extended library (new RBS) |
| L30_26,L6_02 | pSC101 | J23102 | B0034 | GFPmut3 | Extended library (new RBS) |
| L30_27,L6_03 | pSC101 | J23101 | B0034 | GFPmut3 | Extended library (new RBS) |
| L30_28,L6_04 | pGreen | J23106 | B0034 | GFPmut3 | Extended library (new RBS) |
| L30_29,L6_05 | pGreen | J23102 | B0034 | GFPmut3 | Extended library (new RBS) |
| L30_30,L6_06 | pGreen | J23101 | B0034 | GFPmut3 | Extended library (new RBS) |
| L5_1 | pGreen | J23100 | B0030 | GFPmut3 | Extended library (new promoter) |
| L5_2 | pGreen | J23100 | B0032 | GFPmut3 | Extended library (new promoter) |
| L5_3 | pGreen | J23100 | B0034 | GFPmut3 | Extended library (new promoter) |
| L5_4 | pGreen | J23100 | J61100 | GFPmut3 | Extended library (new promoter) |
| L5_5 | pGreen | J23100 | J61101 | GFPmut3 | Extended library (new promoter) |

*S2.2. Optical density and fluorescence calibration*

We used the Cytation 3 Plate Reader (Agilent BioTek) to measure optical density ($\mathrm{OD}_{600}$) and fluorescence, which were converted to calibrated Particles and Molecules of Equivalent Fluorescein (MEFL), respectively, as described in Boada et al. (2019), Beal et al. (2022) and González-Cebrián et al. (2023). This standardization approach provides both a physical and a biological interpretation of the measurements.

In this framework, absorbance values can be expressed in terms of particle concentration, assuming that the particles have optical properties and scattering behavior comparable to *E. coli* cells Beal et al. (2020). Similarly, fluorescence values are expressed in MEFL units, which relate raw fluorescence measurements to the number of fluorescein-equivalent molecules. Thus, MEFL units enable a molecular-scale interpretation of fluorescence intensity, allowing the estimation of the average number of Green Fluorescent Protein (GFP) molecules per cell Boada et al. (2019).

*S2.3. Experimental evaluation of specific cell growth and TU synthesis rates*

**Specific growth rate.** The number of Particles per well was recorded every 5 minutes. To avoid approximation errors at the boundaries of the dataset, we first extrapolated three additional dummy points at each end using MATLAB's *polyfit* and *polyval* functions. The time derivative of the Particles signal was then computed using a fourth-order central difference expression:

$$\dot{y}_k = \frac{-y_{k+2} + 8y_{k+1} - 8y_{k-1} + y_{k-2}}{12\mathrm{dt}} \tag{1}$$

and divided by the number of Particles to obtain the sampled time series of the apparent growth rate. This series was smoothed using a Gaussian kernel with Matlab's *smoothdata* function.

The resulting estimate, $\hat{\mu}$, represents the apparent specific growth rate, which excludes maintenance and death contributions:

$$\frac{\dot{N}}{N} = \mu - d_m \tag{2}$$

We approximated $d_m$ following Biselli et al. (2020):

$$d_m = \frac{0.23}{60 \cdot 24} \cdot \exp\left(0.87 \cdot 60 \cdot \mu\right) \tag{3}$$

Since $\mu = \hat{\mu} + d_m$, we applied the correction

$$\mu = \hat{\mu} + \frac{0.23}{60 \cdot 24} \cdot \exp\left[0.87 \cdot 60 \cdot (\hat{\mu} + d_m)\right] \tag{4}$$

Finally, assuming $d_m \ll \hat{\mu}$, and a basal rate $d_m = 0.03\mu$ we obtained:

$$\mu = \hat{\mu} + \frac{0.23}{60 \cdot 24} \cdot \exp\left(0.9 \cdot 60 \cdot \hat{\mu}\right) \tag{5}$$

**Synthesis rate.** For each well, we obtained the number of MEFL per Particle as a function of time $p(t) = \mathrm{MEFL(t)/Particle(t)}$, and we obtained its time derivative $\dot{p}(t)$ using a fourth-order Central Difference Expression as above. Then, we calculated the synthesis rate as $\Pi(t) = \dot{p}(t) + \mu(t)p(t)$.

**Background correction** To account for the cell auto-fluorescence present in every experiment, we included 10 blank wells (containing only medium) and 10 background wells containing a background TU (pARKA21). The pARKA21 construct is a Level 1 constitutive expression cassette expressing the transcription factor LuxR, and containing the constitutive promoter J23106, the strong RBS B0030, the coding sequence C0062 (LuxR), and the standard bidirectional terminator B0015. The construct is cloned into a pARKA backbone carrying kanamycin resistance. LuxR is a protein with molecular weight similar to GFP. For this reason, pARKA21 was used as background signal to compensate for the fact that cell auto-fluorescence varies depending on the expression of exogenous proteins.

The blank wells were used to correct the MEFL measurements of the background construct pARKA21. For each replicate in every experiment, we extracted the MEFL and Particles data corresponding to pARKA21 from the time when cells reached their maximum specific growth rate until the point when the population achieved the maximum number of particles. This procedure avoided data from the batch phases dominated by cell adaptation and death, respectively.

Instead of following the common approach of subtracting the background fluorescence from the measured signal as a function of absorbance for each replicate and experiment, we hypothesized that a more accurate and physiologically relevant representation of cell auto-fluorescence could be obtained by considering the relationship between the apparent GFP synthesis rate of the background construct pARKA21 and the growth rate. This approach is agreement with our work, in which we characterize TUs using the relationship between synthesis rate and cell growth rate. The apparent auto-fluorescence synthesis rate varied only slightly with growth rate, with an average value around 2.5 $\mathrm{MEFL} \cdot \mathrm{Particle}^{-1} \cdot \mathrm{min}^{-1}$. Nevertheless, we used data from all experiments to fit the following model:

$$\Pi_{\mathrm{bk}}(\mu) = \frac{c_0 + c_1(10^2\mu)^{c_2}}{1 + c_3(10^2\mu)^{c_4}} \tag{6}$$

The estimated parameters were:

$$\begin{aligned} c_0 &= 1.550 \\ c_1 &= 5.041 \\ c_2 &= 1.058 \\ c_3 &= 0.730 \\ c_4 & \quad 2.980 \end{aligned}$$

Figure S2.1 shows the residuals of the model predictions. The residuals are centered around zero, with no systematic bias, confirming the consistency of the fitted model. From these results, we can infer the minimum transcriptional unit (TU) synthesis rates that can be reliably quantified above the background auto-fluorescence signal.

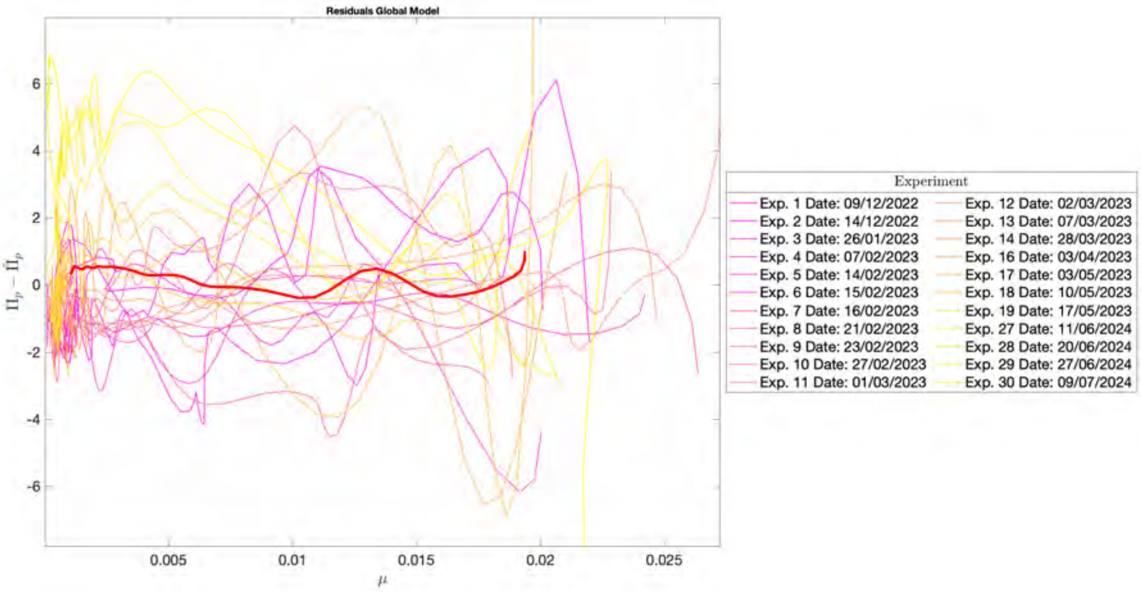

**Figure S2.1:** Background auto-fluorescence model residuals.

#### *S2.4. Experimental data*

We ran a a large set of experiments. In each experiment, several TU constructs were tested, each one with 10 replicas (wells), so that eventually, we had each TU tested in several independent experiments run on different days. Figure S2.2 shows the experimental results of synthesis rate as a function of cell growth for all the TUs. To minimize artefacts associated with non-steady-state dynamics, data corresponding to the lag phase (prior to reaching the maximum specific growth rate) and the death phase (after reaching maximum cell density) were excluded.

Figures S2.3 and S2.4 show the sorted experimental synthesis rates $(\mathrm{MEFL} \cdot \mathrm{Particle}^{-1} \cdot \mathrm{min}^{-1})$ evaluated at the maximum specific growth rate $(\mathrm{min}^{-1})$. Figure S2.3 includes the corresponding cell burden, evaluated as the estimated percentage of ribosomes allocated to the expression of each transcriptional unit. On the other hand, Figure S2.4 shows the Fano Factor and the Coefficient of Variation for the synthesis rate of each transcriptional unit, evaluated at $\mu_{\max}$. For each of the TUs, the representative values evaluated at the maximum cell growth rate are listed in Table S2.

**Table S2: Experimental results.** Synthesis rate given as $\Pi(\mu_m)$ $(\mathrm{MEFL} \cdot \mathrm{Particle}^{-1} \cdot \mathrm{min}^{-1})$. Cell loading $\Phi_h(\mu_m)$ expressed as percentage that the recruited exogenous cell resources represent w.r.t. the endogenous ones. Coefficient of variation and Fano factor for the synthesis rate. All are evaluated at the maximum cell growth rate $\mu_m$ $(\mathrm{min}^{-1})$.

| | $\Pi(\mu_m)$ | | $\mu_m$ | | $\Phi_h(\mu_m)$ | | $CV$ | $FF$ |
|---|---|---|---|---|---|---|---|---|
| TU code | Mean | $\sigma$ | Mean | $\sigma$ | Mean | $\sigma$ | | |
| L30_01, L24_01 | 229.8576 | 58.0663 | 0.0229 | 0.0041 | 0.1195 | 0.0281 | 0.2526 | 14.6686 |
| L30_02, L24_02 | 6.0711 | 4.6234 | 0.0208 | 0.0024 | 0.0035 | 0.0023 | 0.7615 | 3.5209 |
| L30_03, L24_03 | 10.0824 | 10.4076 | 0.0220 | 0.0047 | 0.0054 | 0.0045 | 1.0323 | 10.7433 |
| L30_04, L24_04 | 7.3076 | 4.3793 | 0.0217 | 0.0046 | 0.0042 | 0.0022 | 0.5993 | 2.6245 |
| L30_05, L24_05 | 2091.6717 | 456.1832 | 0.0243 | 0.0025 | 0.9560 | 0.1113 | 0.2181 | 99.4913 |
| L30_06, L24_06 | 15.3932 | 10.3354 | 0.0227 | 0.0035 | 0.0081 | 0.0053 | 0.6714 | 6.9394 |
| L30_07, L24_07 | 90.2712 | 42.7067 | 0.0231 | 0.0051 | 0.0439 | 0.0108 | 0.4731 | 20.2043 |
| L30_08, L24_08 | 16.0466 | 11.0566 | 0.0236 | 0.0062 | 0.0076 | 0.0037 | 0.6890 | 7.6183 |
| L30_09, L24_09 | 2500.9028 | 650.4438 | 0.0213 | 0.0024 | 1.4231 | 0.3041 | 0.2601 | 169.1698 |
| L30_10, L24_10 | 20.2369 | 10.6184 | 0.0204 | 0.0016 | 0.0119 | 0.0052 | 0.5247 | 5.5715 |
| L30_11, L24_11 | 23.9495 | 15.7281 | 0.0224 | 0.0052 | 0.0128 | 0.0091 | 0.6567 | 10.3290 |
| L30_12, L24_12 | 21.6502 | 11.1351 | 0.0249 | 0.0071 | 0.0096 | 0.0027 | 0.5143 | 5.7269 |
| L30_13, L24_13 | 1506.6826 | 1023.9988 | 0.0212 | 0.0055 | 0.9661 | 0.6575 | 0.6796 | 695.9484 |
| L30_14, L24_14 | 33.2031 | 15.4738 | 0.0141 | 0.0028 | 0.0439 | 0.0274 | 0.4660 | 7.2113 |
| L30_15, L24_15 | 59.2506 | 29.5681 | 0.0141 | 0.0030 | 0.0767 | 0.0475 | 0.4990 | 14.7555 |
| L30_16, L24_16 | 61.9032 | 28.0895 | 0.0139 | 0.0027 | 0.0795 | 0.0312 | 0.4538 | 12.7460 |
| L30_17, L24_17 | 4826.0754 | 1585.4326 | 0.0198 | 0.0027 | 3.1235 | 0.8992 | 0.3285 | 520.8366 |
| L30_18, L24_18 | 55.4137 | 28.0574 | 0.0220 | 0.0025 | 0.0304 | 0.0149 | 0.5063 | 14.2062 |
| L30_19, L24_19 | 53.4937 | 21.9909 | 0.0223 | 0.0036 | 0.0291 | 0.0126 | 0.4111 | 9.0403 |
| L30_20, L24_20 | 356.8270 | 263.2260 | 0.0188 | 0.0047 | 0.2223 | 0.0812 | 0.7377 | 194.1780 |
| L30_21, L24_21 | 4750.7889 | 1159.7467 | 0.0226 | 0.0025 | 2.5221 | 0.8266 | 0.2441 | 283.1135 |
| L30_22, L24_22 | 221.6685 | 97.8361 | 0.0119 | 0.0019 | 0.4308 | 0.2832 | 0.4414 | 43.1812 |
| L30_23, L24_23 | 48.5927 | 16.7517 | 0.0222 | 0.0037 | 0.0280 | 0.0134 | 0.3447 | 5.7749 |
| L30_24, L24_24 | 162.7046 | 125.9967 | 0.0218 | 0.0036 | 0.0840 | 0.0485 | 0.7744 | 97.5705 |
| L30_25,L6_01 | 54.4640 | 17.5995 | 0.0236 | 0.0031 | 0.0265 | 0.0076 | 0.3231 | 5.6871 |
| L30_26,L6_02 | 470.6592 | 112.7740 | 0.0252 | 0.0029 | 0.2067 | 0.0496 | 0.2396 | 27.0216 |
| L30_27,L6_03 | 495.9806 | 124.7559 | 0.0232 | 0.0026 | 0.2431 | 0.0372 | 0.2515 | 31.3803 |
| L30_28,L6_04 | 793.8516 | 603.9667 | 0.0163 | 0.0031 | 0.7066 | 0.3537 | 0.7608 | 459.5012 |
| L30_29,L6_05 | 2836.9857 | 898.8222 | 0.0224 | 0.0033 | 1.4842 | 0.2921 | 0.3168 | 284.7675 |
| L30_30,L6_06 | 1226.0209 | 220.1643 | 0.0207 | 0.0034 | 0.7700 | 0.2288 | 0.1796 | 39.5363 |
| L5_1 | 1924.3636 | 400.4894 | 0.0159 | 0.0027 | 1.9301 | 0.4631 | 0.2081 | 83.3480 |
| L5_2 | 177.1840 | 19.3230 | 0.0121 | 0.0029 | 0.3488 | 0.1769 | 0.1091 | 2.1073 |
| L5_3 | 836.4979 | 553.1577 | 0.0197 | 0.0019 | 0.5569 | 0.4063 | 0.6613 | 365.7910 |
| L5_4 | 101.5488 | 21.8780 | 0.0133 | 0.0024 | 0.1622 | 0.0818 | 0.2154 | 4.7135 |
| L5_5 | 48.3428 | 22.3635 | 0.0167 | 0.0021 | 0.0411 | 0.0157 | 0.4626 | 10.3454 |

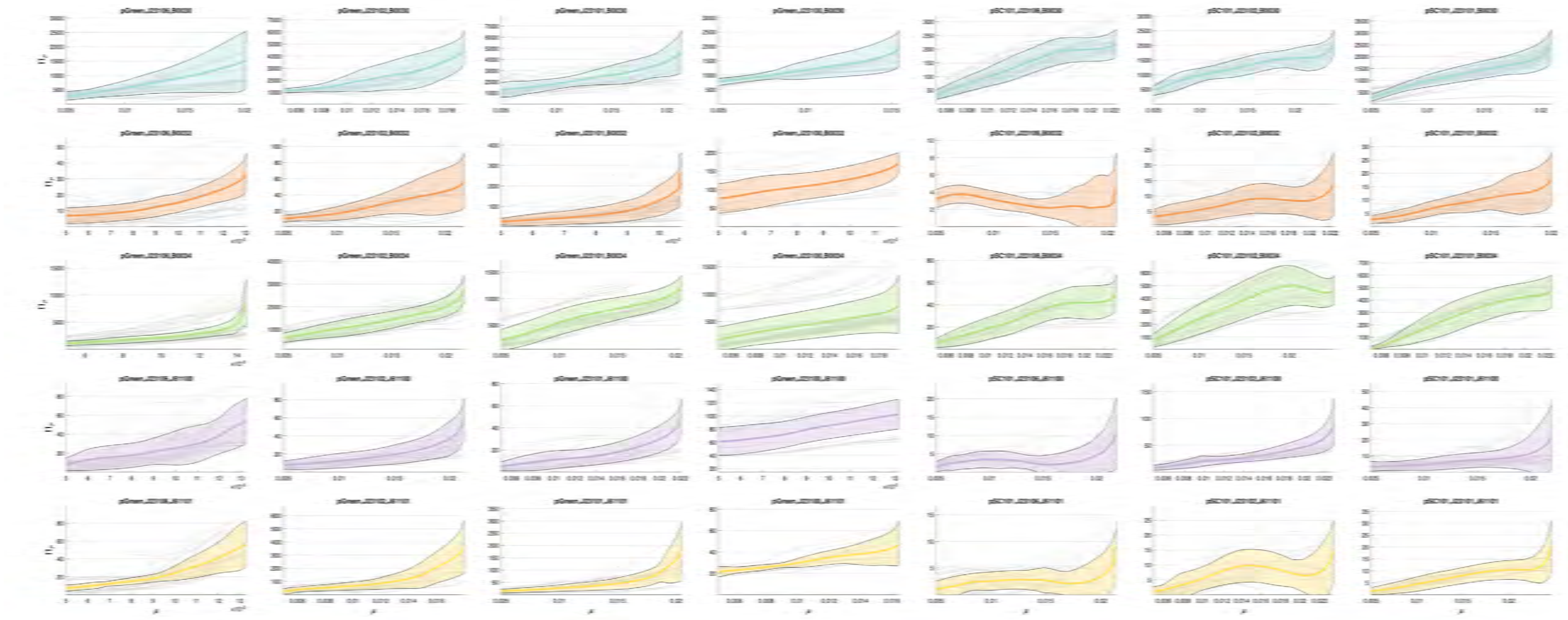

**Figure S2.2:** Experimental synthesis rate $(\mathrm{MEFL} \cdot \mathrm{Particle}^{-1} \cdot \mathrm{min}^{-1})$ as a function of specific growth rate $(\mathrm{min}^{-1})$ for the transcriptional units in libraries $\mathcal{L}_{30} \cup \mathcal{L}_5$. The grey lines correspond to the individual experimental replicas, the colored region corresponds to the $\pm\mathrm{std}$ one, and the thick line is the average synthesis rate.

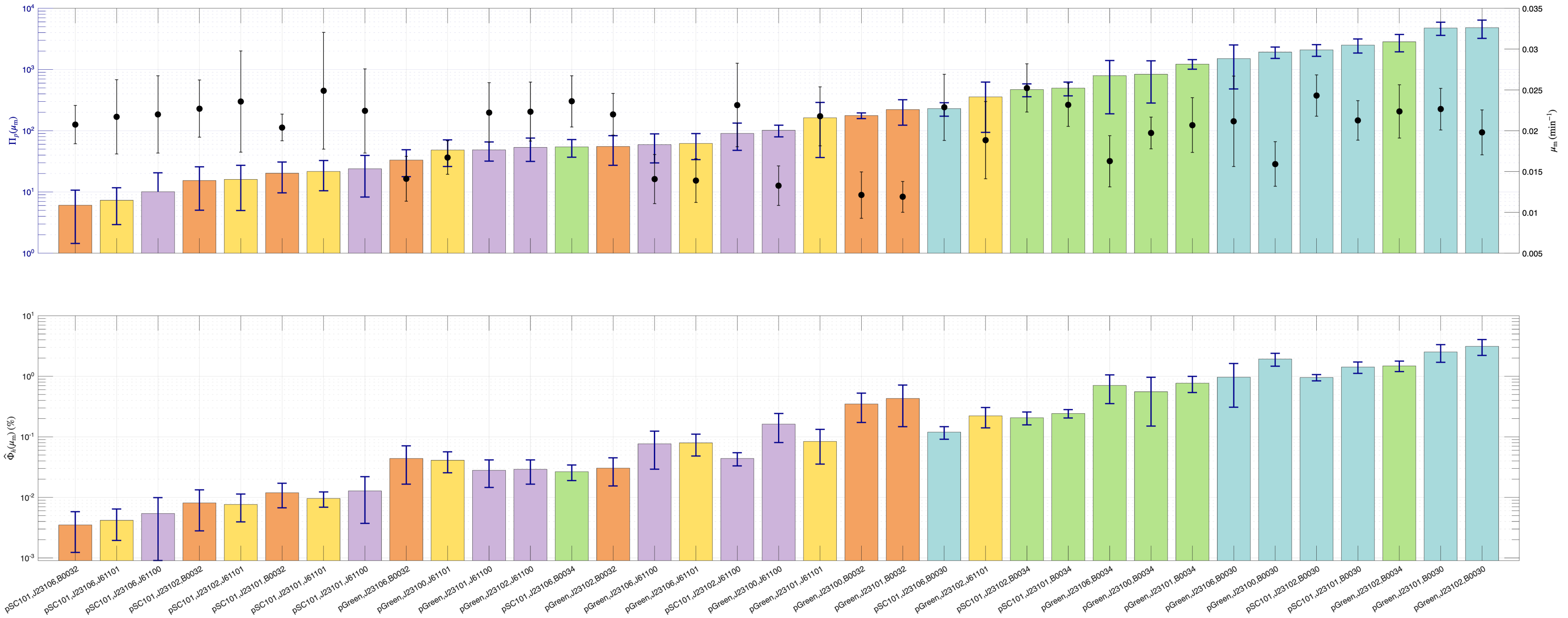

**Figure S2.3:** (Top:) Ordered experimental synthesis rates $(\mathrm{MEFL} \cdot \mathrm{Particle}^{-1} \cdot \mathrm{min}^{-1})$ evaluated at the maximum specific growth rate $(\mathrm{min}^{-1})$ for the transcriptional units in libraries $\mathcal{L}_{30} \cup \mathcal{L}_5$. The black dots show the corresponding maximum growth rate for each TU. Standard deviations are shown in both cases. (Bottom:) Cell burden evaluated as the estimation of the percentage of ribosomes allocated to the expression of each transcriptional unit.

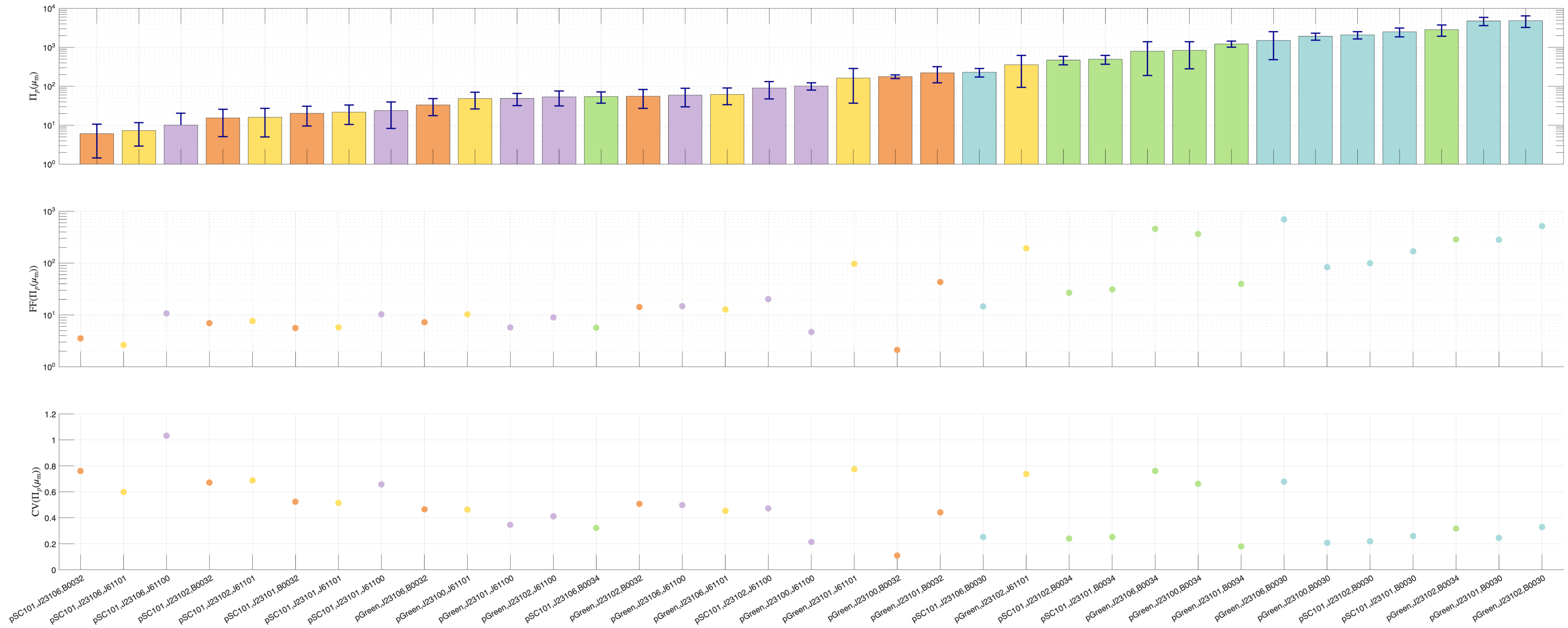

**Figure S2.4:** (Top:) Ordered experimental synthesis rates $\Pi\,(\mathrm{MEFL}\cdot\mathrm{Particle}^{-1}\cdot\mathrm{min}^{-1})$ evaluated at the maximum specific growth rate $\mu_{\max}\,(\mathrm{min}^{-1})$ for the transcriptional units in libraries $\mathcal{L}_{30}\cup\,\mathcal{L}_5$. The black dots show the corresponding maximum growth rate for each TU. Standard deviations are shown in both cases. (Bottom:) Fano Factor for the synthesis rate evaluated at $\mu_{\max}$. (Bottom:) Coefficient of Variation for the synthesis rate evaluated at $\mu_{\max}$.

### S3. *E. coli* digital twin for context-aware synthesis rate prediction

*S3.1. Resources-aware model of constitutive gene expression*

We consider transcriptional units that constitutively express a given protein $A$. Assuming the degradation rate of the protein can be neglected compared to the cell growth rate, the dynamics for the number of copies of the protein are obtained from the balance between the synthesis rate and the dilution rate caused by cell growth:

$$\dot{p}_A = \Pi_{p,A} - \mu p_A \tag{7}$$

where $p_A$ is the number of protein molecules $A$ per cell, $\mu(t)$ is the cell specific growth rate, and $\Pi_{p,A}$ denotes the synthesis rate.

Following results previously obtained in Santos-Navarro et al. (2021), we express the synthesis rate as

$$\Pi_{p,A} = \nu(s_i)\frac{1}{l_A}N_A J_A(s_i,\mu,r)r \tag{8}$$

where

$$\nu(s_i) = \nu_{\max}\frac{s_i}{K_{si}+s_i} \tag{9}$$

is the rate of peptide elongation attainable for a given amount of intracellular substrate available $s_i$, $l_A$ is the length of the protein being expressed (in a.a.), $N_A$ the gene copy number, $r$ the number of free ribosomes in the cell, and $J_A(s_i,\mu,r)$ is the *recruitment strength* of resources, a dimensionless functional coefficient that determines the distribution of resources between the host cell and the genes of interest and the relationship between the use of resources and cell growth (see Santos-Navarro et al. (2021) for details).

The recruitment strength of resources $J_A$ can be expressed as:

$$J_A(s_i,\mu,r) = E_{mA}\frac{\omega_A}{d_{mA}}\frac{K_A^{RBS}(s_i)}{1+\frac{K_A^{RBS}(s_i)}{d_{mA}}\mu r} \tag{10}$$

with $E_{mA} = 0.62\frac{l_A}{l_e}$ accounting for polysomal translation via the ribosomal average density $1/l_e$, $\omega_A$ is the transcription rate, $d_{mA}$ the transcript degradation rate, and

$$K_A^{RBS}(s_i) = \frac{k_A^b}{k_A^u + \frac{\nu(s_i)}{l_e}} \tag{11}$$

the substrate dependent effective RBS strength, which depends on the specific rate of intracellular substrate processing in the RBS (see Santos-Navarro et al. (2021) for details) and the ribosomes-RBS association and dissociation rate constants $k_A^b, k_A^u$.

For *E. coli*, we obtained a phenomenological relationship between the intracellular substrate $s_i$ and the extracellular one (see Supplementary Section S3.5) so that:

$$\nu(s) = \nu_{\max}\frac{s}{0.2081K_s+s} \tag{12}$$

where $\nu_{\max} = 20.5156\,(\mathrm{aa}\cdot\mathrm{sec}^{-1})$ is the maximum peptide elongation rate, that we assumed organism dependent and independent of the sequence of nucleotides. The constant $K_s$ is the substrate-dependent affinity constant for the extracellular Monod growth kinetics. To become independent of the substrate affinity, we defined the normalized substrate $s_n \triangleq \frac{s}{K_s}$ obtaining:

$$\nu(s_n) = \nu_{\max}f(s_n) \tag{13}$$

where $0 \le f(s_n) \le 1$. In the sequel, we used (13) to consider the peptide elongation rate as a function of the normalized extracellular substrate.

Using (13), the effective RBS strength for saturated substrate can be expressed as

$$\bar{K}_A^{RBS} = K_A^{RBS}(f(s_n)=1) = \frac{k_A^b}{k_A^u + \frac{\nu_{\max}}{l_e}} \tag{14}$$

Next we defined the nominal RBS strength as the inverse of the equilibrium dissociation constant of the binding reaction between the ribosomes and the RBS:

$$K_A^0 = \frac{k_A^b}{k_A^u} \tag{15}$$

and the RBS sensitivity as

$$\sigma_A^0 = \frac{K_A^0 - \bar{K}_A^{RBS}}{\bar{K}_A^{RBS}} = \frac{\nu_{\max}}{l_e}\frac{1}{k_A^u} \tag{16}$$

Using (15) and (16), the effective substrate-dependent RBS strength can be expressed as a function of the nominal one and the sensitivity:

$$K_A^{RBS}(s_n) = \frac{K_A^0}{1+\sigma_A^0 f(s_n)} \tag{17}$$

Finally, from (10) and (17), the synthesis rate (8) can be expressed as:

$$\Pi_{p,A} = N_A \frac{\omega_A}{d_{mA}} K_A^t(s_n,\varphi,\mu) \tag{18}$$

where the effective translation rate

$$K_A^t(s_n,\varphi,\mu) = \frac{E_{mA}}{l_A}\frac{K_A^{RBS}(s_n)r}{1+\frac{K_A^{RBS}(s_n)}{d_{mA}}\varphi}\nu(s_n) \tag{19}$$

depends on the RBS nominal strength and sensitivity, the available substrate and the interaction with the cell host via the flux of free ribosomes $\varphi = \mu r$.

The effective translation rate can be expressed as an explicit function of the nominal RBS strength and sensitivity as:

$$K_A^t(s_n,\varphi,\mu) = \nu_{\max}\frac{E_{mA}}{l_A}\frac{\frac{K_A^0}{\sigma_A^0}f(s_n)r}{\frac{1}{\sigma_A^0}+f(s_n)+\frac{K_A^0}{\sigma_A^0}\frac{\varphi}{d_{mA}}} \tag{20}$$

We define the ratio $\kappa^0 = K^0/\sigma^0$ as the *intrinsic resource-normalized translation initiation capacity* of an RBS. This parameter captures the inherent ability of an RBS to recruit ribosomes, independently of host growth state, while explicitly accounting for its sensitivity to translational demand. Throughout the manuscript, we refer to $\kappa$ as the *intrinsic initiation capacity* (IIC).

The parameter $\sigma^0$ quantifies the sensitivity of translation initiation to translational demand and cellular physiological state, while its inverse $\rho^0 = 1/\sigma^0$ provides a measure of robustness to translational load. Thus, RBSs with larger robustness factor (RF) $\rho^0$ values maintain a more stable effective initiation rate across different growth rates and resource availability, while RBSs with smaller $\rho^0$ exhibit stronger context dependence.

Using the definitions above, (20) becomes:

$$K_A^t(s_n,\varphi,\mu) = \nu_{\max}\frac{E_{mA}}{l_A}\frac{\kappa^0 f(s_n)r}{\rho^0+f(s_n)+\kappa^0\frac{\varphi}{d_{mA}}} \tag{21}$$

We used expression (21) in the sequel.

Notice the model above can easily be extended to TF-based transcription units by simply considering the transcription rate $\omega_A(TF)$ as a function of the transcription factor using eg. Hill functions.

#### *S3.2. Host-cell endogenous model (HEM)*

From the expressions for protein synthesis described in Supplementary Section S3.1, we obtained the ones providing an estimation of the specific growth rate, number of free ribosomes and flux of free ribosomes for the host cell when only the cell endogenous protein expression genes are considered. To this end, we considered the effective substrate-dependent RBS strength $K_x^{RBS}(s_n)$ and the resources recruitment strengths $J_x(s_n,\varphi)$ for the average ribosomal and non-ribosomal protein expression genes of the host cell $x = \{r, nr\}$. The specific cell growth rate can be obtained (see Supplementary Section S3.6) as:

$$\mu(s_n) = \frac{m_{\rm aa}}{m_{\rm rib}}\Phi_m \frac{N_r J_r}{1+\Sigma_{w,h}(s_n,\varphi)}\nu(s_n) \tag{22}$$

where $m_{\rm aa}$ is the average mass of amino acids, $m_{\rm rib} = m_{\rm aa}N_r l_r$ is the average protein mass of a ribosome, $\Phi_m$ is the fraction of mature ribosomes, and

$$\Sigma_{w,h}(s_n,\varphi) = \left(1+\frac{1}{E_{mr}}\right)N_r J_r(s_n,\varphi) + \left(1+\frac{1}{E_{mnr}}\right)N_{nr}J_{nr}(s_n,\varphi) \tag{23}$$

where $N_r, N_{nr}$ are the number of ribosomal and non-ribosomal protein expression genes active at any given instant (see (Santos-Navarro et al., 2021) for details).

The flux of free ribosomes $\varphi = \mu r$ can be obtained (see Supplementary Section S3.6) using:

$$\varphi = \frac{m_{\text{aa}}}{m_{\text{rib}}^2}\Phi_m^2 \left(\frac{N_r J_r}{1+\Sigma_{w,h}(s_n,\varphi)}\right)^2 \frac{m_h(\mu)}{N_r J_r + N_{nr} J_{nr}}\nu(s_n) \tag{24}$$

Notice equation (24) is in implicit form, as the flux of free ribosomes depends on the resources recruitment strengths $J_x(s_n, \varphi)$, which in turn depend on the flux of free ribosomes.

We updated the estimation of the protein mass content of the host wild-type *E. coli* cell $(m_h(\mu))$ as a function of its growth rate, as shown in Supplementary Section S3.7.

Next, we used data from (Bremer and Dennis, 2008) and (Santos-Navarro et al., 2021) to update the parameters of the host cell model. We kept the parameters described in Table S3 as in (Santos-Navarro et al., 2021), and estimated the parameter values for the endogenous model shown in Table S4. To this end, we ran 200 optimizations. The values shown correspond to the averages and deviations of the best 10% runs. The estimated values kept the same qualitative relationships already detected in (Santos-Navarro et al., 2021) and essentially are within the same orders of magnitude. From these parameters, we obtained the association and dissociation rate constants between RBSs and ribosomes shown in Table S5. The model accurately reproduces ribosomal mass fractions and ribosome counts as functions of cell growth rate (see Figure S3.3).

**Table S3:** General model parameters for the *E. coli* HEM. Sources: Santos-Navarro et al. (2021),Bremer and Dennis (2008)

| Parameter | Description | Value | Units |
|---|---|---|---|
| $\nu_{\max}$ | Maximum peptide elongation rate | $1320$ | $\text{aa} \cdot \text{min}^{-1}$ |
| $m_{\text{aa}}$ | average amino acid mass | $182.6 \cdot 10^{-24}$ | g |
| $N_r$ | Number of active ribosomal genes | 57 | adim |
| $N_{nr}$ | Number of active non-ribosomal genes | 1735 | adim |
| $l_{nr}$ | average non-ribosomal protein length | 333 | aa |
| $l_r$ | average ribosomal protein length | 195 | aa |
| $l_{e,r}$ | ribosome occupancy length (ribosomal transcripts) | 24 | aa |
| $l_{e,nr}$ | ribosome occupancy length (non ribosomal transcripts) | 25 | aa |
| $d_{mr}$ | ribosomal mRNA degradation rate constant | 0.16 | $\text{min}^{-1}$ |
| $d_{mnr}$ | non-ribosomal mRNA degradation rate constant | 0.2 | $\text{min}^{-1}$ |

**Table S4:** Estimated parameters for the *E. coli* HEM.

| Parameter | Description | Mean | Units |
|---|---|---|---|
| $K_r^0$ | Nominal RBS strength | 0.050 | $\text{molec}^{-1}$ |
| $K_{nr}^0$ | Nominal RBS strength | 9.075 | $\text{molec}^{-1}$ |
| $\sigma_r^0$ | RBS sensitivity | 0.135 | adim |
| $\sigma_{nr}^0$ | RBS sensitivity | 69.255 | adim |
| $\omega_r$ | Transcription rate | 4.050 | $\text{molec} \cdot \text{min}^{-1}$ |
| $\omega_{nr}$ | Transcription rate | 0.090 | $\text{molec} \cdot \text{min}^{-1}$ |
| $\Phi_m$ | Mature ribosome fraction | 0.805 | adim |

**Table S5:** Association and dissociation rate constants between the RBS and the ribosomes for the *E. coli* HEM. The subindexes $\text{r}, \text{nr}$ stand for ribosomal and non-ribosomal protein expression genes respectively.

| Parameter | Description | Mean | Units |
|---|---|---|---|
| $k_r^b$ | association rate constant RBS-ribosome | 4.67 | $\text{molec}^{-1} \cdot \text{min}^{-1}$ |
| $k_r^u$ | dissociation rate constant RBS-ribosome | 21.24 | $\text{min}^{-1}$ |
| $k_{nr}^b$ | association rate constant RBS-ribosome | 4.50 | $\text{molec}^{-1} \cdot \text{min}^{-1}$ |
| $k_{nr}^u$ | dissociation rate constant RBS-ribosome | 1.33 | $\text{min}^{-1}$ |

#### *S3.3. Use of the E. coli digital twin to estimate cell growth and loading*

Exogenous gene expression affects host growth by diverting resources. In the presence of an exogenous TU, equation (23) becomes:

$$\Sigma_{w,s}(s_n, \varphi, \theta) = \Sigma_{w,h}(s_n, \varphi) + \theta, \tag{25}$$

where $\theta$ quantifies the resource load imposed by the TU.

For a constitutive TU:

$$\theta(s_n, \varphi, \Theta) = \left(1 + \frac{1}{E_{m,\text{exo}}}\right) N_{\text{exo}} J_{\text{exo}}(s_n, \varphi, \Theta), \tag{26}$$

where $\Theta = \{l_{\text{exo}}, N_{\text{exo}}, \omega_{\text{exo}}, \kappa^0_{\text{exo}}, \rho^0_{\text{exo}}\}$ defines the TU.

To evaluate the interaction host-circuit induced by an exogenous gene circuit, the Host Equivalent Model can be used in two complementary ways depending on whether experimental data are used or not.

When experimental data are not used, the HEM can be used to estimate the cell growth rate $\mu$ and the flux of free resources $\varphi$ corresponding to given extracellular substrate and exogenous loading. From these, the synthesis rate of the exogenous gene circuit can be obtained. The procedure is summarized in the algorithm 1 and Figure S3.1.

Instead, experimental measurements of the specific growth rate can be leveraged to decouple the interactions between the exogenous genes being expressed, the host cell, and the environmental conditions. In this model-in-the-loop approach, the HEM is used as a digital twin, feeding it with the experimental values of the cell growth rate obtained for a given exogenous transcriptional unit. By combining the HEM with the TU model, the digital twin yields consensus estimates of $\theta$ and $\varphi$, which are then used in (18) to predict the TU synthesis rate. Additionally, this approach constrains the range of substrate availability consistent with observed growth and resource fluxes. The procedure is summarized in the algorithm 2 and Figure S3.2.

**Algorithm 1** Obtaining the host-circuit interaction and circuit synthesis rate without use of experimental data.

**Require:** the HEM model $\theta = \text{HEM}(f(s_n), \varphi)$ that relates the host loading $\theta$ and the flux of resources $\varphi$ parametrized by the normalized extracellular substrate function $f(s_n)$. See equations (24),(25), and Figure S3.1(a).
**Require:** the set of parameters $\Theta = \{l_{\text{exo}}, N_{\text{exo}}, \omega_{\text{exo}}, \kappa^0_{\text{exo}}, \rho^0_{\text{exo}}\}$ that characterize the exogenous circuit.
**Require:** the cell loading induced by the exogenous circuit $\theta = \theta(s_n, \varphi, \Theta)$ parametrized as a function of the substrate $f(s_n)$ and flux of resources $\varphi$. See equation (26) and Figure S3.1(b).

Consider a set of values $\{0 \leq f(s_n) \leq 1\}$
**for** each value $f(s_n)$ **do**
  calculate the intersection between $\theta = \text{HEM}(s_n, \varphi)$ and $\theta = \theta(s_n, \varphi, \Theta)$ obtaining the host-circuit *consensus* pair $(\theta^*, \varphi^*)$. See Figure S3.1(c).
  use (22) to obtain the corresponding cell specific growth rate. See Figure S3.1(d).
  estimate the synthesis rate of the exogenous circuit using (18) and (19). See Figure S3.1(e).
**end for**

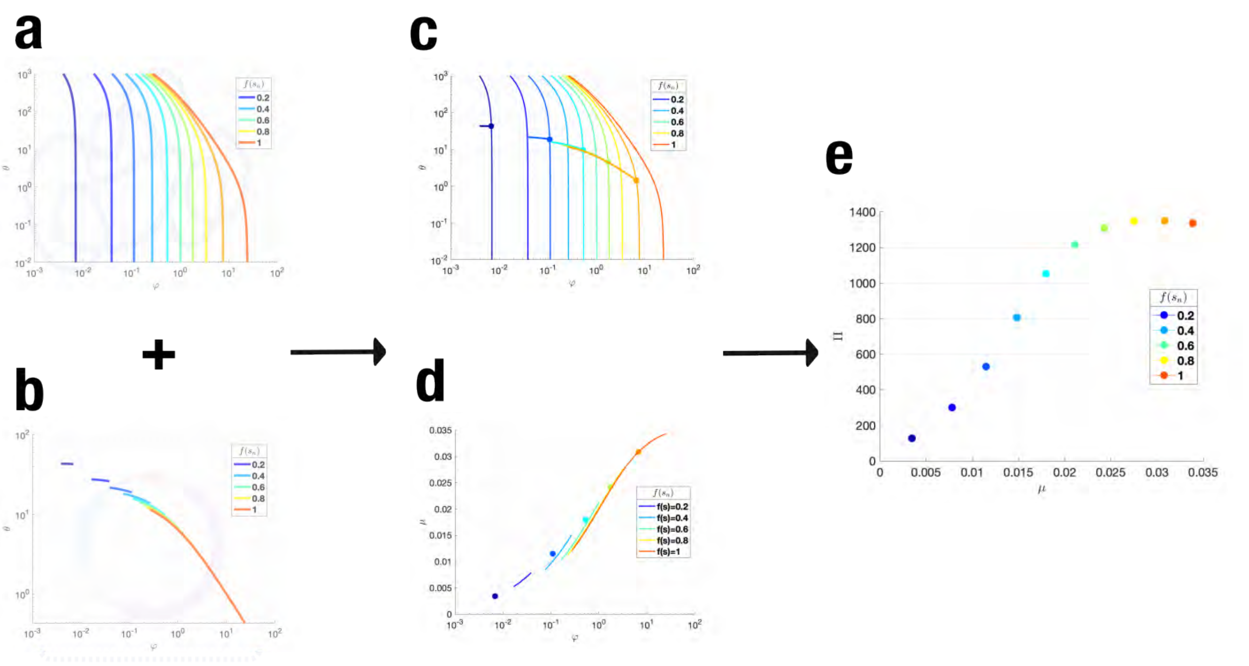

**Figure S3.1:** (a) Relationship $\theta = \text{HEM}(f(s_n), \varphi)$ between the load $\theta$ and the flux of resources $\varphi$ parametrized by the normalized substrate function $f(s_n)$ for the host cell. (b) Relationship $\theta = \theta(s_n, \varphi, \Theta)$ between the load $\theta$ and the flux of resources $\varphi$ parametrized by the normalized substrate function $f(s_n)$ for the exogenous circuit. (c) Interaction host-circuit and (d) resulting mapping between the growth rate and the flux of resources, parametrized by the normalized substrate function $f(s_n)$. (e) Predicted synthesis rate as a function of the cell growth rate and the corresponding values of $f(s_n)$.

**Algorithm 2** Leveraging the experimental growth rate to obtain the host-circuit interaction and circuit synthesis rate.

**Require:** the HEM model $\theta = \mathrm{HEM}(f(s_n), \varphi)$ that relates the host loading $\theta$ and the flux of resources $\varphi$ parametrized by the normalized extracellular substrate function $f(s_n)$. See equations (24), (25) and Figure S3.1(a).
**Require:** the set of parameters $\Theta = \{l_{\mathrm{exo}}, N_{\mathrm{exo}}, \omega_{\mathrm{exo}}, \kappa^0_{\mathrm{exo}}, \rho^0_{\mathrm{exo}}\}$ that characterize the exogenous circuit.
**Require:** the cell loading induced by the exogenous circuit $\theta = \theta(f(s_n), \varphi, \Theta)$ parametrized as a function of the substrate $f(s_n)$ and flux of resources $\varphi$. See equation (26) and Figure S3.1(b).
**Require:** the host-circuit mapping $\mu = g(\varphi, f(s_n))$ as obtained in Algorithm 2, Figure S3.1(d). See Figure S3.2(a, bottom).
**Require:** the experimental specific cell growth rate $\mu^*$.

1: using $\mu^* = g(\varphi, f(s_n))$, obtain the relationship $\varphi_c = h_c(\mu^*, f(s_n))$.
2: using $\varphi_c$ obtain the corresponding circuit load relationship $\theta_c = \theta(f(s_n), \varphi_c, \Theta)$
3: using $\theta_c$ obtain the corresponding HEM flux of resources $\varphi_h = h_h(f(s_n), \theta_c)$ from the relationship $\theta_c = \mathrm{HEM}(f(s_n), \varphi)$
4: obtain the intersection point $(\varphi^*, f^*(s_n))$ between the mappings $\varphi_c = h_c(\mu^*, f(s_n))$ and $\varphi_h = h_h(f(s_n), \theta_c)$. See Figure S3.2(a, top).
5: use $(\varphi^*, f^*(s_n))$ to estimate the synthesis rate of the exogenous circuit from (18) and (19). See Figure S3.2(b).

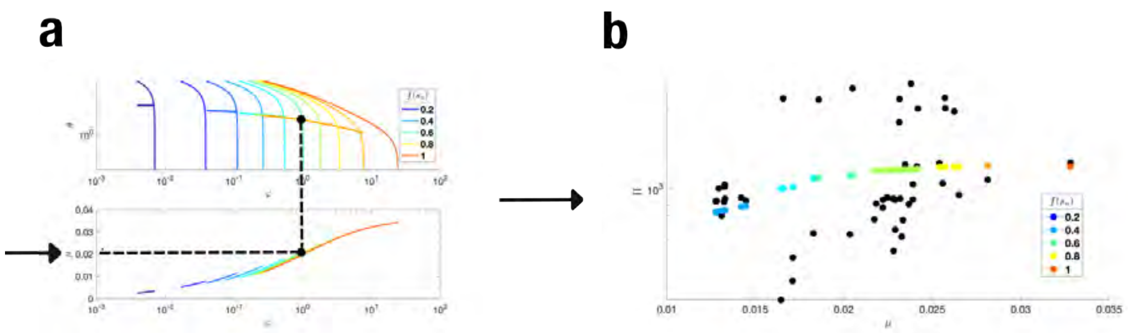

**Figure S3.2:** (a,top) Relationship between the load $\theta$, the flux of resources $\varphi$ and the specific growth rate $\mu$ for different amounts of normalized substrate for the host cell and a given TU circuit. The intersection between the host characteristic loading curve and the circuit one gives the resulting flux of free ribosomes in the strain, and the load induced by the exogenous circuit. (a, bottom) Relationship between the flux of resources and the specific growth rate resulting from the host-circuit interaction. The black dots correspond to the estimation of the resulting host-circuit *consensus* load $\theta$ and flux $\varphi$ when the experimental growth rate is used. (b) Predicted synthesis rate as a function of several experimental measurements of the cell growth rate (color dots). The color indicates the predicted value of the normalized substrate function $f(s_n)$. The black dots correspond to experimental values of synthesis rate and growth rate for the TU used as example.

*S3.4. Cell burden.*

Considering that the fraction of shared resources (ribosomes) captured by an exogenous construct is directly related to its global affinity for them (its resources recruitment strength), we defined the burden induced by the expression of an exogenous protein $A$ on the strain as the fraction of resources recruitment strength of the exogenous construct relative to the total sum of endogenous and exogenous resources recruitment strengths:

$$\Phi_{s,A} = \frac{N_A J_A(\mu, r)}{N_r J_r(\mu, r) + N_{nr} J_{nr}(\mu, r) + N_A J_A(\mu, r)} \tag{27}$$

Analogously, we defined the burden induced on the host as the fraction of exogenous resources recruitment strength relative to the endogenous ones:

$$\Phi_{h,A} = \frac{N_A J_A(\mu, r)}{N_r J_r(\mu, r) + N_{nr} J_{nr}(\mu, r)} \tag{28}$$

To estimate $\Phi_{s,A}$ we used the expression (29) for the mass dynamics of the exogenous protein A, taken from Santos-Navarro et al. (2021):

$$\dot{m}_A = \mu \left[ m_s(\mu) \frac{N_A J_A(\mu, r)}{N_r J_r(\mu, r) + N_{nr} J_{nr}(\mu, r) + N_A J_A(\mu, r)} - m_A \right] \tag{29}$$

where $m_s(\mu)$ is the protein mass of the strain $m_s(\mu) = m_h(\mu) + m_A$. To estimate the protein mass we used the approximation described in Supplementary Section S3.7.

Using (29) we estimated:

$$\Phi_{s,A} = \frac{\dot{m}_A + \mu m_A}{\mu m_s(\mu)} = \frac{m_{GFP} \Pi_{p,A}}{\mu m_s(\mu)} \tag{30}$$

where $m_{GFP} = 4.346 \cdot 10^{-5}$ (fg) is the protein mass per molecule of GFP.

From the definition of the protein mass of the strain and equation (29) evaluated at exponential steady state, we obtained the relationship:

$$\Phi_{s,A} m_s(\mu) = \Phi_{h,A} m_h(\mu) \tag{31}$$

from which we had:

$$\Phi_{h,A} = \frac{m_{GFP}\Pi_{p,A}}{\mu m_h(\mu)} \tag{32}$$

*S3.5. Peptide elongation rate as a function of extracellular substrate*

The peptide elongation rate can be obtained as the ratio between the total number of amino acids over the number of ribosomes actively translating times the specific growth rate of the cell. This provides the peptide elongation rate as a function of the cell growth rate:

$$\nu_t(\mu) = \frac{m_h(\mu)}{m_{\text{aa}}} \frac{\mu}{r_t} \tag{33}$$

where $m_h(\mu)$ is the protein mass of the host cell (see Supplementary Section S3.7), $m_{\text{aa}}$ the average amino acid mass and $r_t$ the number of active ribosomes. This is related to the number of total ribosomes in the cell as $r_t = \Phi_t \Phi_m r_T$ where $\Phi_m$ is the fraction of mature ribosomes with respect to the total number of ribosomes, and $\Phi_t$ is the fraction of mature ribosomes that are actively involved in the elongation of the polypeptide chains.

We estimated the number of active ribosomes as a function of of growth rate using data of the fraction of ribosomes and the peptide elongation rate as a function of growth rate available from Chure and Cremer (2023). Considering the mass fraction of ribosomes:

$$\Phi_R^T = \frac{m_{\text{aa}} r_T}{m_h(\mu)} \tag{34}$$

In addition, observing the data of the fraction of ribosomes in Chure and Cremer (2023) (see Figure S3.3 (Left)) we considered a phenomenological affine relationship between the fraction of ribosomes and the specific growth rate:

$$\Phi_R^T = a + b\mu \tag{35}$$

From (33)–(35) we obtain:

$$\nu_t(\mu) = \gamma_1 \frac{\mu}{\gamma_2 + \mu} \tag{36}$$

with

$$\begin{aligned} \gamma_1 &= \frac{N_r l_{p,r}}{b \Phi_t \Phi_m} \\ \gamma_1 &= \frac{a}{b} \end{aligned} \tag{37}$$

We estimated the parameters $a, b, \Phi_t\Phi_m$ using a bi-level optimization approach. First, for a range of values of the product $\Phi_t\Phi_m$, we obtained the parameters $a, b$ achieving the optimal fitting of the values $\Phi_R^C/\Phi_t\Phi_m$, where $\Phi_R^C$ are the values reported in Chure and Cremer (2023). From the resulting set of optimal solutions, we obtained the optimal value of the product $\Phi_t\Phi_m$ that best fitted the data of $\nu_t(\mu)$ also available from Chure and Cremer (2023). The optimal solutions we obtained were:

$$\begin{aligned} a &= 0.0660 \\ b &= 7.1502 \\ \Phi_t\Phi_m &= 0.72 \end{aligned} \tag{38}$$

From these, and the values of $N_r, l_{p,r}$ in Table S3, we obtained

$$\begin{aligned} \gamma_1 &= 25.9083\,(\text{aa} \cdot \text{sec}^{-1}) \\ \gamma_2 &= 0.0092\,(\text{min}^{-1}) \end{aligned} \tag{39}$$

Figure S3.3 shows the model prediction results. Notice the results are very similar to those obtained in Sechkar et al. (2024).

Considering the model (36), and taking into account that the cell growth rate depends on the extracellular substrate, $\mu = \mu(s)$, we assumed Monod kinetics for the specific growth rate of the cell as a function of the extracellular substrate:

$$\mu(s) = \frac{\mu_{\max} s}{K_s + s} \tag{40}$$

where the maximum specific growth rate $\mu_{\max}$ will depend on the host cell and its interaction with the exogenous genes, and $K_s$ is the substrate affinity constant.

Using (40) in (36), we obtained:

$$\nu_t(s) = \frac{\alpha s}{\beta + s} \tag{41}$$

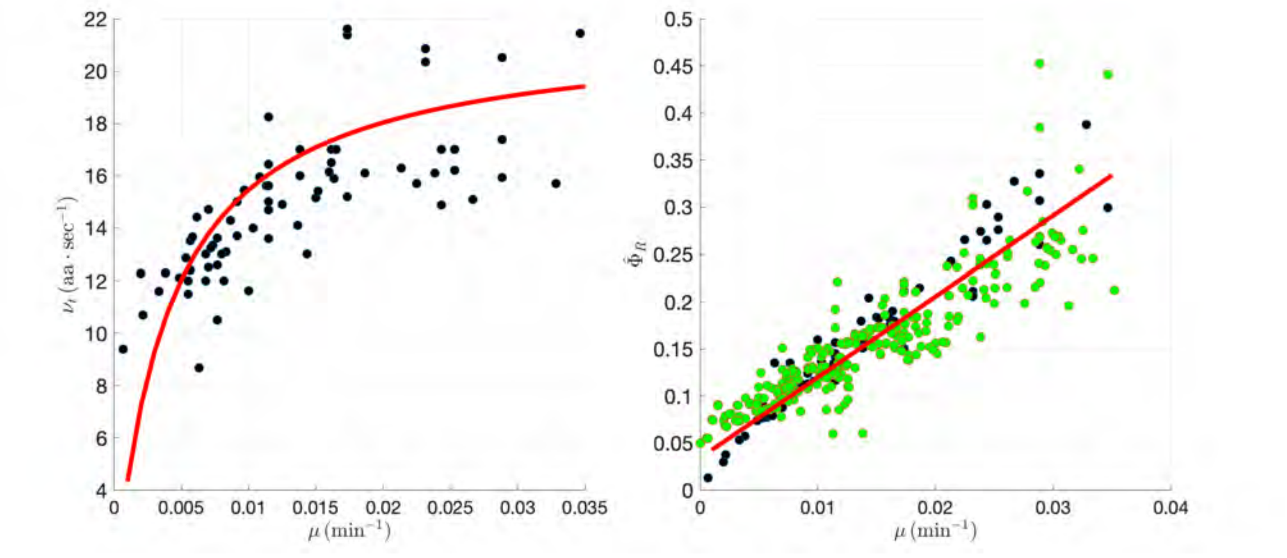

**Figure S3.3:** (Left:) Best fitting results for the relationship between the peptide elongation rate and the specific growth rate. (Right:) Estimation of the fraction of mature active ribosomes and comparison with the data from Bremer and Dennis (2008); Chure and Cremer (2023) scaled as $\Phi_R^C/\Phi_t\Phi_m$, with $\Phi_t\Phi_m = 0.72$

with

$$\begin{aligned} \alpha &= \frac{\gamma_1}{\gamma_2 + \mu_{\max}}\mu_{\max} \\ \beta &= \frac{\gamma_2}{\gamma_2 + \mu_{\max}} K_s \end{aligned} \tag{42}$$

Notice from (42) we could obtain

$$\begin{aligned} K_s &= \frac{\gamma_1 \beta}{\gamma_1 - \alpha} \\ \mu_{\max} &= \frac{\gamma_2 \alpha}{\gamma_1 - \alpha} \end{aligned} \tag{43}$$

Next, we considered the expression of the peptide elongation rate as a function of the intracellular substrate (an abstraction for the energy and amino acids required for translation):

$$\nu(s_i) = \nu_{\max}\frac{s_i}{K_{si} + s_i} \tag{44}$$

As a first approximation, we assumed that $\nu_{\max}$ is organism dependent but does not depend on the sequence of nucleotides of the gene being expressed.

To relate the intracellular and extracellular substrate, we resorted to the theoretical approaches to derive the Monod equation. Several alternatives exist Liu (2007); Zeng and Yang (2020); Ugalde-Salas et al. (2020). We followed a reasoning derived from the model developed in Weiße et al. (2015), where the quantity of intracellular substrate $s_i$ is related to the one of extracellular substrate $s$ through the dynamics of nutrient import and catabolism. Thus, from Santos-Navarro et al. (2021) we obtained the relationship between the intracellular and extracellular substrate:

$$s_i = \frac{\frac{k_m}{c-1}s}{\frac{k_t}{V_m}\frac{c}{c-1} + s} \tag{45}$$

where $V_m$ is the harvest volume Ugalde-Salas et al. (2020), $c = \frac{e_m v_m}{e_t v_t}$ with $e_t$ and $e_m$ transport and catabolism enzymes, and Michaelis-Menten kinetics are assumed (see Weiße et al. (2015); Santos-Navarro et al. (2021)) with rates $v_t$ and $v_m$. Notice if the maximum import and catabolism fluxes are balanced, ie. $c \approx 1$, then there is a linear relationship between the intracellular amount of substrate and its extracellular concentration. Otherwise, the intracellular amount of substrate $s_i$ saturates with increasing values of $s$.

We then took into account that the peptide elongation rate must give the same value independent of what argument is used to express it (intracellular or extracellular substrate or specific growth rate). Therefore, from (41) and (44):

$$\nu_{\max}\frac{s_i}{K_{si} + s_i} = \frac{\alpha s}{\beta + s} \tag{46}$$

Then, using (45) we obtained:

$$
\begin{aligned}
\alpha &= \nu_{\max}\frac{k_m}{K_{si}(c-1)+k_m}\\
\beta &= \frac{K_{si}k_t c}{V_m\left[K_{si}(c-1)+k_m\right]}
\end{aligned}
\tag{47}
$$

Under the hypothesis that the efficiency of nutrient import and catabolism are balanced, we assumed $c \approx 1$, so (47) becomes:

$$
\begin{aligned}
\alpha &= \nu_{\max}\\
\beta &= \frac{K_{si}k_t}{V_m k_m}
\end{aligned}
\tag{48}
$$

Note also that under the assumed condition $c = 1$, the relationship between the intracellular and extracellular substrate obtained from (45) becomes a linear one:

$$
s_i = V_m\frac{k_m}{k_t}s
$$

In our experimental data, we observed some samples grew at a maximum specific growth rate $\mu_{\max} = 0.035\,(\min^{-1})$ $(t_d = 19.8\,\min)$. Notice this is the maximum attainable specific growth rate for the wild-type host cell. Indeed, for a strain with an exogenous loading circuit, the maximum attainable specific growth rate will decrease as a function of loading (see Supplementary Sections S3.3 and S3.6). We took it as the maximum attainable specific growth rate. Then, from the results (39) and using (42) and (48) we obtained:

$$
\begin{aligned}
\nu_{\max} &= 20.5156\,(\mathrm{aa}\cdot\mathrm{sec}^{-1})\\
\beta &= 0.2081K_s\,(\mathrm{molec})
\end{aligned}
\tag{49}
$$

Note that the value obtained for $\nu_{\max}$ agrees with the maximum ones reported in the literature.

In summary, we obtained the phenomenological expressions relating the peptide elongation rate to the extracellular substrate and the specific growth rate for the host *E. coli*:

$$
\begin{aligned}
\nu(\mu) &= 20.5156\frac{1.2629\mu}{9.20\cdot 10^{-3}+\mu}\\
\nu(s) &= 20.5156\frac{s}{0.2081K_s+s}
\end{aligned}
\tag{50}
$$

Next we considered the normalized substrate $s_n \triangleq \frac{s}{K_s}$, so that:

$$
\nu(s_n) = 20.5156\frac{s_n}{0.2081+s_n}
\tag{51}
$$

As an alternative, defining $\mu_n \triangleq \frac{\mu}{\mu_{\max}} \subset [0,1]$ then:

$$
\nu(\mu_n) = 20.5156\frac{1.2629\mu_n}{0.2629+\mu_n}
\tag{52}
$$

Using (51) or (52) in place of (9) allows the expression of the substrate-dependent effective RBS strength (11) and the synthesis rate (8) as functions of the normalized extracellular substrate or, tantamount, the fraction of cell growth rate relative to its maximum one.

*S3.6. Derivation of the expressions for growth rate and flux of free ribosomes*

We started from the expression (53) in Santos-Navarro et al. (2021) evaluated under the condition of balanced efficiency between the transport of nutrients and their catabolism:

$$
\mu(s_n) = \frac{m_{aa}}{m_h(\mu)}\nu_{\max}\Phi_t^h r_a \nu(s_n)
\tag{53}
$$

where $\Phi_t^h$ is the fraction of ribosomes actively translating endogenous proteins (both ribosomal and non-ribosomal) at a given time instant, $r_a$ is the number of mature ribosomes available for protein translation (comprising the free ribosomes $r$ and the ones bound to translating complexes) , $\nu_{\max}$ is the maximum translation elongation rate, $m_{aa}$ the average amino acid mass, and $m_h(\mu)$ the growth-dependent protein mass content of the wild-type host cell (ie. without exogenous genes).

The fraction of ribosomes actively translating endogenous proteins can be expressed as a function of the ratio between the resources recruitment strengths of the endogenous genes and the total sum considering both endogenous and exogenous genes (see Supplementary Information in Santos-Navarro et al. (2021) for details):

$$\Phi_t^h = \frac{N_r J_r + N_{nr} Jnr}{1 + \Sigma_{w,h}(s_n, \varphi)} \tag{54}$$

with

$$\Sigma_{w,h}(s_n, \varphi) = \left(1 + \frac{1}{E_{mr}}\right) N_r J_r(s_n, \varphi) + \left(1 + \frac{1}{E_{mnr}}\right) N_{nr} J_{nr}(s_n, \varphi) \tag{55}$$

The number of mature ribosomes, $r_a$, can be expressed as a function of the free ones and the total loading in the cell defined by the total sum of resources recruitment strengths:

$$r_a = [1 + \Sigma_{w,h}(s_n, \varphi)]\, r \tag{56}$$

Using (54) and (56) in (53) we obtain equation:

$$\mu(s_n) = \frac{m_{aa}}{m_h(\mu)} [N_r J_r + N_{nr} J_{nr}]\, r \nu(s_n) \tag{57}$$

At exponential balanced growth the mass fractions and the resource recruitment strength ones are equivalent. Thus, for the ribosomal fraction:

$$\frac{m_{\text{rib}} r_T}{m_h(\mu)} = \frac{N_r J_r}{N_r J_r + N_{nr} J_{nr}} \tag{58}$$

where the total amount of ribosomes, $r_T$ can in turn be expressed as:

$$r_T = \frac{N_r J_r + N_{nr} J_{nr}}{\Phi_m} r \tag{59}$$

Using (59) in (58) we obtain:

$$\frac{1}{m_h(\mu)} = \frac{N_r J_r}{N_r J_r + N_{nr} J_{nr}} \frac{1}{m_{\text{rib}}} \frac{\Phi_m}{[1 + \Sigma_{w,h}(s_n, \varphi)]\, r} \tag{60}$$

that used in (57) results in

$$\mu(s_n) = \frac{m_{\text{aa}}}{m_{\text{rib}}} \Phi_m \frac{N_r J_r}{1 + \Sigma_{w,h}(s_n, \varphi)} \nu(s_n) \tag{61}$$

On the other hand, the number of free ribosomes can be obtained by equating the total number of ribosomes $r_T$ derived from (58) and (59), obtaining:

$$r = \Phi_m \frac{m_h(\mu)}{m_{\text{rib}}} \frac{N_r J_r}{N_r J_r + N_{nr} J_{nr}} \frac{1}{1 + \Sigma_{w,h}(s_n, \varphi)} \tag{62}$$

The flux of free ribosomes, defined as $\varphi \triangleq \mu r$, resulting from (61) and (62) is:

$$\varphi = \frac{m_{\text{aa}}}{m_{\text{rib}}^2} \Phi_m^2 \left( \frac{N_r J_r}{1 + \Sigma_{w,h}(s_n, \varphi)} \right)^2 \frac{m_h(\mu)}{N_r J_r + N_{nr} J_{nr}} \nu(s_n) \tag{63}$$

*S3.7. Estimation of the host protein mass and dry weight as a function of growth rate.*

In Santos-Navarro et al. (2021) we considered a model relating the cell dry mass and the protein mass with the specific growth rate. for the wild type *E. coli* cell. Different from the approach used there, here we used updated data from Bremer and Dennis (2008) relating growth rate, cell dry weight and protein mass content, and fitted a Hill-like model to approximate it.

Thus, we considered the model:

$$m_{cDW}(\mu) = \frac{c_0 + c_1 \mu^{c_3}}{1 + c_2 \mu^{c_3}} \tag{64}$$

for the cell dry weight, and an analogous one for the endogenous protein mass content:

$$m_h(\mu) = \frac{d_0 + d_1 \mu^{d_3}}{1 + d_2 \mu^{d_3}} \tag{65}$$

We used the experimental data reported in Bremer and Dennis (2008), shown in Table S6.

The best fits were obtained for the parameters shown in Table S7, and the corresponding results are shown in Figure S3.4.

**Table S6:** Experimental data relating cell and protein dry mass and specific growth rate for *E. coli*. Source: Bremer and Dennis (2008).

| | $t_d$ (min) | 100 | 60 | 40 | 30 | 24 | 20 |
|---|---|---|---|---|---|---|---|
| | $\mu$ (min$^{-1}$) | 0.0069 | 0.0116 | 0.0173 | 0.0231 | 0.0289 | 0.0347 |
| $m_h$ (fg) | new | 136 | 214 | 295 | 387 | 431 | 426 |
| $m_{cDW}$ (fg) | new | 226 | 374 | 555 | 774 | 921 | 1023 |

**Table S7:** Best fitting parameters for the phenomenological mass models. Values of parameters obtained so the masses are given as femtograms. *E. coli* cell dry weight as described by equation (64). *E. coli* cell protein mass as described by equation (65). The results were obtained from 10 runs of the fitting optimization.

| | | Estimated parameters | | | |
|---|---|---|---|---|---|
| $m_{cDW}$ (fg) | | $c_0$ | $c_1$ | $c_2$ | $c_3$ |
| | mean | $211.524$ | $214.880 \cdot 10^6$ | $182.632 \cdot 10^3$ | $3.120$ |
| | std | $1.90$ | $39.51$ | $34.73$ | $0.004$ |
| $m_h$ (fg) | | $d_0$ | $d_1$ | $d_2$ | $d_3$ |
| | mean | $115.603$ | $476.989 \cdot 10^6$ | $1076.9 \cdot 10^3$ | $3.356$ |
| | std | $0.19$ | $17.34$ | $39.70$ | $0.009$ |

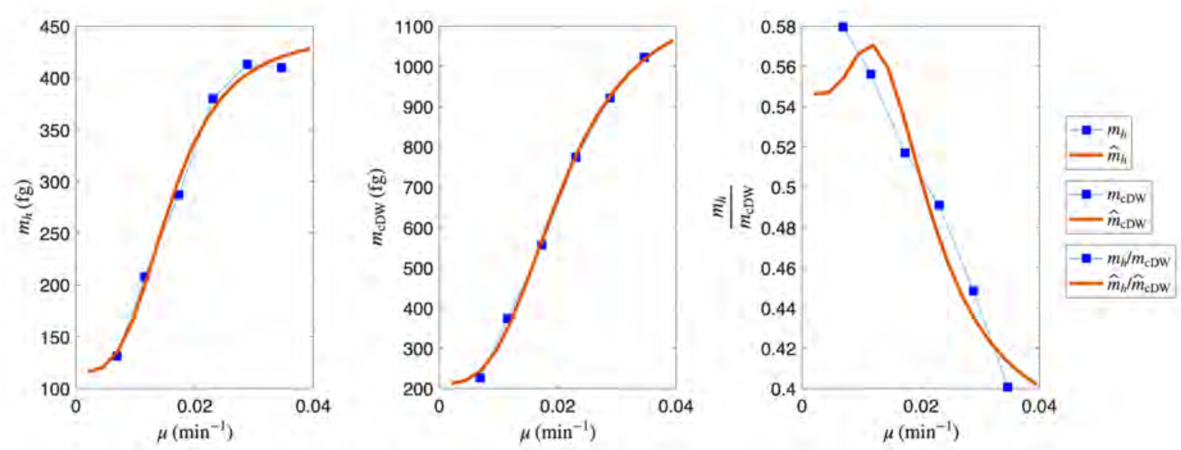

**Figure S3.4:** Fitting of the mass models (65) (left), (64) (center) and estimated ratio between the cell protein mass content and the cell dry weight (right)

### S4. Bioparts characterization. Optimization algorithms for parameter estimation.

#### *S4.1. Optimization algorithms*

In our study, we employed the Enhanced Scatter Search (eSS) method from the open-source toolboxes MEIGO Egea et al. (2014), and BADS Acerbi and Ma (2017) to address the challenging parameter estimation problems.

MEIGO provides a flexible platform for global optimization tailored to bioinformatics and systems biology applications, supporting black-box objective functions that encapsulate intricate simulations or nested optimizations. For our parameter estimation tasks (aimed at minimizing the discrepancy between model predictions and experimental data) we configured eSS with mostly default solver options, including a modest reference set size and standard iteration limits, which proved sufficient for robust convergence without extensive tuning. This approach allowed us to efficiently explore the high-dimensional, non-convex parameter space, yielding reliable estimates that captured key model dynamics. The eSS solver is a hybrid metaheuristic that uses local solvers to refine promising solutions generated by the global scatter search process, improving convergence and efficiency in challenging nonlinear optimization problems. In this strategy, local solvers (either gradient-based or direct search) are applied as an intensive local improvement step exclusively to the most promising solutions in a reference set and any new high-quality points entering it. This refinement occurs after each iteration, uses a merit-and-distance filter to avoid redundant searches on nearby points, making eSS a powerful global-local hybrid that combines broad exploration with targeted local convergence. The eSS method is also a population-based evolutionary metaheuristic that uses a small reference set of diverse solutions, generating new vectors via systematic combinations (unlike random crossovers in genetic algorithms). Enhancements include a $1{+}1$ replacement strategy (offspring replace only their generating parent to maintain diversity and avoid stagnation), a "go-beyond" mechanism to extend promising directions for deeper exploration, and memory heuristics to steer local searches from suboptimal traps. These enable effective balancing of global diversification and local intensification. Unlike computationally prohibitive exact methods or other heuristics (e.g., simulated annealing or particle swarm optimization), eSS excels on multimodal landscapes, outperforming other solvers in benchmarking studies from the domain of dynamic model calibration Villaverde et al. (2018); Penas et al. (2024).

On the other hand, BADS (Bayesian Adaptive Direct Search), is a hybrid algorithm combining fast Bayesian Optimization (BO) with systematic exploration, for efficient model fitting, especially for expensive black-box functions. BADS is computationally very efficient for small-medium size sets of estimated parameters for both deterministic and stochastic cost functions.

#### *S4.2. Cost function for parameter estimation.*

All parameter estimations were performed by minimising a weighted logarithmic error between predicted and experimental synthesis rates. Given the wide dynamic range of the measured $\Pi(\mu)$ values (spanning more than two orders of magnitude), we defined the cost function

$$= \frac{1}{N_{\mathrm{data}}} \sum_{i=1}^{N_{\mathrm{data}}} \Big| \log_{10}\big(\Pi_{\mathrm{pred}}^{(i)}\big) \log_{10}\big(\Pi_{\mathrm{exp}}^{(i)}\big) - \big(\log_{10}\big(\Pi_{\mathrm{exp}}^{(i)}\big)\big)^2 \Big| \tag{66}$$

with $N_{\mathrm{data}}$ the total number of datapoints (number of TUs $\times$ number of growth-rate values $\times$ number of replicates). Working in log–space distributes the contribution of low- and high-expression constructs more evenly than a linear least–squares objective that would over–weight high-expression constructs. The multiplicative factor $\log_{10}(\Pi_{\mathrm{exp}})$ introduces a mild weighting towards data points with larger expression, where relative errors matter. This formulation therefore balances sensitivity across experimental conditions in the presence of large expression differences among TUs.

### S5. Bioparts characterization. Identifiability analysis.

*S5.1. Structural identifiability analysis*

The effective translation rate (21) of each RBS depends on the composite parameters $\kappa^0 = \frac{K_A^0}{\sigma_A^0}$(RBS IIC) and $\rho^0 = \frac{1}{\sigma_A^0}$ (RBS RF). The logarithmic sensitivities of the effective translation rate with respect to the composite parameters:

$$\mathcal{S}_{K_A^t}^{\kappa^0} = \frac{1}{K_A^t(s_n, r)} \frac{\partial K_A^t(s_n, r)}{\partial \kappa^0} = \frac{\rho^0 + f(s_n)}{\kappa^0} \frac{1}{\rho^0 + f(s_n) + \kappa^0 \frac{\varphi}{d_{mA}}}$$
$$\mathcal{S}_{K_A^t}^{\rho^0} = \frac{1}{K_A^t(s_n, r)} \frac{\partial K_A^t(s_n i, r)}{\partial \rho^0} = \frac{-1}{\rho^0 + f(s_n) + \kappa^0 \frac{\varphi}{d_{mA}}}$$

satisfy the near-linear relation

$$\mathcal{S}_{K_A^t}^{\kappa^0} + \frac{\rho^0 + f(s_n)}{\kappa^0} \mathcal{S}_{K_A^t}^{\rho^0} = 0 \tag{67}$$

Moreover, we noticed that consistent estimated values of $\rho^0$ concentrate within a narrow physiological range $\rho^0 < 0.03$ across RBSs in the library (see Supplementary Sections S5.2 and S5.3). From the identity between the values of (51) and (52) in Supplementary Section S3.5, and considering that the minimum experimental specific growth rate of practical use for the parameters estimation is $\mu = 0.005\,(\mathrm{min}^{-1})$ and a maximum specific growth rate $\mu_m = 0.036\,(\mathrm{min}^{-1})$, we expected a minimum value $\left.\frac{\nu(s_n)}{\nu_{\max}}\right|_{\min} \approx 0.4 \gg \rho^0$. Therefore:

$$\mathcal{S}_{K_A^t}^{\kappa^0} + \frac{f(s_n)}{\kappa^0} \mathcal{S}_{K_A^t}^{\rho^0} = 0 \tag{68}$$

For a given experiment involving a TU with one RBS, the substrate function $f(s_n)$ varies little (in the typical range $[0.4, 0.9]$) along all experimental growth rates. This implies that only one effective combination of $(\kappa^0, \rho^0)$ is strongly identifiable from data containing only one RBS biopart, and that there exists a nearly one-dimensional sensitivity manifold in $(\kappa^0, \rho^0)$-space along which $K_A^t$ is essentially unchanged.

Notice that for parameters estimation we use measurements of the synthesis rate and the expression (18):

$$\Pi_{p,A} = N_A \frac{\omega_A}{d_{mA}} K_A^t(s_n, \varphi, \mu)$$

from which it is clear that the sensitivity of the synthesis rate with respect to the RBS composite translation parameters is the same as the one of the effective translation rate

$$\begin{aligned} \mathcal{S}_{\Pi_{p,A}}^{\kappa^0} &= \frac{1}{\Pi_{p,A}} \frac{\partial \Pi_{p,A}}{\partial \kappa^0} = \mathcal{S}_{K_A^t}^{\kappa^0} \\ \mathcal{S}_{\Pi_{p,A}}^{\rho^0} &= \frac{1}{\Pi_{p,A}} \frac{\partial \Pi_{p,A}}{\partial \rho^0} = \mathcal{S}_{K_A^t}^{\rho^0} \end{aligned} \tag{69}$$

On the other hand, for the copy number and the transcription rate, we have:

$$\begin{aligned} \mathcal{S}_{\Pi_{p,A}}^{\omega_A} &= \frac{1}{\Pi_{p,A}} \frac{\partial \Pi_{p,A}}{\partial \omega_A} = \frac{1}{\omega_A} \\ \mathcal{S}_{\Pi_{p,A}}^{N_A} &= \frac{1}{\Pi_{p,A}} \frac{\partial \Pi_{p,A}}{\partial N_A} = \frac{1}{N_A} \end{aligned} \tag{70}$$

For a single transcriptional unit, any two multiplicative factors (e.g. $\omega_A$ and $N_A$) are not separable from a purely structural identifiability standpoint, as only their product enters the expression for $\Pi_{p,A}$. However, in our combinatorial libraries each copy-number parameter $N_A$ is shared by many TUs (all constructs carrying the same origin), while different promoters contribute distinct $\omega_A$ values across those TUs. This breaks the degeneracy between copy number and transcription rate at the library level and leads to a full-rank sensitivity matrix in the $(N_A, \{\omega_A\})$ block. Moreover, although the effective translation model structurally exhibits a nearly one-dimensional sensitivity manifold in $(\kappa^0, \rho^0)$ for any single RBS, this limitation is mitigated at the library level in practice by the large combinatorial structure of our libraries. Each RBS appears in multiple TUs combined with different promoters and plasmid origins. Libraries $\mathcal{L}_{24}$, $\mathcal{L}_{30}$ provide enough independent information to constrain all promoter strengths, plasmid copy numbers, and RBS parameters, generating distinct host–circuit interactions and spanning a diverse range of growth states $(\mu)$. As a result, structural non-identifiability at the single-construct level does not propagate to the library level.

To illustrate the origin of the identifiability properties discussed above, Figure S5.1 shows the sparsity pattern of the global sensitivity matrix (Jacobian) for our full combinatorial libraries ( $\mathcal{L}_{30} = \mathcal{L}_{24} \cup \mathcal{L}_6$, and $\mathcal{L}_5$). Each TU activates only the parameters corresponding to its promoter, RBS and plasmid origin, while all other part parameters have identically zero sensitivity for that TU. Because different promoters, RBSs and origins appear across partially overlapping subsets of TUs, the resulting Jacobian is structurally full rank: each parameter column has a distinct activation pattern that cannot be replicated by any other column. This block-sparse activation pattern of the global Jacobian is what breaks the trivial multiplicative degeneracies (e.g. between copy number and transcription rate) and enables robust identification of promoter strengths, RBS translation parameters and plasmid copy numbers at the library scale.

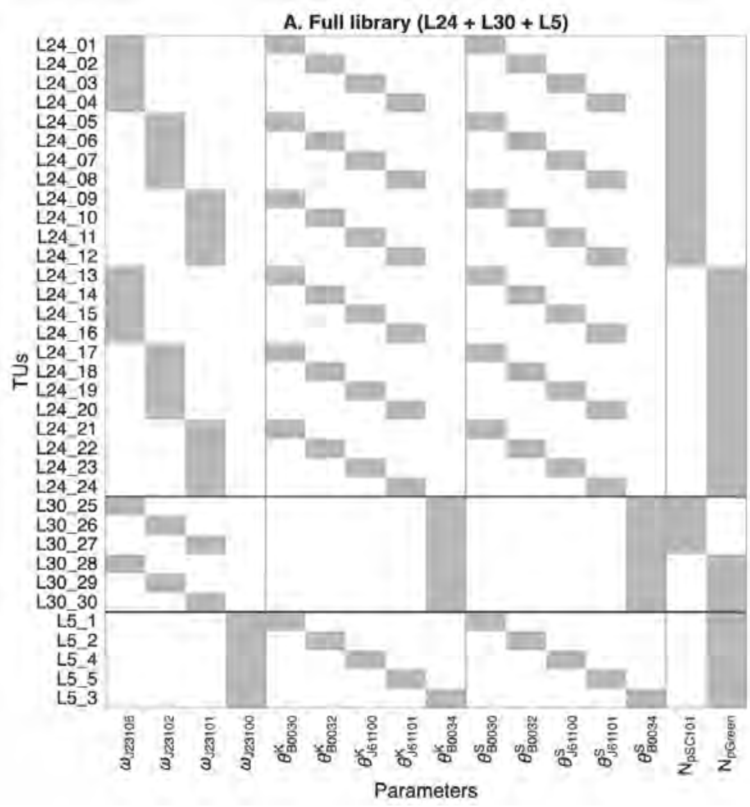

**Figure S5.1: Structure of the global Jacobian and effect of collapsing to a single-promoter sublibrary.** Binary activation pattern of the sensitivity/Jacobian matrix for the full combinatorial libraries: $\mathcal{L}_{30} = \mathcal{L}_{24} \cup \mathcal{L}_6$(B0034 only), and $\mathcal{L}_5$ (J23100 only). Grey = active, White = inactive. Rows correspond to transcriptional units (TUs) and columns correspond to part parameters, grouped by parameter type: promoter strengths ($\omega_p$), RBS translation parameters ($\theta^p_K = \kappa^0_p$ and $\theta^p_\sigma = \rho^0_p$), and copy numbers ($N_{\mathrm{ori,p}}$). Each TU activates only the parameters of the promoter, RBS and plasmid origin it contains, resulting in a sparse but structurally full-rank Jacobian in the libraries $\mathcal{L}_{24}$ and $\mathcal{L}_{30}$ , where each parameter exhibits a distinct and partially overlapping activation pattern across TUs. This combinatorial excitation is what resolves the multiplicative degeneracies (e.g. between copy number and transcription rate) that arise in single constructs, and it is essential for the robust identification of context-independent part parameters. In contrast, restricting the analysis to the B0034-only sublibrary ($\mathcal{L}_6$) collapses the Jacobian to a much poorer structure in which all TUs share the same RBS and differ only in the promoter and plasmid origin blocks. Because the two composite translation parameters of each RBS span a nearly one-dimensional sensitivity manifold, the reduced Jacobian becomes rank-deficient. Similarly, restricting the analysis to the J23100-only sublibrary ($\mathcal{L}_5$) also collapses the Jacobian to a much poorer structure in which all TUs share the same promoter and plasmid origin and differ only in the RBS block. In this case, the reduced Jacobian becomes close to rank-deficient, explaining why residual translational variability is reabsorbed into $\omega_{J23100}$ and why LOOCV estimates exhibit multi-modality in this subset.

#### *S5.2. Initial bioparts characterization using Library $\mathcal{L}_{30}$.*

This section summarizes the initial characterization of all bioparts using the full combinatorial library $\mathcal{L}_{30}$ and the complete translation model. Table S8 reports the estimated plasmid copy numbers, promoter transcription rates, and RBS translational parameters obtained from both full-library fits and leave-one-out cross-validation (LOOCV). The accompanying figures illustrate the corresponding parameter distributions and the relationships between the composite RBS parameters $(\kappa_0, \rho_0)$ and the original parameterization $(K_0, \sigma_0)$. These results provide an empirical view of the practical identifiability properties discussed in Supplementary Section S5.3 and are the basis for fixing the robustness factor $\rho_0$ in the approximate translation model used in subsequent analyses.

#### *S5.3. Practical identifiability and constrained parameter manifolds*

In addition to the structural identifiability analysis presented above, we performed a practical identifiability assessment based directly on the experimental data and the digital-twin model. For each TU and each experimental growth rate $\mu$, we computed the sensitivities of the synthesis rate with respect to every biopart parameter using the analytical expressions (69)-(70), evaluated at the experimental $\Pi_A$ and at the parameter estimates. Stacking

**Table S8: Estimated characterization of the bioparts in library $\mathcal{L}_{30}$ using the complete model of translation.** For the estimations using all the TUs in the library (Full Library Estimates) 60 estimation runs were carried out. For the leave-one-out cross-validation estimates (LOOCV Library Estimates) 10 estimation runs were carried out for each one of the 30 LOOCV cases. In both cases (Full and LOOCV) estimates we carried out considering the predicted and averaged synthesis rates for each experimental instance (group of 10 replica wells) of each of the 30 TUs. Units $N_A$: $\mathrm{adim}$. The gene copy number $N_A = 5$ for the low-copy plasmid pSC101 was known beforehand from Thompson et al. (2018). Units $\omega_A$: $\mathrm{molec} \cdot \mathrm{min}^{-1}$. Units $\kappa_A^0$: $\mathrm{molec}^{-1}$, Units $\rho_A^0$: $\mathrm{adim}$

**Full Library Estimates**

| | Plasmids | | Promoters | | |
|---|---|---|---|---|---|
| | $N_A$ | | $\omega_A$ | | |
| **Biopart** | **pSC101** | **pGreen** | **J23106** | **J23102** | **J23101** |
| Mean | 5 | 18.128 | 0.0501 | 0.1953 | 0.1530 |
| Std | 0 | 0.293 | 0.0018 | 0.0059 | 0.0050 |

| | Ribosome Binding Sites | | | | |
|---|---|---|---|---|---|
| | $\kappa_A^0 = K_A^0/\sigma_A^0$ | | | | |
| **Biopart** | **B0030** | **B0032** | **J61100** | **J61101** | **B0034** |
| Mean | 0.8667 | 0.0073 | 0.0060 | 0.0091 | 0.1811 |
| Std | 0.0538 | 0.0003 | 0.0002 | 0.0003 | 0.0083 |
| | $\rho_A^0 = 1/\sigma_A^0$ | | | | |
| **Biopart** | **B0030** | **B0032** | **J61100** | **J61101** | **B0034** |
| Mean | 0.0221 | 0.0060 | 0.0078 | 0.0077 | 0.0065 |
| Std | 0.0139 | 0.0025 | 0.0063 | 0.0061 | 0.0042 |

**LOOCV Library Estimates**

| | Plasmids | | Promoters | | |
|---|---|---|---|---|---|
| | $N_A$ | | $\omega_A$ | | |
| **Biopart** | **pSC101** | **pGreen** | **J23106** | **J23102** | **J23101** |
| Mean | 5 | 18.071 | 0.0513 | 0.1978 | 0.1551 |
| Std | 0 | 1.632 | 0.0052 | 0.0156 | 0.0092 |

| | Ribosome Binding Sites | | | | |
|---|---|---|---|---|---|
| | $\kappa_A^0 = K_A^0/\sigma_A^0$ | | | | |
| **Biopart** | **B0030** | **B0032** | **J61100** | **J61101** | **B0034** |
| Mean | 0.8698 | 0.0074 | 0.0060 | 0.0092 | 0.1770 |
| Std | 0.1170 | 0.0010 | 0.0009 | 0.0013 | 0.0144 |
| | $\rho_A^0 = 1/\sigma_A^0$ | | | | |
| **Biopart** | **B0030** | **B0032** | **J61100** | **J61101** | **B0034** |
| Mean | 0.0222 | 0.0036 | 0.0057 | 0.0073 | 0.0049 |
| Std | 0.0164 | 0.0052 | 0.0087 | 0.0101 | 0.0072 |

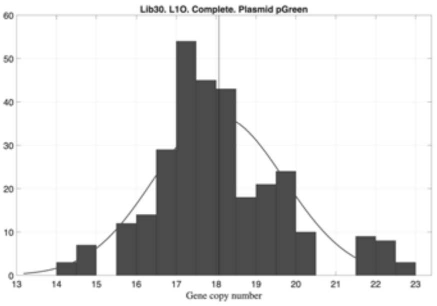

**Figure S5.2: Library $\mathcal{L}_{30}$. Histogram of the estimated gene copy number for the biopart Ori pGreen.** Obtained from the LOOCV Library Estimates (see Table S8).

these sensitivity rows across all $\mu$ values and replicate experiments yields an empirical Jacobian of the full model at the estimated parameter point. This object is mathematically equivalent to the local sensitivity matrix or Fisher Information Matrix (FIM) used in standard practical identifiability analyses Raue et al. (2009); Villaverde and Banga

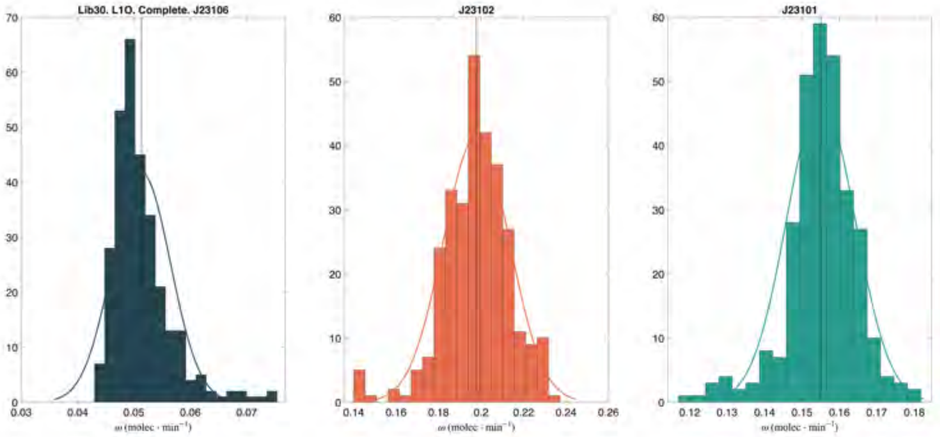

**Figure S5.3: Library $\mathcal{L}_{30}$. Histograms of the estimated promoter strengths.** Obtained from the LOOCV Library Estimates (see Table S8).

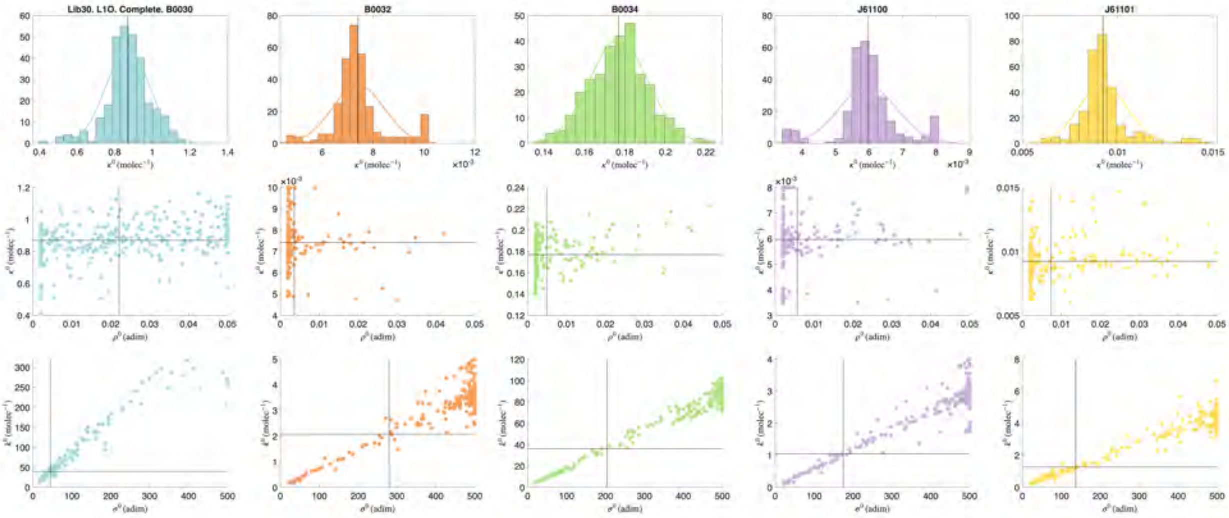

**Figure S5.4: Library $\mathcal{L}_{30}$. Histograms of the estimated RBS intrinsic initiation capacity $\kappa_A^0$ and projections along the one-dimensional $(\kappa_0^A, \rho_0^A)$ manifold.** Obtained from the LOOCV Library Estimates (see Table S8). The distributions reveal a clear unimodal structure for $\kappa_0^A$, indicating practical identifiability of the intrinsic initiation capacity across RBSs, whereas the broader dispersion observed for $\rho_0^A$ reflects the weak identifiability of the robustness factor. This behaviour is consistent with the near-colinear sensitivity structure of the pair $(\kappa_0^A, \rho_0^A)$ derived analytically in Supplementary Section S5.1. The bottom panels show the corresponding scatter plots in the original parameter space $(K_0^A, \sigma_0^A)$, where the practically invariant one-dimensional manifold becomes explicit, illustrating that multiple combinations of $(K_0^A, \sigma_0^A)$ yield indistinguishable effective translation rates. These results provide direct empirical support for fixing $\rho_0$ to a consensus value in the approximate translation model.

(2017); Gábor et al. (2017): its singular values quantify the number of independent directions in parameter space that are effectively constrained by the data.

Using the parameter estimates obtained for $\mathcal{L}_{30}$ (see Supplementary Section S5.2) and experimental data, we obtained the Jacobian ranks shown in Table S9. We carried out the practical identifiability analysis using the global mean of synthesis rate and growth rate for each TU across all the experiments, the instances means of all the well replicas of each TU for each experiment, and the whole set of all the well replicas for each TU. This a posteriori Jacobian analysis confirms experimentally the interpretation from structural analysis.

**Effect of including global means, instance means, or individual wells.** It is important to note that adding more experimental replicas (e.g. using all individual wells rather than the mean of each TU or the mean of each experimental instance) does not increase the effective rank of the Jacobian. We computed the empirical Jacobian $J(\hat{\theta})$ using three different levels of aggregation of the experimental sensitivities: (i) the global mean sensitivity for each TU and growth rate, (ii) the mean sensitivity within each independent experimental instance, and (iii) all individual wells. The three analyses yield identical ranks and effective ranks for all libraries. The only difference was the magnitude of the largest singular value, and therefore the numerical value of the tolerance $\tau = 10^{-2}\sigma_{\max}$ used to define the effective rank. This behaviour is expected: replicate wells probe the same genetic context, the same promoter–RBS combination and nearly the same host state $\mu$, so the corresponding rows of $J$ are highly colinear, increasing the

**Table S9: Jacobian rank and effective rank for the different libraries.** The exact rank is computed directly from the empirical Jacobian $J$ associated with each library, obtained using the parameter estimates obtained for the corresponding library and experimental data. The *effective rank* quantifies the intrinsic dimensionality of the information content in $J$, and is obtained from the singular value decomposition of $J$ as the number of singular values larger than $1\%$ of the largest one. This threshold separates well-informed directions in parameter space from directions that are effectively uninformative given the noise level and the structure of the sensitivity matrix.

| Library | Data replicates | Parameters | Rank | Effective rank | Tolerance |
|---|---|---|---|---|---|
| $\mathcal{L}_{30}$ | *global mean* | 14 | 14 | 9 | 1.22e3 |
| | *instance means* | 14 | 14 | 9 | 2.27e3 |
| | *all wells* | 14 | 14 | 9 | 7.99e3 |
| $\mathcal{L}_{24}$ | *global mean* | 12 | 12 | 11 | 1.24e3 |
| | *instance means* | 12 | 12 | 11 | 2.38e3 |
| | *all wells* | 12 | 12 | 11 | 8.25e3 |
| $\mathcal{L}_{6}$ | *global mean* | 2 | 2 | 2 | 1.46e2 |
| | *instance means* | 2 | 2 | 2 | 3.40e2 |
| | *all wells* | 2 | 2 | 2 | 1.08e3 |

overall scale of the Fisher Information Matrix but not introducing new independent directions in parameter space. Consequently, adding instances or wells increases the statistical robustness of parameter estimation but does not alter the identifiability structure of the model. In contrast, the effective rank increases principally when the data contain measurements at multiple growth states $\mu$ or when the TU design introduces new combinations of bioparts that excite previously unobserved sensitivity directions. This behaviour explains why using all wells is beneficial for the robustness of parameter estimation, but why the ability to identify a certain number of parameters is governed almost entirely by the combinatorial structure of the library and the diversity of $\mu$ values, rather than by the number of replicates.

**Effect of fixing known parameters.** When a biopart parameter is known *a priori* and is kept fixed during parameter estimation (e.g. the copy number of $\mathrm{pSC101}$, $N_{\mathrm{pSC101}} = 5$), the corresponding column must be removed from the Jacobian. This is not a loss of information: fixing a parameter effectively reduces the dimensionality of the estimation problem, eliminating weak or colinear directions and thereby improving the practical identifiability of the remaining parameters Raue et al. (2009); Villaverde et al. (2018); Gábor et al. (2017).

**Combinatorial diversity structures practical identifiability.** The Jacobian rank analysis reveals that combinatorial library design plays a central role for the practical identifiability of biopart parameters, not by eliminating intrinsic model degeneracies, but by constraining their expression across genetic contexts and host states. In the full libraries $\mathcal{L}_{24}$ and $\mathcal{L}_{30}$, each promoter, RBS and plasmid origin appears across multiple, partially overlapping transcriptional units and growth conditions. This combinatorial excitation generates a block-structured but full-rank Jacobian, resolving trivial multiplicative confoundings (e.g. between copy number and transcription rate) and enabling robust estimation of promoter strengths and plasmid copy numbers at the library scale.

For translation parameters, however, the situation is more subtle. Although each RBS is observed across multiple transcriptional units, the effective translation model exhibits an intrinsic near-colinearity between the composite parameters $(\kappa^0, \rho^0)$ at the single-RBS level. The combinatorial structure of $\mathcal{L}_{24}$ and $\mathcal{L}_{30}$ does not remove this invariant sensitivity direction: instead, it restricts the admissible solutions to a narrow, one-dimensional manifold in parameter space along which the effective translation rate remains nearly unchanged. Along this manifold, the marginal distributions of $\kappa^0$ are consistently unimodal across LOOCV folds for all RBSs, whereas $\rho^0$ remains weakly constrained, reflecting a genuine practical non-identifiability rather than a lack of combinatorial diversity.

In contrast, reduced sublibraries such as $\mathcal{L}_6$ collapse the Jacobian to a low-rank structure, in which only a single effective translational degree of freedom can be inferred for a new biopart, and residual contextual variability is absorbed by the remaining free parameters.

**Fixing the robustness factor $\rho^0$ to improve identifiability.** In the complete translation model, the nominal RBS parameters $(K^0, \sigma^0)$ are practically confounded, such that the data primarily constrain the reparameterized pair $(\kappa^0 = K^0/\sigma^0, \rho^0 = 1/\sigma^0)$. Indeed, when estimating the full library using LOOCV, the inferred samples collapse onto a thin, nearly one-dimensional manifold in $(\kappa^0, \rho^0)$ space, revealing a practical non-identifiability associated with an invariant direction along which the effective translation rate remains essentially unchanged. In contrast, the marginal distributions of $\kappa^0$ are consistently unimodal across LOOCV folds for all RBSs (Fig.S5.4), indicating that $\kappa_0$ constitutes the practically identifiable translational degree of freedom supported by the data, whereas $\rho_0$ remains weakly constrained unless additional assumptions are introduced.

To enable parameter transferability and improve numerical conditioning, we therefore fix the RBS robustness factor $\rho^0$ in the approximate translation model, which can be interpreted as selecting a representative gauge within this invariant subspace without altering the data-supported estimate of $\kappa^0$. Based on the full-model estimates obtained for the extended library $\mathcal{L}_{30}$, the inferred values of $\rho^0$ concentrate within a narrow physiological range across RBSs, with

B0030 yielding a representative value close to $\rho^0 \approx 0.02$ and the remaining RBSs taking equal or smaller values. We therefore adopt a consensus value $\rho^0 = 0.02$ for all approximate-model estimations.

Importantly, under the experimental conditions explored here, the elongation-related factor satisfies $f(s_n) \gtrsim 0.4$, such that $f(s_n) \gg \rho^0$ for all inferred values of $\rho^0$ (typically $\rho^0 < 0.03$). Consequently, the contribution of $\rho^0$ to the effective translation rate is negligible in the relevant growth regime, and fixing $\rho^0$ does not affect the inferred effective initiation capacity. Choosing $\rho^0 = 0.02$ further constitutes a conservative assumption, as it lies at the upper end of the empirically compatible range and therefore bounds the maximal possible influence of robustness effects on translation; only in substantially slower-growth regimes, where $f(s_n)$ becomes comparable to $\rho^0$, would variations in $\rho^0$ be expected to have a pronounced impact.

**Physiological interpretation of the estimated $\rho^0$.** The small estimated value of the robustness factor indicates that the effective translation initiation rate is highly sensitive to translational demand in the range of physiological states explored in the experiments. In our framework, the modulation of translation initiation by cellular state enters through the normalized elongation factor $f(s_n)$, which takes values in the interval $0 \leq f(s_n) \leq 1$ (see expression (21)). For the experimental growth conditions considered, the inferred values of $f(s_n)$ typically lie in the range $0.4$–$0.9$, corresponding to moderate to fast growth regimes. In this regime, $f(s_n) \gg \rho^0$, so that the sensitivity term dominates the effective RBS strength. As a consequence, for most experimentally observed growth states, the translation initiation process operates in a regime where RBSs are strongly modulated by translational demand, and differences in $\rho^0$ play a limited role in discriminating RBS behaviour. Only under low-growth conditions, where $f(s_n)$ becomes comparable to or smaller than $\rho^0$, would differences in RBS robustness be expected to have a pronounced effect on effective initiation rates. This observation explains why, under the explored experimental conditions, RBS behaviour is primarily captured by the intrinsic initiation capacity $\kappa^0 = K^0/\sigma^0$, while the individual estimation of $K^0$ and $\sigma^0$ becomes practically confounded. At the same time, it highlights that $\sigma^0$ retains a clear physiological meaning and would become experimentally identifiable in regimes of stronger resource limitation or slower growth.

*S5.4. Implications of the Jacobian rank for incremental expansion of part-characterization libraries.*

The full libraries $\mathcal{L}_{24}$ and $\mathcal{L}_{30}$ succeed because every biopart occurs in multiple, partially overlapping contexts. This resolves all multiplicative degeneracies. Small sublibraries fail because they collapse to low-rank Jacobians. Therefore, any incremental expansion strategy must guarantee new TUs that excite new sensitivity directions. The key principle is: *combinatorial diversity determines identifiability.* Some guidelines for incremental expansion are:

1. **Minimal *reference* library**. A library containing 2 ORIS, 3 promoters and 4 RBSs matches the $\mathcal{L}_{24}$ structure and guarantees a structurally full-rank sensitivity matrix for any newly added biopart.
2. **When adding a new RBS**. To characterize a single new RBS without re-estimating the full $\mathcal{L}_{30}$: (i) Combine it with all promoters in the core (*reference*) library, and (ii) include at least two plasmid origins. This ensures the new RBS is tested across sufficiently distinct host-load regimes. Then only the RBS IIC $\kappa^0$ must be estimated; the RBS RF $\rho^0$ remains poorly identifiable alone and can be fixed at the empirical consensus value 0.02. This is what happened in $\mathcal{L}_6$, where only one translational degree of freedom was identifiable.
3. **When adding a new promoter**. Combine the promoter with all existing RBS and preferably with at least two ORIs, as small single-promoter sublibraries with all translational parameters fixed lack the contextual diversity required to resolve residual translational variability, which is then absorbed by the remaining free parameter (typically the promoter strength). This is the case in library $\mathcal{L}_5$. Supplementary Fig. S5.1B shows the Jacobian restricted to the J23100 sublibrary only. Here all TUs share the same promoter and the same plasmid origin, and differ only in their RBS. As a result, the Jacobian collapses to a much poorer structure in which the only varying columns correspond to the RBS translation parameters. Because these parameters enter the model through a nearly one-dimensional sensitivity manifold, the restricted Jacobian is effectively close to rank-deficient, explaining why the promoter parameter $\omega_{\mathrm{J23100}}$ absorbs contextual variability and exhibits apparent instability under LOOCV (see Supplementary Section S9).
4. **When adding a new ORI**. The new ORI should appear in multiple promoters and multiple RBSs, so the copy number parameter can be separated from transcription and translation.
5. **When is full reestimation necessary?**. In large libraries, full re-estimation of all biopart parameters becomes neither necessary nor computationally desirable. The effective rank of the Jacobian for $\mathcal{L}_{24}$ and $\mathcal{L}_{30}$ shows that once a sufficiently diverse core set of promoters, RBSs and ORIs has been characterised, the remaining parameters form a well-identified block whose estimates can be propagated without loss of practical identifiability. Introducing a new biopart therefore requires estimating only its associated parameter(s), using a small contextual sublibrary in which the new part is combined with a selected subset of previously characterised parts that provide independent sensitivity directions. This "patchwork" design preserves the block-sparse structure of the Jacobian and guarantees identifiability of the new parameter without reintroducing confounding among the existing ones. Such incremental expansion strategies allow large-scale, continually growing part libraries to remain identifiable without ever requiring a global refit of all previously estimated parameters. Thus, accurate host-aware characterisation fundamentally relies on combinatorial designs that span several axes of genetic context.

### S6. Characterization of library $\mathcal{L}_{24}$.

This section presents the characterization of the core combinatorial library $\mathcal{L}_{24}$ using the approximate translation model in which the robustness factor is fixed to $\rho^0 = 0.02$, as justified in Supplementary Section S5.3. The aim of this analysis is twofold: first, to verify that fixing $\rho^0$ preserves the stability and consistency of the estimated biopart parameters when compared to the full-model results obtained for $\mathcal{L}_{30}$; and second, to establish a reliable reference set of promoter strengths, plasmid copy numbers, and RBS intrinsic initiation capacities that can be reused for the incremental characterization of new bioparts. Parameter estimates are reported both for full-library fits and for leave-one-out cross-validation (LOOCV), allowing direct assessment of robustness and context-independence.

**Table S10: Estimated characterization of the bioparts in library $\mathcal{L}_{24}$ using the approximated model of gene translation with fixed** $\theta_\sigma = 0.02$. For each one of the 24 leave-one-out cases, 10 estimation runs were carried out. For the full library estimation, 50 estimation runs were carried out. Units $N_A$: $\mathrm{adim.}$ The gene copy number $N_A = 5$ for the low-copy plasmid pSC101 was known beforehand from Thompson et al. (2018). Units $\omega_A$: $\mathrm{molec} \cdot \mathrm{min}^{-1}$. Units $\kappa^0$: $\mathrm{molec}^{-1}$.

| Leave-One-Out Cross-Validation Estimates | | | | | |
|---|---|---|---|---|---|
| | Plasmids | | Promoters | | |
| | $N_A$ | | $\omega_A$ | | |
| **Biopart** | **pSC101** | **pGreen** | **J23106** | **J23102** | **J23101** |
| Mean | 5 | 17.255 | 0.056 | 0.2371 | 0.2245 |
| Std | 0 | 3.438 | 0.0092 | 0.0385 | 0.0418 |
| | Ribosome Binding Sites | | | | |
| | $\kappa^0 = K_A^0/\sigma_A^0$ | | | | |
| **Biopart** | **B0030** | **B0032** | **J61100** | **J61101** | |
| Mean | 0.6765 | 0.0055 | 0.0037 | 0.0073 | |
| Std | 0.0668 | 0.0010 | 0.0007 | 0.0013 | |

| Full library Estimates | | | | | |
|---|---|---|---|---|---|
| | Plasmids | | Promoters] | | |
| | $N_A$ | | $\omega_A$ | | |
| **Biopart** | **pSC101** | **pGreen** | **J23106** | **J23102** | **J23101** |
| Mean | 5 | 16.51 | 0.055 | 0.237 | 0.220 |
| Std | 0 | 0.50 | 0.002 | 0.010 | 0.010 |
| | Ribosome Binding Sites | | | | |
| | $\kappa^0 = K_A^0/\sigma_A^0$ | | | | |
| **Biopart** | **B0030** | **B0032** | **J61100** | **J61101** | |
| Mean | 0.6955 | 0.0053 | 0.0036 | 0.0074 | |
| Std | 0.0328 | 0.0003 | 0.0002 | 0.0003 | |

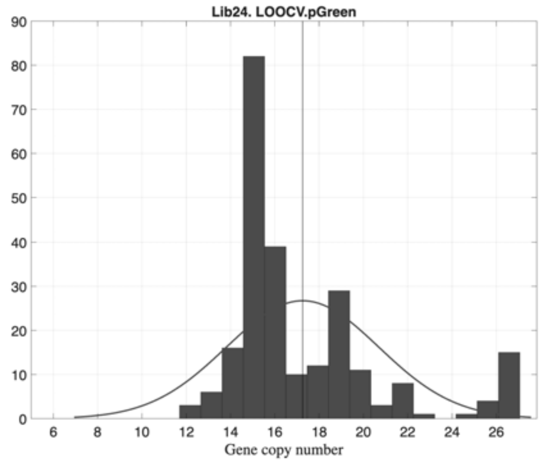

**Figure S6.1: Library $\mathcal{L}_{24}$. Histogram of the estimated gene copy number for biopart Ori pGreen.** Obtained from all the LOOCV estimations using the approximated translation model with $\theta_\sigma = 0.02$.

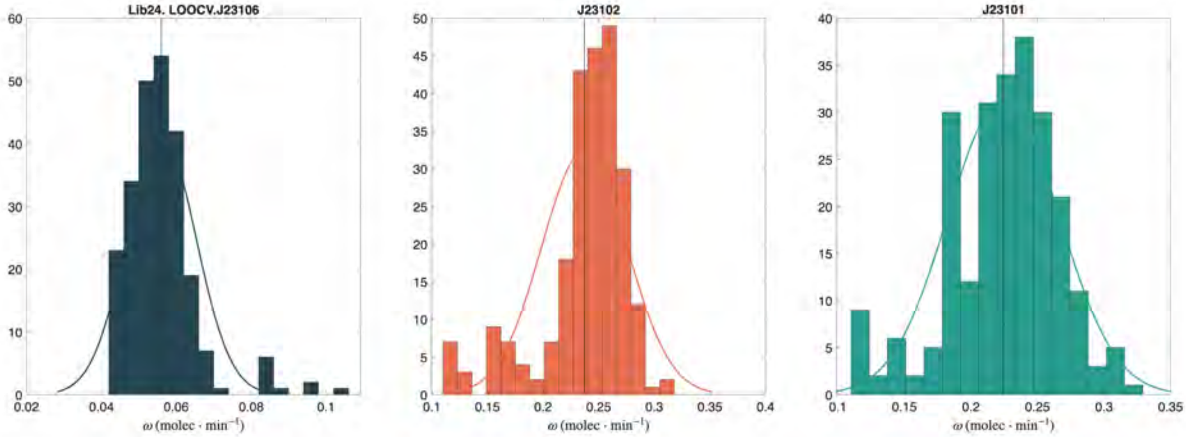

**Figure S6.2: Library $\mathcal{L}_{24}$. Histograms of the estimated promoter strengths.** Obtained from all the LOOCV estimations using the approximated translation model with $\theta_\sigma = 0.02$.

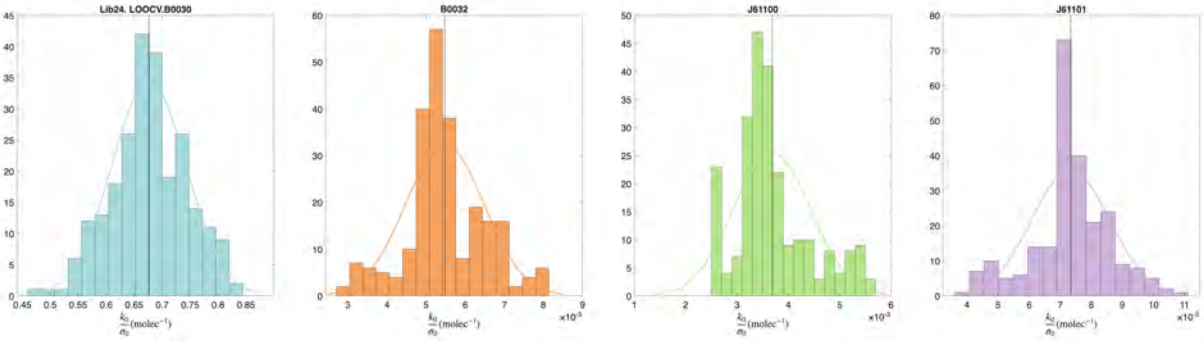

**Figure S6.3: Library $\mathcal{L}_{24}$. Histograms of the estimated RBS translational efficiency $K_A^0/\sigma_A^0$.** Obtained from all the LOOCV estimations using the approximated translation model with $\theta_\sigma = 0.02$.

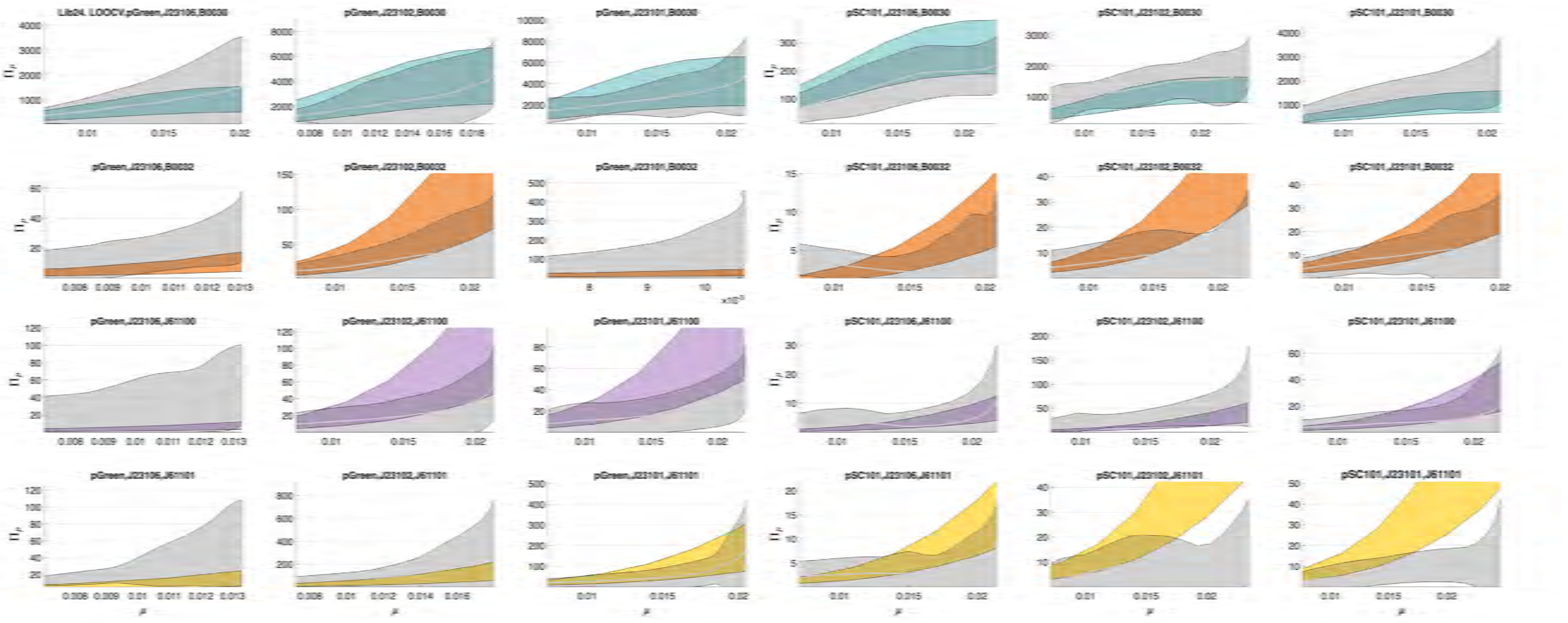

**Figure S6.4: Library $\mathcal{L}_{24}$. Experimental and predicted synthesis rate as a function of the specific growth rate using the approximated translation model with $\theta_\sigma = 0.02$.** The regions correspond to the 95% confidence interval: colored for the predicted synthesis rate and grey for the experimental data.

### S7. Characterization of RBS B0034 using the library $\mathcal{L}_6$

To characterize the RBS B0034 we employed the approximate translation model with a fixed sensitivity value of $\rho^0 = 0.02$. The remaining biopart parameters were inherited from the LOOCV characterization of the $\mathcal{L}_{24}$ library: for each promoter $\omega$ and plasmid copy number $N_A$ we used the posterior means and variances obtained previously. A hierarchical Monte–Carlo procedure was then applied to propagate this uncertainty into the estimation of the IIC parameter $\kappa^0$ for B0034. Specifically, samples were drawn from the inherited parameter distributions, and for each sampled set a full estimation of $\kappa^0_{\text{B0034}}$ was performed using the complete experimental dataset of the $\mathcal{L}_6$ library. This yielded a posterior distribution for the B0034translation parameter that reflects both experimental variability and uncertainty inherited from upstream biopart characterizations. In parallel, a leave-one-out cross-validation (LOOCV) was conducted by estimating $\kappa^0_{\text{B0034}}$ on all $6$ subsets of the $\mathcal{L}_6$ library (each omitting one TU). This analysis assesses the robustness and context-independence of the inferred parameter across the different genetic backgrounds in which B0034 appears.

**Table S11:** Estimated characterization of RBS B0034. Comparison of estimated results using the libraries $\mathcal{L}_6$ and $\mathcal{L}_{30}$ with the approximated and the complete translation model respectively. The individual LOOCV estimates gave the same results as their global mean. The results for $\mathcal{L}_{30}$ correspond to the estimation using the averages over the wells in each experiment (*instances*). Units $\kappa^0$: $\text{molec}^{-1}$.

| RBS B0034 | $\kappa^0 = K^0_A/\sigma^0_A$ | | |
|---|---|---|---|
| **Library** | $\mathcal{L}_6$ | $\mathcal{L}_6$ (LOOCV) | $\mathcal{L}_{30}$ |
| Mean | 0.1421 | 0.1433 | 0.1811 |
| Std | 0.0221 | 0.0200 | 0.0083 |

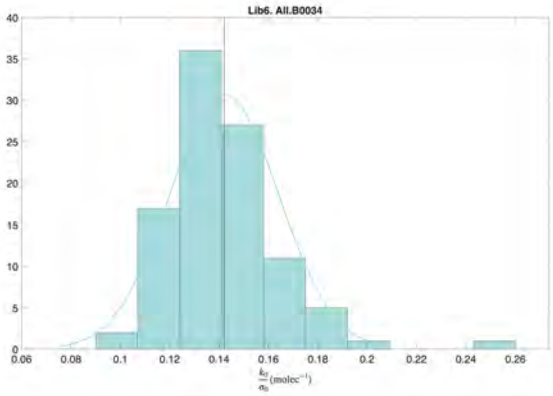

**Figure S7.1: Library $\mathcal{L}_6$. Histogram of the estimated IIC $\kappa^0_{\text{B0034}}$ for RBS B0034.** Obtained from 100 hierarchical Monte-Carlo estimations using all the TUs in library $\mathcal{L}_6$, $\rho^0 = 0.02$, and the remaining parameters inherited from the LOOCV estimations in $\mathcal{L}_{24}$.

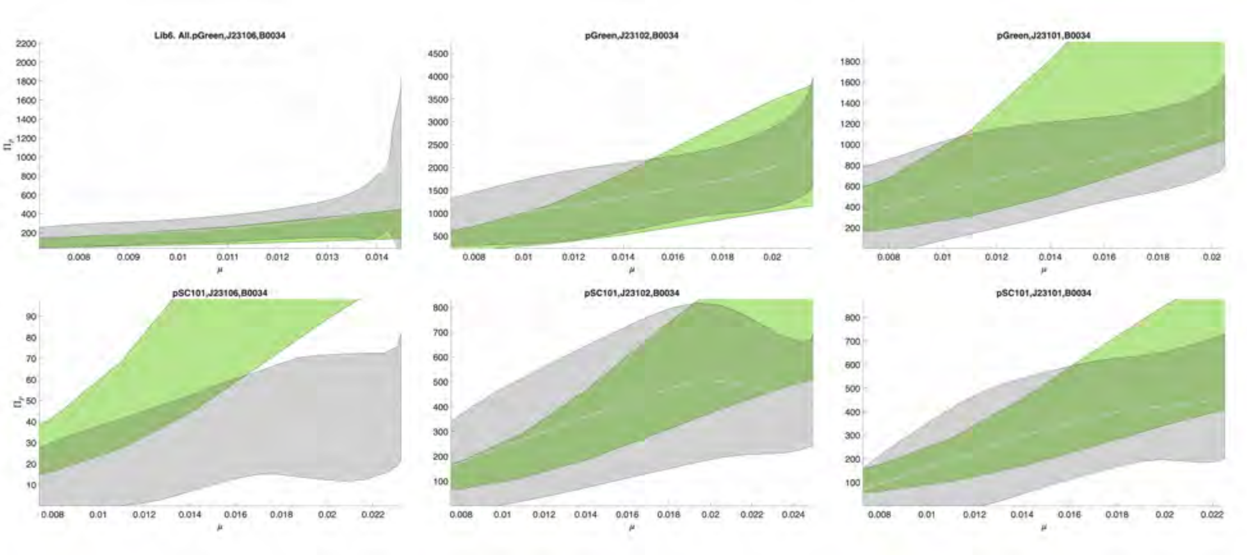

**Figure S7.2: Experimental and predicted synthesis rate as a function of $\mu$ for the TUs containing the RBS biopart B0034 in library $\mathcal{L}_6$.** The regions correspond to the 95% confidence interval: colored for the predicted synthesis rate and grey for the experimental data.

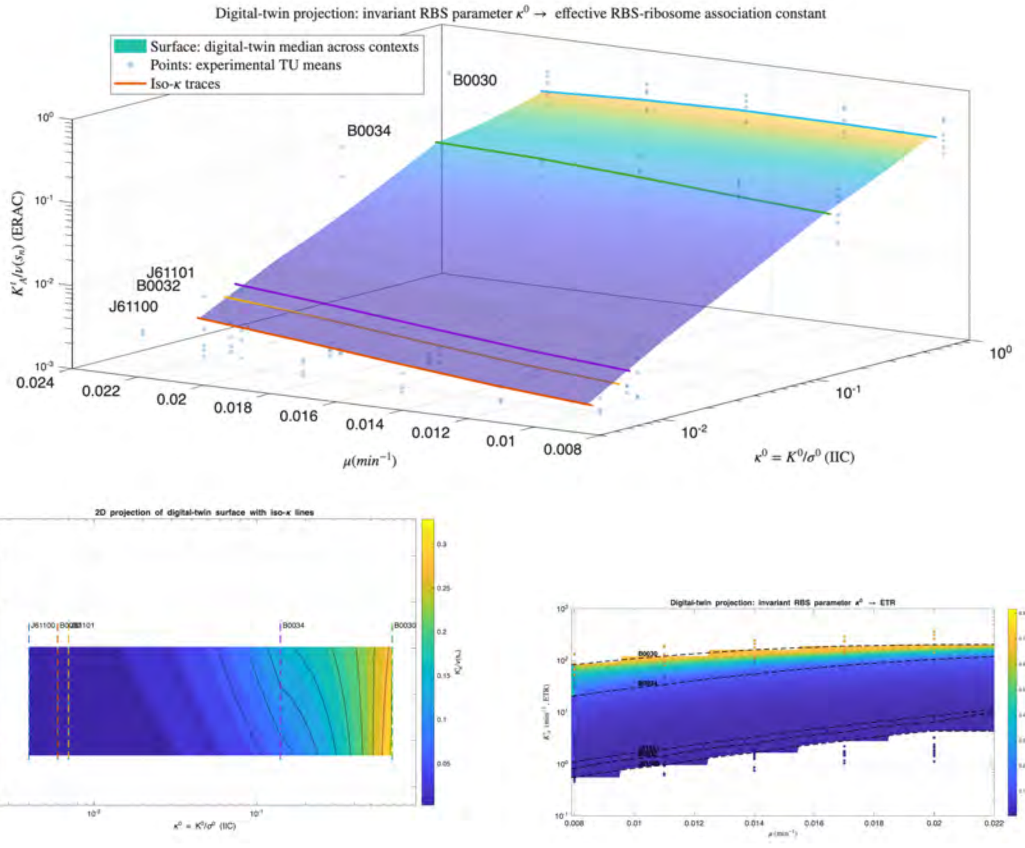

**Figure S8.1: Digital-twin projection from an invariant RBS parameter to effective translation.** (**a**) Three-dimensional representation of the same surface. Blue points correspond to experimentally measured TU means across distinct genetic contexts. Solid colored curves show iso-$\kappa^0$ traces, illustrating how context-dependent measurements project onto a single RBS-intrinsic parameter through changes in host growth state. (**b**) Two-dimensional projection of the digital-twin surface showing the effective, growth-normalized RBS–ribosome association constant $K_A^t/\nu(s_n)$ as a function of the intrinsic initiation capacity $\kappa^0 = K^0/\sigma^0$ and the specific growth rate $\mu$. Color indicates the median digital-twin prediction across genetic contexts, while black contours denote iso-surfaces of constant effective association. Vertical dashed lines indicate the estimated $\kappa^0$ values for the five RBSs used in this study. (**c**) Two-dimensional projection of the digital-twin surface showing the effective translation rate $K_A^t$ as a function of the intrinsic initiation capacity $\kappa^0 = K^0/\sigma^0$ and the specific growth rate $\mu$. Color indicates the median digital-twin prediction across genetic contexts, while black contours denote iso-surfaces of constant intrinsic initiation capacity.

### S8. RBS intrinsic initiation capacity (IIC) and effective ribosome–RBS association capacity (ERAC)

To connect sequence-level RBS properties to physiologically effective translation, we define the invariant ratio $\kappa^0 = K^0/\sigma^0$ as the intrinsic initiation capacity (IIC) of an RBS. While $\kappa^0$ is identified as a part-intrinsic parameter, the effective translation rate $K_A^t$ remains context-dependent through the host state, captured by $(\varphi, f)$ inferred by the digital twin from growth-rate measurements. Consequently, different TU contexts (copy number and promoter strength) generate a spread of $K_A^t$ values at fixed $\kappa^0$, which the model explains via load-dependent changes in $\varphi$ rather than by allowing $\kappa^0$ to vary with context. For comparison with initiation-centric metrics in the literature, we report the elongation-normalized effective ribosome–RBS association capacity (ERAC), $K_A^t/\nu(s_n)$.

The effective translation rates $K_A^t$ inferred from the digital-twin projections span values in the range $1$ and $200\ \mathrm{min}^{-1}$ (approximately $0.01$–$3\ \mathrm{s}^{-1}$), well within the range reported for effective translation rates and initiation-limited protein production in *E. coli* Bonde et al. (2016); Salis et al. (2009); Bremer and Dennis (2008). These values correspond to values of elongation-normalized effective ribosome–RBS association capacity in the range $10^{-3} - 10^{-1}\,\mathrm{molec}^{-1}$ (see figure S8.1) that correspond to weak-moderate associations and non saturated regime of ribosomes. These values are consistent with known elongation speeds, polysome loading, and ribosomal availability, and reflect physiologically constrained translation rather than idealized initiation rates. Importantly, in our framework $K_A^t$ emerges as a context-dependent projection of the invariant RBS parameter $\kappa^0$, enabling direct comparison with experimental estimates of effective translation rates while preserving a mechanistic separation between intrinsic biopart properties and host-state modulation.

Together, these representations show that apparent variability in translation efficiency across contexts arises primarily from host physiological state, while $\kappa^0$ remains invariant across genetic backgrounds.

### S9. Characterization of promoter J23100 using the library $\mathcal{L}_5$

The $\mathcal{L}_5$ sublibrary contains only five transcriptional units with promoter J23100, all in the same plasmid background (pGreen) and differing only in their RBS. Because this design provides a limited diversity of genetic contexts, the associated empirical Jacobian is intrinsically low-rank (see Section S5.1), and the leave-one-out (LOOCV) estimates of $\omega_{23100}$ exhibit confounding between promoter strength and RBS-dependent translational context. As a result, the LOOCV estimates do not provide an independent validation of $\omega_{23100}$ in this sublibrary. Notice the higher values (0.422,0.142) of the individual LOOCV estimates (see Table S12) correspond to the estimated promoter strength $\omega$ when the RBSs B0030 and B0034 are left out. These act as anchors for the estimation.

This behaviour does *not* reflect a limitation of the host-aware modelling framework; rather, it is the expected consequence of the reduced contextual diversity in $\mathcal{L}_5$. In full combinatorial libraries (e.g. $\mathcal{L}_{24}$ and $\mathcal{L}_{30}$), where each RBS appears with multiple promoters and plasmid origins, this confounding is resolved and the transcriptional strengths are consistently and robustly estimated. Here, the LOOCV variability in $\mathcal{L}_5$ is therefore interpreted as a result of (i) insufficient combinatorial excitation and (ii) promoter–RBS interactions at the DNA or $5^{'}$-UTR level that are not captured by the effective biophysical model. These interactions are visible in the experimental $\Pi(\mu)$ curves (see Figures S9.2 and S2.2), where certain TUs ( B0032 and J61100) deviate from their expected behaviour considering the constructs that share the same Ori and RBSs.

Despite the low-rank nature of $\mathcal{L}_5$, full-library estimates from $\mathcal{L}_5$ yield $\omega_{23100} \approx 0.16$ (see Table S12), a value lower than the original Anderson ranking but consistent with previous reports documenting context-dependent weakening of J23100 Mutalik et al. (2013); Brophy and Voigt (2014). This value should therefore be considered the reliable estimate of the promoter strength, whereas the LOOCV variability within $\mathcal{L}_5$ primarily reflects the limited identifiability of this small sublibrary rather than uncertainty in $\omega_{23100}$ itself.

**Table S12: Estimated characterization of promoter J23100.** Compararison of estimated results using the library $\mathcal{L}_5$. The individual LOOCV estimates are shown in parenthesis. Units $\omega_A$: $\mathrm{molec} \cdot \mathrm{min}^{-1}$.

| Global estimation | | LOOCV estimation | |
|---|---|---|---|
| | $\omega_A$ | | $\omega_A$ |
| **Library $\mathcal{L}_5$** | | **Library $\mathcal{L}_5$** | |
| Mean | 0.161 | Mean | 0.226 (0.422,0.140,0.266,0.142,0.160) |
| Std | 0.039 | Std | 0.122 (0.093,0,029,0.066,0.029,0.033) |

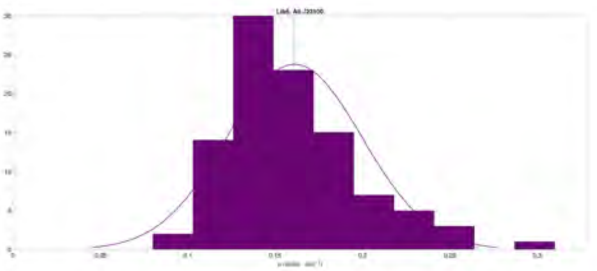

**Figure S9.1: Library $\mathcal{L}_5$. Histogram of the estimated $\omega_A$ for promoter** J23100. Obtained from 100 hierarchical Monte-Carlo estimations using all the TUs in library $\mathcal{L}_5$. The remaining bioparts parameters were inherited from the LOOCV estimations obtained for libraries $\mathcal{L}_{24}$ and library $\mathcal{L}_6$

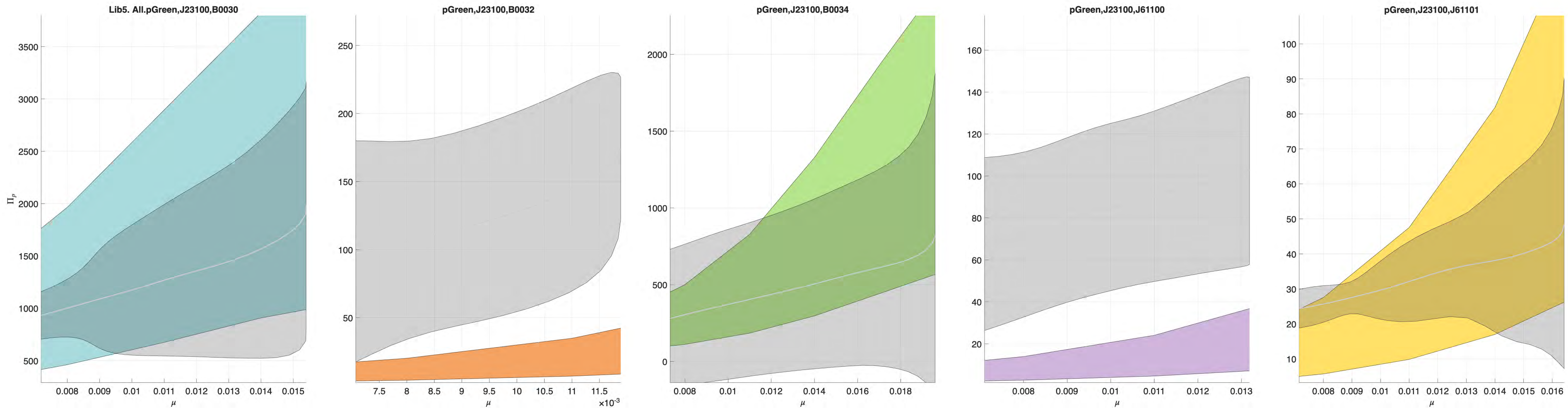

**Figure S9.2: Experimental and predicted synthesis rate as a function of $\mu$ for the TUs containing the promoter biopart** J23100 **in library $\mathcal{L}_5$.** The regions correspond to the 95% confidence interval: colored for the predicted synthesis rate and grey for the experimental data.

### S10. Summary of estimated parameters and their distributions.

This section provides a consolidated summary of the intrinsic biopart parameters estimated throughout the study, combining the results obtained from the core library $\mathcal{L}_{24}$, the incremental characterization of RBS B0034 using $\mathcal{L}_6$, and the promoter J23100 characterization using $\mathcal{L}_5$. The reported distributions correspond to leave-one-out cross-validation estimates and therefore reflect both experimental variability and practical identifiability limitations discussed in previous sections. Presenting all parameters together facilitates direct comparison across bioparts and highlights the relative separation between promoters, RBS intrinsic initiation capacities, and plasmid copy numbers achieved by the host-aware, combinatorial characterization framework.

**Table S13: Complete set of estimated parameters.** The results shown consist of the estimations obtained using LOOCV for libraries $\mathcal{L}_{24}$, $\mathcal{L}_6$(B0034), and $\mathcal{L}_5$(J23100) as described in Supplementary Sections S6, S7 and S9. Units $N_A$: $\mathrm{adim.}$ The gene copy number $N_A = 5$ for the low-copy plasmid pSC101 was known beforehand from Thompson et al. (2018). Units $\omega_A$: $\mathrm{molec} \cdot \mathrm{min}^{-1}$. Units $\kappa^0$: $\mathrm{molec}^{-1}$.

| | Plasmids $N_A$ | Promoters $\omega_A$ | | | |
|---|---|---|---|---|---|
| **Biopart** | **pGreen** | **J23106** | **J23102** | **J23101** | **J23100** |
| Mean | 17.255 | 0.056 | 0.2371 | 0.2245 | 0.161 |
| Std | 3.438 | 0.0092 | 0.0385 | 0.0418 | 0.039 |
| | RBS $\kappa_A^0$ | | | | |
| **Biopart** | **B0030** | **B0032** | **J61100** | **J61101** | **B0034** |
| Mean | 0.6765 | 0.0055 | 0.0037 | 0.0073 | 0.1433 |
| Std | 0.0668 | 0.0010 | 0.0007 | 0.0013 | 0.0200 |

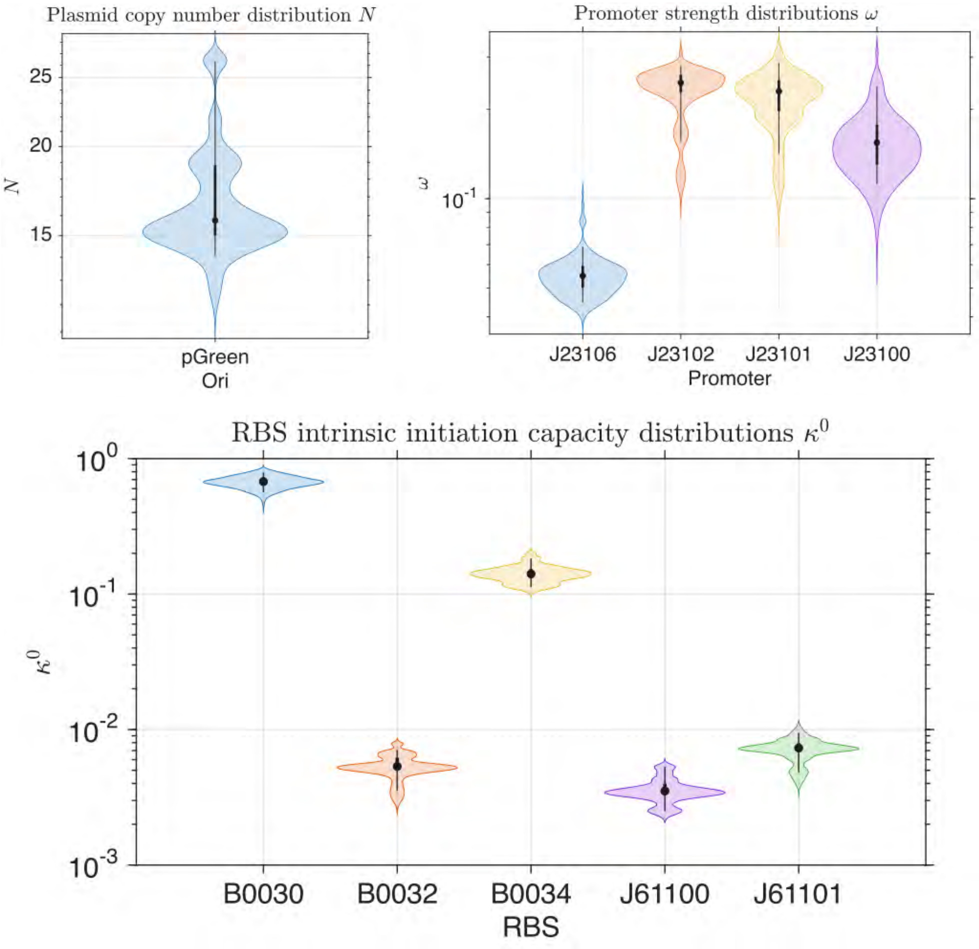

**Figure S10.1: Complete set of estimated parameters.** Violin plots for the distributions of the estimations obtained using LOOCV for libraries $\mathcal{L}_{24}$, $\mathcal{L}_6$(B0034), and $\mathcal{L}_5$(J23100) .
